# Comprehensive profiling of anaesthetised brain dynamics across phylogeny

**DOI:** 10.1101/2025.03.22.644729

**Authors:** Andrea I. Luppi, Lynn Uhrig, Jordy Tasserie, Golia Shafiei, Antoine Légaré, Kanako Muta, Junichi Hata, Hideyuki Okano, Daniel Golkowski, Andreas Ranft, Rudiger Ilg, Denis Jordan, Silvia Gini, Zhen-Qi Liu, Yohan Yee, Camilo M. Signorelli, Rodrigo Cofre, Alain Destexhe, David K. Menon, Emmanuel A. Stamatakis, Patrick Desrosiers, Paul De Koninck, Christopher W. Connor, Alessandro Gozzi, Ben D. Fulcher, Bechir Jarraya, Bratislav Misic

**Affiliations:** Division of Anaesthesia, Department of Clinical Neurosciences, and St John’s College, University of Cambridge, Cambridge, UK; Montréal Neurological Institute, McGill University, Montréal, QC, Canada; Centre for Eudaimonia and Human Flourishing, Department of Psychiatry, University of Oxford, Oxford, UK; Cognitive Neuroimaging Unit, CEA, INSERM, Université Paris-Saclay, NeuroSpin Center, Gif-sur-Yvette, France; Department of Anesthesiology and Critical Care, Necker Hospital, Université de Paris Cité, Paris, France; Center for Brain Circuit Therapeutics, Brigham and Women’s Hospital, Harvard Medical School, Boston, MA, USA; Department of Psychiatry, Perelman School of Medicine, University of Pennsylvania, Philadelphia, PA, USA; Centre de recherche CERVO, Québec QC, Canada; Graduate School of Human Health Sciences, Tokyo Metropolitan University, Arakawa, Tokyo, Japan; Laboratory for Marmoset Neural Architecture, Center for Brain Science, RIKEN, Wako, Saitama Japan; Department of Physiology, Keio University School of Medicine, Shinjuku, Tokyo, Japan; Department of Neurology, Klinikum rechts der Isar, Technical University Munich, Munich, Germany; Department of Anesthesiology and Intensive Care, Technical University of Munich, Munich, Germany.; Asklepios Clinic, Department of Neurology, Bad Tolz, Germany; Department of Anaesthesiology and Intensive Care Medicine, Klinikum rechts der Isar, Technical University Munich, Munich, Germany; University of Applied Sciences and Arts Northwestern Switzerland, Muttenz, Switzerland; Center for Neuroscience and Cognitive Systems, Istituto Italiano di Tecnologia, Rovereto, Italy; Centre for Mind/Brain Sciences, University of Trento, Italy; Center for Philosophy of Artificial Intelligence, University of Copenhagen, Copenhagen, Denmark; Paris-Saclay University, CNRS, Paris-Saclay Institute for Neuroscience (NeuroPSI), Saclay, France; Division of Anaesthesia, University of Cambridge, Cambridge, UK; Department of Clinical Neurosciences, University of Cambridge, Cambridge, UK; Université Laval, Québec QC, Canada; Department of Anesthesiology, Perioperative and Pain Medicine, Brigham and Women’s Hospital, Boston, MA, USA; Department of Biomedical Engineering, Physiology and Biophysics, Boston University, Boston, Massachusetts; School of Physics, The University of Sydney, Sydney, Australia; Department of Neurology, Foch Hospital, Suresnes, France

## Abstract

Intrinsic dynamics of neuronal circuits shape information processing. Combining neuroimaging with causal perturbation offers the opportunity to understand how local dynamics mediate the link between neurobiology and functional repertoire. We compile a unique dataset of multi-scale neural activity during wakefulness and anaesthetic-induced suppression of information processing encompassing human, macaque, marmoset, mouse, zebrafish and nematode. Applying massive feature extraction, we comprehensively characterise local neural dynamics across >6,000 time-series features. Using dynamics as a common space for cross-species comparison reveals a conserved dynamical profile of anaesthesia across species, characterised by shorter intrinsic timescales of neural activity and dampened interregional synchrony. This dynamical regime is experimentally reversed *in vivo* by deep-brain stimulation of the macaque centromedian thalamus, restoring behavioural responsiveness. Spatially, this conserved dynamical phenotype covaries with conserved transcriptional profiles of excitatory and inhibitory neurotransmission across human, macaque, marmoset and mouse cortex. Biophysical modelling provides a mechanistic link between the macroscale dynamical phenotype of anaesthesia, and microscale effects of key molecular targets on the timescales of synaptic excitation and inhibition. Altogether, comprehensive dynamical phenotyping reveals a shared neural endpoint of anaesthesia: across species and scales, anaesthetics induce spatio-temporal isolation of local neural activity.

## INTRODUCTION

From invertebrates to primates, an essential function of the nervous system is to enable the organism to respond to an ever-changing environment. Indeed, brain activity is in constant flux—reflected in rich dynamics of neuronal activity. Local signaling events then propagate via axonal projections, manifesting as coherent and patterned neural activity over the cortex as revealed by multiple neuroimaging modalities, ranging from single-neuron calcium imaging, to electrophysiology, to functional MRI. As a result, these neural dynamics span multiple spatial and temporal scales [1–5]. Mapping how dynamics support brain function is a key goal in the neurosciences.

A prominent paradigm for understanding neural dynamics and their function is to manipulate them via general anaesthesia. Anaesthetic agents modulate neuronal signaling, altering local and global dynamics, reversibly suppressing the brain’s ability to process information and respond to the external environment. Although different species have evolved unique ways to respond to their specific environments, the behavioural effects of anaesthetics (namely, suppression of behaviour and inability to interact with the environment) are highly conserved across species, from primates to nematode worms [6–8], hinting at shared and fundamental underlying mechanisms. Neural changes observed at the macroscale can then be related to downstream effects on cognition and behaviour, and to upstream cellular, molecular, and synaptic mechanisms at the microscale. Thus, systematically and reversibly perturbing brain function with anaesthesia while recording neural activity provides a unique opportunity to understand how the dynamics of local neural activity mediate the link between anatomy, chemoarchitecture, and the organism’s functional repertoire. Indeed, numerous imaging studies have reported evidence of changes in neural activity that accompany transitions between wakefulness and anaesthesia [6, 8–23].

However, the current picture of how anaesthetics influence neural dynamics remains fragmented and incomplete. On one hand, some effects may be specific to a particular species, or a particular anaesthetic agent. On the other hand, despite some notable exceptions (e.g., [24–28]) most studies tend to focus on a single species, a single anaesthetic, and specific hand-picked features of neural activity—such as spectral power, amplitude, or temporal entropy. Yet, each time-series can be characterised by thousands of different properties: from statistics of the distribution such as mean, variance, and outliers, to periodicity and autocorrelation at different lags, stationarity, and measures of signal complexity and selfsimilarity [29–31]. Therefore, the currently prevalent approach of focusing on a handful of pre-selected properties leaves open the risk of missing key aspects of neural dynamics, potentially leaving large gaps in our understanding.

To go beyond these limitations of the current literature and triangulate on a potential ‘final common endpoint’ of anaesthesia [6, 32, 33], here we systematically map how anaesthetics perturb the entire dynamic profile of the brain across several species and across several pharmacological agents. We first compile a unique multiscale dataset spanning single-neuron recordings in invertebrates (*c.elegans*), non-mammalian vertebrates (larval zebrafish), as well as functional MRI in murine (mouse) and primate species (marmoset, macaque, human) undergoing imaging during loss and recovery of responsiveness induced by multiple volatile and intravenous anaesthetics, as well as reawakening from anaesthesia induced by deep-brain stimulation (DBS) of the macaque central thalamus (Fig. 1a). Altogether, the species included here encompass key model organisms for neuroscience, spanning over 700 million years of evolution [34] (Fig. S1). We then apply state-of-the-art methods for massive feature extraction [29, 30], to systematically characterise the activity of every cortical region/neuron in this broad and diverse dataset, in terms of more than 6 000 univariate features from the vast modern literature on timeseries analysis [31, 35–40] (Fig. 1b). The result is an exhaustive characterisation of how anaesthesia reshapes the dynamics of neural activity: across dynamical features, across pharmacological agents, across species, and across scales, from single neurons to entire brain regions.

**Figure 1.**
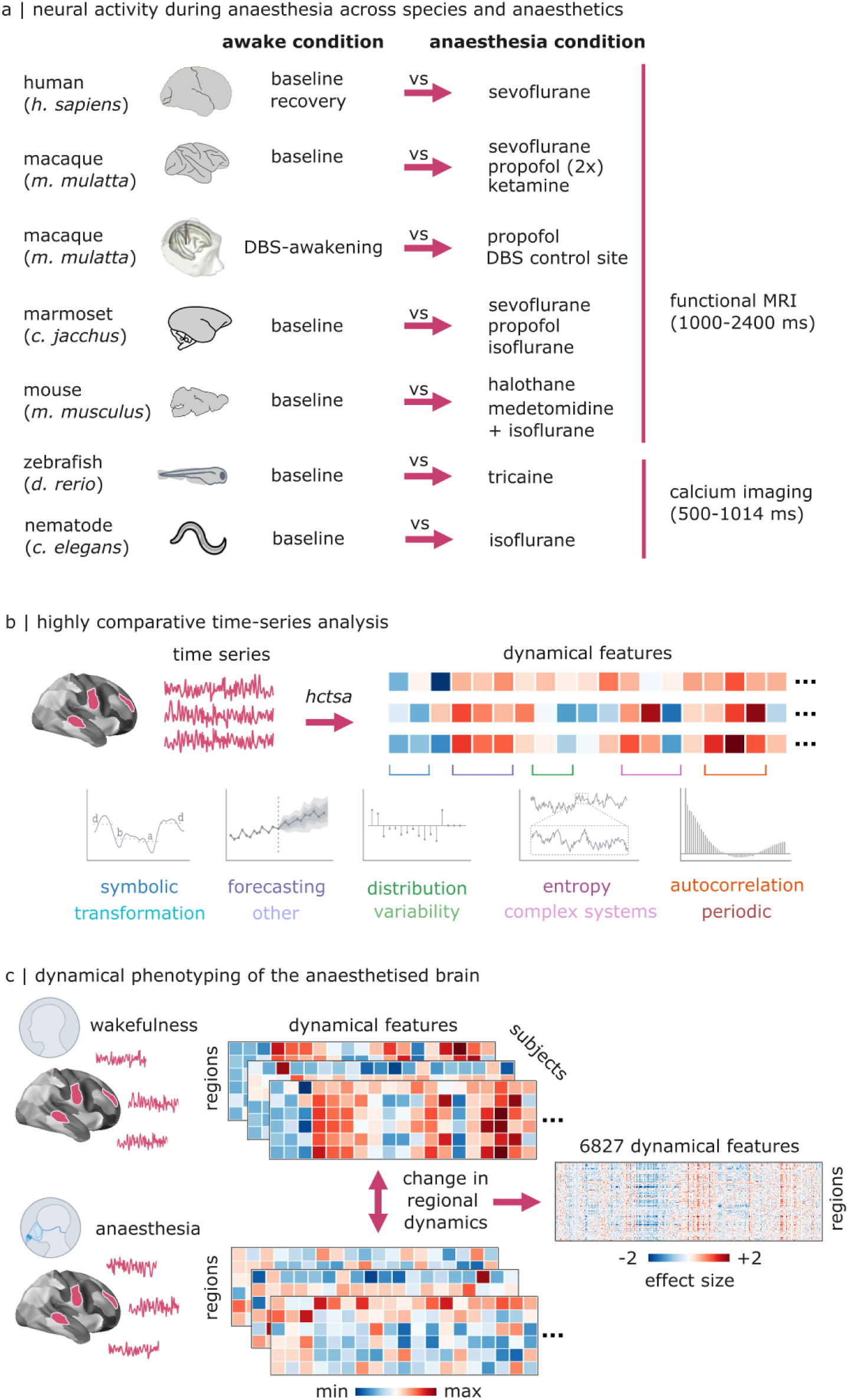
Systematic phenotyping of brain dynamics under anaesthesia. (**a**) Overview of datasets and contrasts included in the present study. Human: awake vs vol 3% sevoflurane, and recovery vs vol 3% sevoflurane [42]. Macaque (two datasets): awake vs sevoflurane; awake vs propofol; awake vs ketamine [15]; and awake vs propofol (no DBS); central thalamus (CT) DBS versus propofol (no DBS); and CT DBS vs ventral thalamus (VT) DBS [43] Marmoset: awake vs sevoflurane; awake vs propofol; and awake vs isoflurane [43]. Mouse: awake vs halothane; and awake vs medetomidine-isoflurane [14]. Zebrafish: awake and tricaine anaesthesia. Nematode: 0% isoflurane vs 4% isoflurane [44]. For the mammalian species, time-series were acquired using functional MRI. For the nematode and zebrafish, time-series were acquired from GCamP calcium imaging. (**b**) Using *highly comparative time-series analysis* (hctsa [29]), we extract >6 000 dynamical features to characterise each time-series of regional/neuronal brain activity. (**c**) Analytic strategy: for each brain region, we compare dynamical features during wakefulness and during anaesthesia. We obtain a regions-by-features matrix of effect sizes for each contrast, which can be compared across datasets. See Table S1 for full details on sample sizes, imaging techniques, and acquisition parameters description.

This comprehensive characterisation enables us to pursue one key hypothesis: that across different species, different anaesthetics exert their common effect of suppressing behaviour by eliciting convergent changes in local neural dynamics. Although an early hypothesis that all anaesthetic agents share precisely the same molecular mechanism (altering neuronal lipid membrane) was ultimately rejected [41], the question remains of how a common functional outcome is achieved by pharmacologically diverse substances, across species with widely different neural systems [6, 32, 33]. Here we hypothesise that despite their divergent molecular targets, anaesthetics induce convergent changes in local neural dynamics. Focusing on effects that are consistently shared across multiple anaesthetics and species enables us to exclude any physiological or methodological confounds that are specific to any one drug, or species, or imaging modality, and instead triangulate on what they all have in common: breakdown of responsiveness to the environment. In sum, our strategy is to combine comprehensive data-driven screening of thousands of dynamical features, with a stringent criterion for consistency across species and anaesthetics demanded by our driving hypothesis (Fig. 1c).

To foreshadow our results, comprehensive and systematic phenotyping of dynamical features under anaesthesia reveals a dynamical regime of neural activity that is conserved across species, scales, and drugs. Under anaesthesia, local neural activity fails to propagate in space and time. Deep-brain stimulation of the central thalamus can reverse this effect, demonstrating bidirectional control over brain dynamics. Finally, biophysical modelling provides a mechanistic link between the macroscale dynamical phenotype of anaesthesia, and microscale effects of key molecular targets on the timescales of synaptic excitation and inhibition.

## RESULTS

Here we perform comprehensive phenotyping of the changes in local neural dynamics induced by different anaesthetic agents, across over 6 000 dynamical features of univariate time-series from across the scientific literature [29–31]. We compile and systematically phenotype multiple datasets, each including at least one awake and one anaesthetised condition (Fig. 1a). Please see *Methods* for specific details regarding acquisition and processing of each dataset; an overview of the key parameters is provided in Table S1. Specifically, we include: a dataset of resting-state functional MRI data obtained from N=15 healthy humans (*Homo sapiens*) who were scanned at baseline and during deep anaesthesia with the inhalational anaesthetic sevoflurane, as well as during spontaneous recovery of consciousness [42]; a dataset of N=5 macaque monkeys (*Macaca mulatta*) scanned several times during wakefulness and during anaesthesia with sevoflurane, propofol, or ketamine [15]; a dataset of N=4 marmoset monkeys (*Callithrix jacchus*) scanned during wakefulness and during anaesthesia with isoflurane, sevoflurane, or propofol [45]; a dataset of N=43 mice (*Mus musculus*) scanned either during wakefulness (N=10) or during anaesthesia with halothane (N=19) or combined medetomidine-isoflurane (N=14) [14]. To broaden our investigation beyond mammalian species, we also include a new dataset of N=7 larval zebrafish (*Danio rerio*) with neuronal calcium imaging acquired during wakefulness or in the presence of the anaesthetic tricaine, having a comparable temporal resolution to our fMRI recordings (1014ms). Since the effects of anaesthesia are shared even with invertebrates [6–8], we also include a dataset of N=10 nematode worms (*Caenorhabditis elegans*) with calcium imaging acquired with or without isoflurane anaesthesia, having a comparable temporal resolution to our fMRI recordings (500ms) [46]. For all the above datasets, we contrast every wakeful condition against every anaesthetised condition. Finally, we also include an additional macaque dataset, comprising N=2 macaque monkeys scanned several times with fMRI during wakefulness, during propofol anaesthesia, and during anaesthesia combined with deep-brain stimulation of different thalamic nuclei: centromedian thalamus (CT), which induced awakening despite continuous anaesthetic infusion; and ventral-lateral thalamus (VT) which did not induce awakening [43]. For this dataset, we contrast baseline wakefulness against propofol anaesthesia. We also contrast awakening induced by CT DBS during anaesthesia (i.e., propofol is present but the animal is awake), against anaesthesia without DBS, and against anaesthesia with VT DBS (which does not induce awakening). Inclusion of this unique dataset enables us to dissociate the mere presence of a high dose of propofol from its effect on wakefulness.

Starting from the preprocessed time-series of each brain region’s fMRI activity (respectively, each neuron/neuron group’s calcium imaging activity in the nematode and zebrafish data) we use the highly comparative time-series analysis (hctsa) toolbox [29, 30] to extract its full dynamical profile across the most comprehensive available set of scientific time-series features (Fig. 1b). The hctsa features include methods from across the interdisciplinary scientific literature on timeseries analysis, including biology but also physics, engineering, and economics. Features range from basic statistics of the distribution of time-points (e.g. mean, variance), outliers, periodicity, and stationarity, to linear and nonlinear autocorrelation at different lags, predictability of future activity from its past using models of various complexity, motifs of the discretised time-series and their prevalence, and many more [29, 30]. Each time-series feature takes a univariate time-series as input, and returns a single, real-valued summary statistic. Filtering for features that could not be computed for all regions/neurons yielded 6 827 features for further analysis. For each wakefulness-anaesthesia contrast (e.g., baseline vs isoflurane anaesthesia in marmoset; recovery versus sevoflurane in human; deep-brain stimulation vs no stimulation in macaque), we perform a statistical test to compare the value of each feature during wakefulness and during anaesthesia. This is performed separately for each region (resp., neuron). For each contrast, this procedure returns a matrix of size brain regions/neurons × 6 827 features (Fig. 1c). Note that the features’ raw values can vary widely across features, spanning several orders of magnitude and hindering comparison. However, here we are not interested in raw feature values, but rather in how they change under anaesthesia. Rather than quantifying this anaesthetic-induced difference in terms of the original feature units (e.g., mean difference), which would incur in the same problem, we can instead quantify it by its effect size. Being expressed in units of standard deviation, effect sizes are comparable across features and also across regions, species, and datasets. We measure effect size using Hedge’s *g*, which is interpreted in the same way as Cohen’s *d*, but more appropriate for small sample sizes [47]. Since the same features are used across all species, this procedure ensures that our datasets can be meaningfully compared: same features and same units.

### Systematic phenotyping of local neural dynamics under anaesthesia

To test the hypothesis that diverse anaesthetics exert their evolutionarily conserved effect of suppressing behaviour by eliciting similar changes in local neural dynamics, we do not pre-specify some dynamical features of putative interest. Rather, we systematically and comprehensively assess thousands of features from across the scientific literature on time-series analysis. Fig. 2 shows the matrix of regions × dynamical features obtained by averaging across all contrasts and datasets belonging to the same species. Each regions-by-features matrix shows the overall effect size when contrasting the awake condition versus the anaesthesia condition, such that positive (red) values indicate an increase during anaesthesia. Within each species, the majority of features appear to be affected by anaesthesia in the same way across regions/neurons (i.e. most columns are either predominantly red or predominantly blue, indicating consistent increase/decrease). Nonetheless, this consistency does not imply uniformity: differences among regions can also be observed. Fig. S3 shows the same data after z-scoring each column and reordering the rows within each species, to further highlight patterns of regional heterogeneity. As expected, some differences among species are evident: some features show inconsistent patterns of increase and decrease in different species; and the mouse, zebrafish, and nematode data display overall greater sensitivity (evidenced by overall greater effect sizes). Although inter-species differences and differences in the types and doses of anaesthetics used may play a role, some of the observed differences may potentially be explained by the higher spatial and temporal resolution of their respective acquisitions. Namely, the mouse data has the fastest fMRI sampling and longest time-series, while the nematode and zebrafish data use calcium imaging to measure activity of individual neurons at even greater spatial and temporal resolution (Table S1).

**Figure 2.**
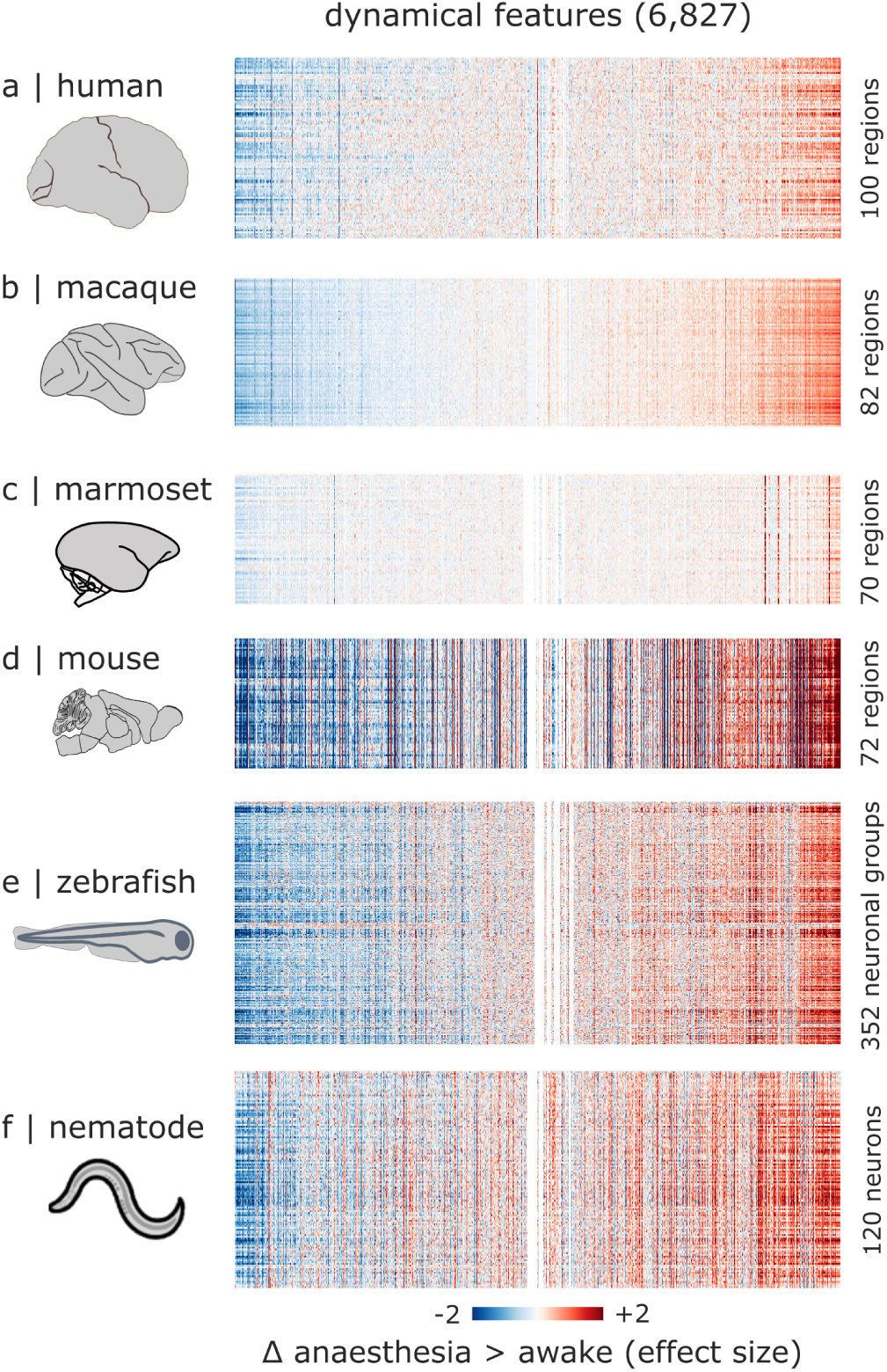
Anaesthetic-induced changes in neural dynamics across species. (**a**) Human: mean effect sizes across awake vs vol 3% sevoflurane, and recovery vs vol 3% sevoflurane. (**b**) Macaque: mean effect sizes across awake vs sevoflurane; awake vs propofol; awake vs ketamine (for the Multi-anaesthesia dataset); awake vs propofol (no DBS); reawakening induced by CT DBS versus propofol anaesthesia without DBS; and reawakening induced by CT DBS vs VT DBS (which does not reawaken the animal), both during propofol anaesthesia (for the DBS dataset). (**c**) Marmoset: mean effect sizes across awake vs sevoflurane; awake vs propofol; and awake vs isoflurane. (**d**) Mouse: mean effect sizes across awake vs halothane; and awake vs medetomidine-isoflurane. (**e**) Zebrafish: effect size for awake vs tricaine. (**f**) Nematode: effect size for awake vs isoflurane. For all species, positive effect sizes indicate anaesthesia > awake. The order of features (columns) is the same in each species, sorted to highlight common patterns. For visualisation purposes, the color range is capped at [-2, 2]. Fig. S3 shows the same data after column-wise z-scoring and reordering the rows, to highlight patterns of regional heterogeneity.

Importantly, despite differences in acquisition and spatial resolution, processing strategies, model organism and anaesthetic agent and dose, we observe prominently aligned vertical bands of the same colour (indicating consistent change) among datasets, suggesting that diverse anaesthetics elicit similar increases and decreases in local dynamics across species (Fig. 2). Notably, a very similar profile of changes in brain dynamics is observed in the macaque, when instead of comparing anaesthesia against baseline wakefulness, anaesthesia is compared against reawakening induced by deep-brain stimulation of the centromedian thalamus (CT). Likewise, a similar profile of changes is observed when comparing DBS of the ventral thalamus, which does not reawaken the animal, against CT stimulation (Fig. S2). In other words, when deep-brain stimulation of the thalamus successfully reverses the behavioural effects of anaesthesia, it also reverses the effects of anaesthesia on the profile of local neural dynamics from fMRI.

We next home in on the phylogenetically-conserved dynamic profile associated with anaesthesia. We proceed as follows. For each contrast (comparison between an awake condition and an anaesthetised condition, using a particular anaesthetic in a particular species) we first average the effect size of each feature across all regions/neurons, to obtain a single overall value representing how each feature is affected by that anaesthetic in that species. This generates one summary effect size per feature, per contrast (Fig. 3a). Next, we compare the contrasts to identify time-series features that are consistently altered across species and across anaesthetics.

**Figure 3.**
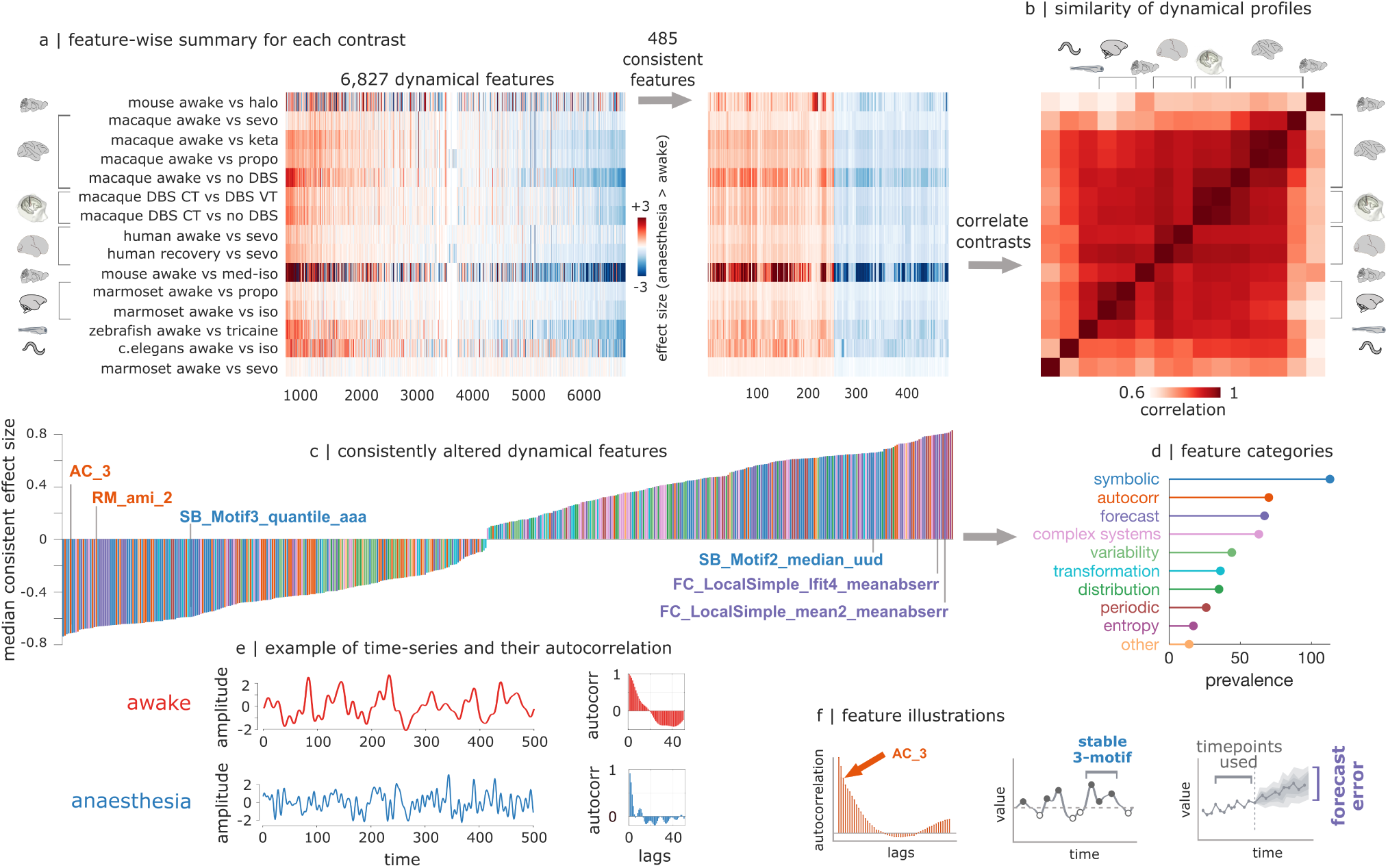
Consistent effects of anaesthesia on neural dynamics. (**a**) Summary of feature-wise dynamical changes for each contrast and each species. For each contrast, effect sizes are averaged across brain regions/neurons to obtain a single value for each feature, representing its overall change across the brain. To identify dynamical features that are consistently altered across species and across anaesthetics, features are filtered to retain only those whose direction of effect is the same in each species (all positive or all negative). The outcome of this procedure is a set of 485 dynamical features that are consistently altered across regions/neurons, across anaesthetics, and across species. For visualisation purposes, the color range is capped at [-3, +3]. (**b**) The dynamical signatures of anaesthesia are positively correlated across species and across anaesthetics (*mean* = 0.86). This value is significantly greater than the level of correlation that would be expected by chance purely based on the fact that features have the same sign, as quantified by a null distribution generated by shuffling the features within each contrast while preserving their sign (p < 0.0001; Fig. S7). (**c**) The median effect sizes (Hedge’s *g*) are shown for each of the 485 consistent features, color-coded according to membership of 10 broad categories of dynamics (feature types; Table S2). Exemplary features are highlighted. (**d**) Prevalence of 10 broad categories of features (see *Methods*) among the consistent ones, revealing a predominance of features pertaining to autocorrelation, forecasting error, and symbolic motifs. (**e**) Examples of time-series from an awake (red) and an anaesthetised (blue) macaque, showing faster decay of autocorrelation under anaesthesia. (**f**) Illustrations of representative features from the most prevalent categories: autocorrelation; a time-series discretised into symbols above and below a cut-off value to produce motifs (symbol sequences); and error in forecasting the future of the time-series from a model trained on past values.

Note that this analysis includes the contrast between CTDBS and no-DBS in the macaque, as well as between CTDBS and VT-DBS, i.e. when the animal is re-awakened despite the presence of propofol in the bloodstream— thereby ensuring that we only retain features that track the anaesthetic’s effect on behavioural responsiveness, not its mere presence. Specifically, we filter the features to retain only those whose overall direction of change (increase or decrease) is the same across all awake-anaesthesia contrasts, in every species considered here (i.e., all positive or all negative). Note that this is a stringent criterion: any feature would be excluded from this set if its direction of change were inconsistent in even just one contrast of one dataset. With 15 contrasts, the probability of a feature exhibiting such a consistent change purely by chance is 1*/*(2^14^) or 0.00006; hence we should expect a single consistent feature due to chance for every 16,384 (more than twice as many features as we include). Instead, we find 485 time-series features that are consistently altered across regions/neurons, across anaesthetics, and across species (Fig. 3a). These dynamical signatures of anaesthesia are highly correlated across different anaesthetics and different species (Fig. 3b).

Broadly, we find that on one hand, anaesthesia reduces the value of features related to linear and nonlinear temporal autocorrelation in the brain (Fig. 3c,d). For example, some of the features exhibiting large reductions are *AC_3* and *RM_AMI_2*, which index the autocorrelation at lag 3 and automutual information at lag 2 of the timeseries, respectively. On the other hand, anaesthesia increases the error of forecasting the future of a time-series based on its past (Fig. 3c,d). For example, *FC_LocalSimple_lfit4_meanabserr* quantifies the mean absolute error of a linear fit from the past 4 timepoints, and *FC_LocalSimple_mean2_meanabserr* reflects the mean absolute error of forecasting based on the mean of the last 2 time-points. Fig. 3e illustrates how these features capture changes in the underlying dynamics of neural time-series, focusing on autocorrelation as a feature that is straightforward to compute and readily interpretable.

The vastness and diversity of the scientific literature on time-series analysis comes with a challenge: not all features are equally straightforward to interpret. Some features may come from more obscure pockets of the literature, or reflect more convoluted properties or transformations on the data. We used a range of approaches to address this issue. First, we used domain knowledge across the author group to identify meaningful and interpretable properties among the consistent ones (pointing to autocorrelation and forecasting error). We also stratified dynamical features into 10 broad categories, based on associated keywords in hctsa [29, 30] and further inspection (see *Methods* and Table S2). This is not a formal taxonomy: some features may plausibly belong to more than one category, and some categories are only loosely defined. Nonetheless, stratifying features can be helpful to narrow down on what the consistent features returned by our comprehensive screening have in common. In particular, we find that among the consistent features, in addition to forecasting and autocorrelation categories there is also a high prevalence of the symbolic category (Fig. 3d). Guided by this observation, we find that anaesthesia reduces the prevalence of ‘stable’ or ‘uniform’ motifs, whereby the same symbol occurs multiple times in a row (for example, a motif *SB_motif3_quantile_aaa* whereby the time-series value is in the top of three quantiles three times in a row) and instead it increases the prevalence of ‘unstable’ or ‘mixed’ motifs (e.g. *SB_motif2_median_uud*, whereby the time-series mixes values higher and lower than a median split), hinting at overall destabilisation of local dynamics. Illustrations of the three most prevalent categories of dynamical features are provided in Fig. 3f. Additional example time-series for each contrast and species, including the effects of DBS, are provided in Fig. S4, and statistical comparisons for each contrast are provided for *RM_AMI_2* in Fig. S5. Altogether, we find that anaesthesia induces a breakdown in the relationship between past and future of neural activity and its stability, which is reversed upon reawakening induced by by thalamic deep-brain stimulation in the macaque.

Temporal autocorrelation of spontaneous neural activity reflects how long neural information persists in a local brain area to influence its future behaviour: the “temporal receptive window” over which inputs coming into that brain region are integrated [2, 4, 48–52]. Our datadriven analysis suggests that this fundamental organisational property of the brain is compromised under anaesthesia. We therefore follow up this data-driven analysis in a targeted manner, by explicitly quantifying the intrinsic timescale at which each region operates. We define intrinsic timescale as in [53], as the product between repetition time (TR) and the sum of autocorrelation function values up to the point where autocorrelation drops to zero, such that the future of the time-series becomes independent of its past (Fig. 4a). A larger value of this measure indicates a longer intrinsic timescale [53]. For each species (human, macaque, marmoset, mouse, zebrafish, and nematode), we compare the distribution of regional intrinsic timescales between wakefulness and anaesthesia. Within every single species we find that on average across regions, intrinsic neural timescales are significantly reduced under anaesthesia (Fig. 4b-g; results for each individual contrast are provided in Fig. S6). This effect is neither uniform across regions, nor random. Rather, anaesthetic-induced reduction of intrinsic timescales is proportional to each region’s baseline level of intrinsic timescale, such that regions/neurons with longer intrinsic timescales see the most severe reductions, indicated by significant negative correlations across regions between baseline level and anaesthetic-induced change (Fig. 4b-g). The anaesthetic-induced shortening of intrinsic local timescales is reversed by deep-brain stimulation of the centromedian thalamus in macaques, tracking behavioural responsiveness: restoration is not observed upon ineffective stimulation of the control site in ventral thalamus (Fig. S8). Altogether, at the temporal resolution afforded by fMRI and GCamP calcium imaging, we find that anaesthesia consistently induces a compression of the brain’s intrinsic timescales, leading to reduced local persistence of information, and shorter temporal windows for integration.

**Figure 4.**
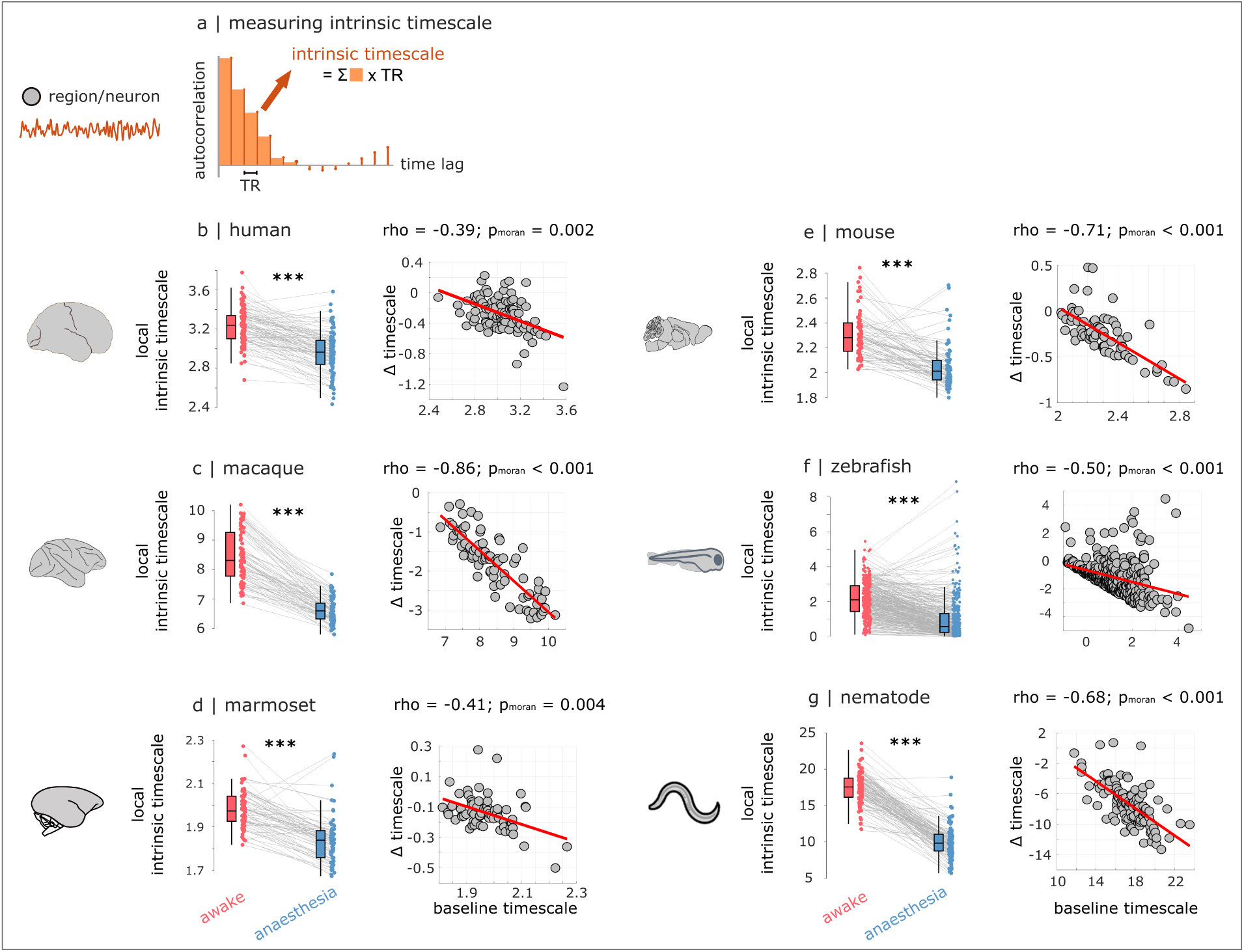
Anaesthesia reduces intrinsic neural timescales across species. (**a**) Quantifying intrinsic neural timescales from spontaneous activity. For each region/neuron, the autocorrelation function is obtained by correlating the original time-series with a time-lagged version of itself, for increasing lags. Intrinsic neural timescale is computed by summing the values of the autocorrelation function (each multiplied by the TR) up to the point where autocorrelation drops to zero. A larger value of this measure therefore indicates a longer intrinsic timescale [53]. (**b-g**) For each species (human, macaque including the DBS contrasts, marmoset, mouse, zebrafish, and nematode), we obtain the mean intrinsic timescale of each region across wakefulness (red), and across anaesthesia conditions (blue). Within each species, we compare these distributions of regional intrinsic timescales. Each data-point represents one region/neuron. Box-plots: center line, median; box limits, upper and lower quartiles; whiskers, 1.5*×* interquartile range. ***, *p <* 0.001, from non-parametric paired-samples t-test. We also show the correlation between anaestheticinduced change in regional intrinsic timescale, and each region’s intrinsic neural timescale while awake. Significance of correlations is assessed against null models generated with Moran spectral randomisation (see *Methods*).

### From local dynamics to global networks

Up to this point we have focused on local dynamics. However, individual neurons and regions are embedded in a larger synaptic network. In a complex network such as the brain, changes in regional dynamics may both shape and be shaped by changes in inter-regional functional interactions. For instance, if two populations begin to operate at faster or slower timescales, what effect does this have on their capacity to spontaneously synchronize with each other? To investigate the relationship between local dynamics and network-wide communication, we estimate two measures of inter-regional dependence (Fig. 5a). First, we compute the temporal synchrony of co-fluctuations in neural activity, operationalised as the magnitude of functional connectivity (zero-lag Pearson’s correlation between BOLD/calcium imaging time-series; Fig. 5a). Second, we compute the magnitude of pairwise similarity between regional dynamical features generated by our large set of time-series features [31], measuring the dynamical profile similarity (DPS) of two neuronal populations [31](Fig. 5c). The two measurements allow us to identify distributed regions that potentially display common local dynamics— suggesting common functional roles—without necessarily displaying time-locked activity. We find that across species and anaesthetics, anaesthesia desynchronises the interactions between different regions (Fig. 5b). At the same time, we find that anaesthesia does not only change the dynamical properties of individual regions: it also weakens the relationships between the dynamical profiles of different regions, significantly reducing their similarity (Fig. 5d). Additionally, in the awake brain, regions (or neurons) with similar dynamics—suggesting similar computational properties—tend to display time-locked activity, suggesting that they also perform the same function (Fig. 5e). To provide an analogy: co-workers (regions) with similar job descriptions (dynamical profiles) may be likely to work similar hours (i.e., be synchronised). However, in the anaesthetised brain, regions with similar dynamics are less likely to exhibit synchronous fluctuations, and lose their coordination (Fig. 5f; see also Fig. S9 for statistical comparisons for each contrast and dataset). Notably, just like anaesthesia’s effects on regional dynamics, these effects of anaesthesia on interregional interactions are all reversed upon reawakening induced by deep-brain stimulation of the centromedian thalamus, whether contrasted with no stimulation, or with inefficacious stimulation (at same intensity) of the control site in ventral thalamus (Fig. 5b,d,f). Altogether, anaesthesia changes both local neural dynamics, and inter-regional interactions.

**Figure 5.**
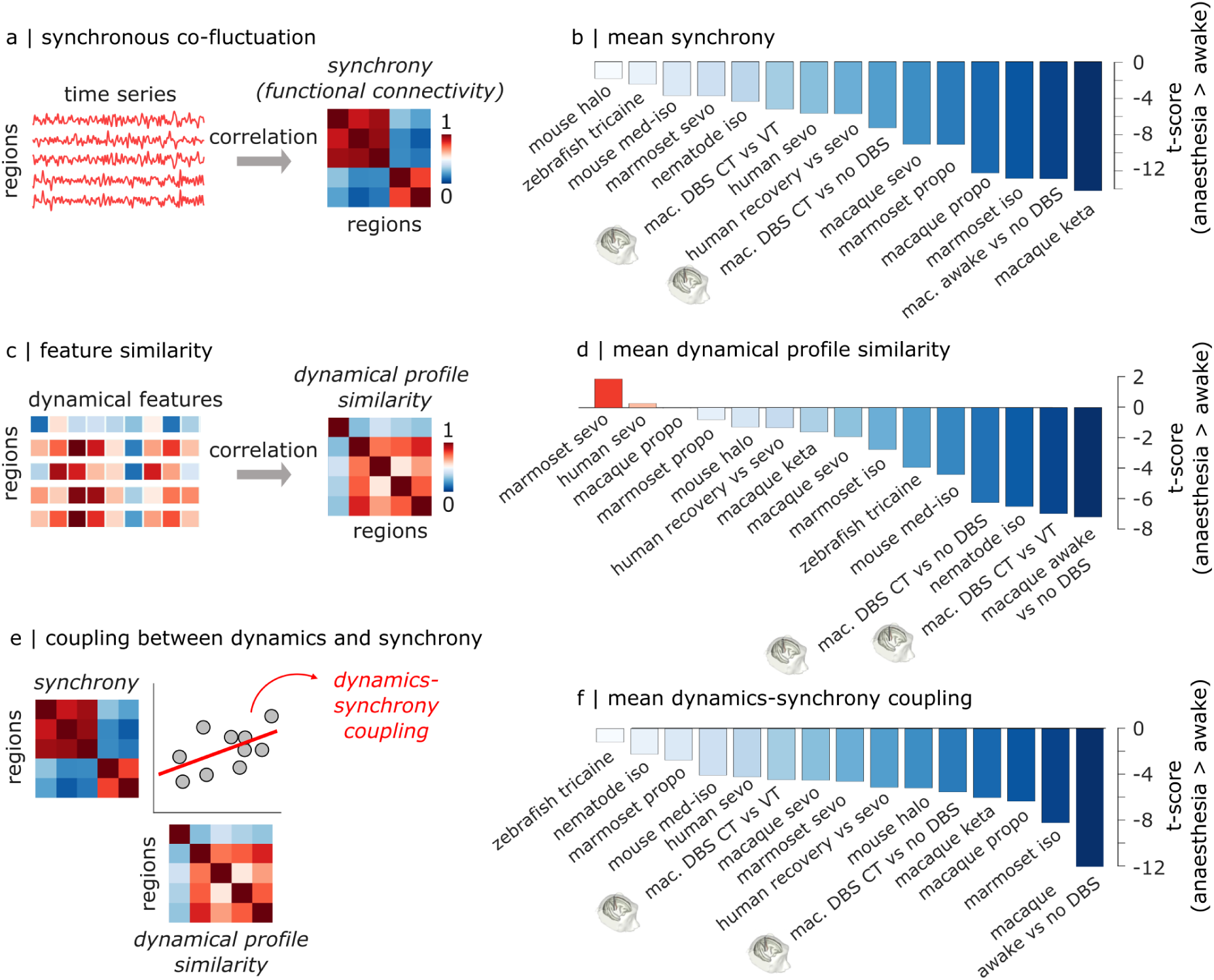
Anaesthesia desynchronises the brain and decouples synchrony from local dynamics. (**a**) Synchrony is measured as the magnitude of correlation between two regions’ time-series over time, reflecting co-fluctuation. (**b**) Across species and anaesthetics, anaesthesia significantly reduces the mean magnitude of synchronous co-fluctuations of neural activity. (**c**) Dynamical profile similarity (DPS) is measured as the magnitude of correlation between the dynamical features of two regions’ time-series. For both synchrony and DPS, the result is a region-by-region matrix of pairwise similarities. (**d**) Anaesthesia significantly reduces the mean similarity between regions’ dynamical profiles. (**e**) Coupling between synchrony and dynamics is obtained by correlating the matrix of inter-regional synchrony and the matrix of inter-regional dynamical profile similarity. It quantifies the extent to which regions characterised by similar time-series features also exhibit similar temporal co-fluctuations. (**f**) Across species and anaesthetics, anaesthesia reduces the coupling between synchrony and dynamics.

### Mapping local neural dynamics to gene expression across species

So far, we observed that diverse anaesthetics induce consistent changes in local neural dynamics across species. Since anaesthetics can perturb local biophysics by acting on various molecular targets and ion channels, for example altering postsynaptic excitability in response to neurotransmitter release [41, 54, 55], we next ask whether these phylogenetically conserved changes in local dynamics might be underpinned by patterns of gene expression that are similarly conserved across species. To address this question, we use species-specific databases of cortical transcriptomics for human (microarray [56]), macaque (stereo-seq [57]), mouse (in situ hybridization [58]) and marmoset (in situ hybridization [59, 60]) (Fig. 6a). For each species, we map spatial patterns of cortical expression for a list of 23 orthologous brain-related genes that are available across all four species. This list is a subset at the intersection of the previously published lists of human-macaque [61] and human-mouse [62] orthologous brain-related genes, and genes for which quantitative data are also available throughout the cortex for the marmoset. It comprises genes pertaining to neurotransmitter and neuropeptide receptors, including key glutamate and GABA receptors (AMPA, NMDA, GABA*_A_*) as well as the dopaminergic, serotonergic, noradrenergic, and cholinergic systems; myelin; and interneuron cell-type markers (parvalbumin, somatostatin, calbindin, vasoactive intestinal polypeptide) (see Table S4 for the list of gene names).

**Figure 6.**
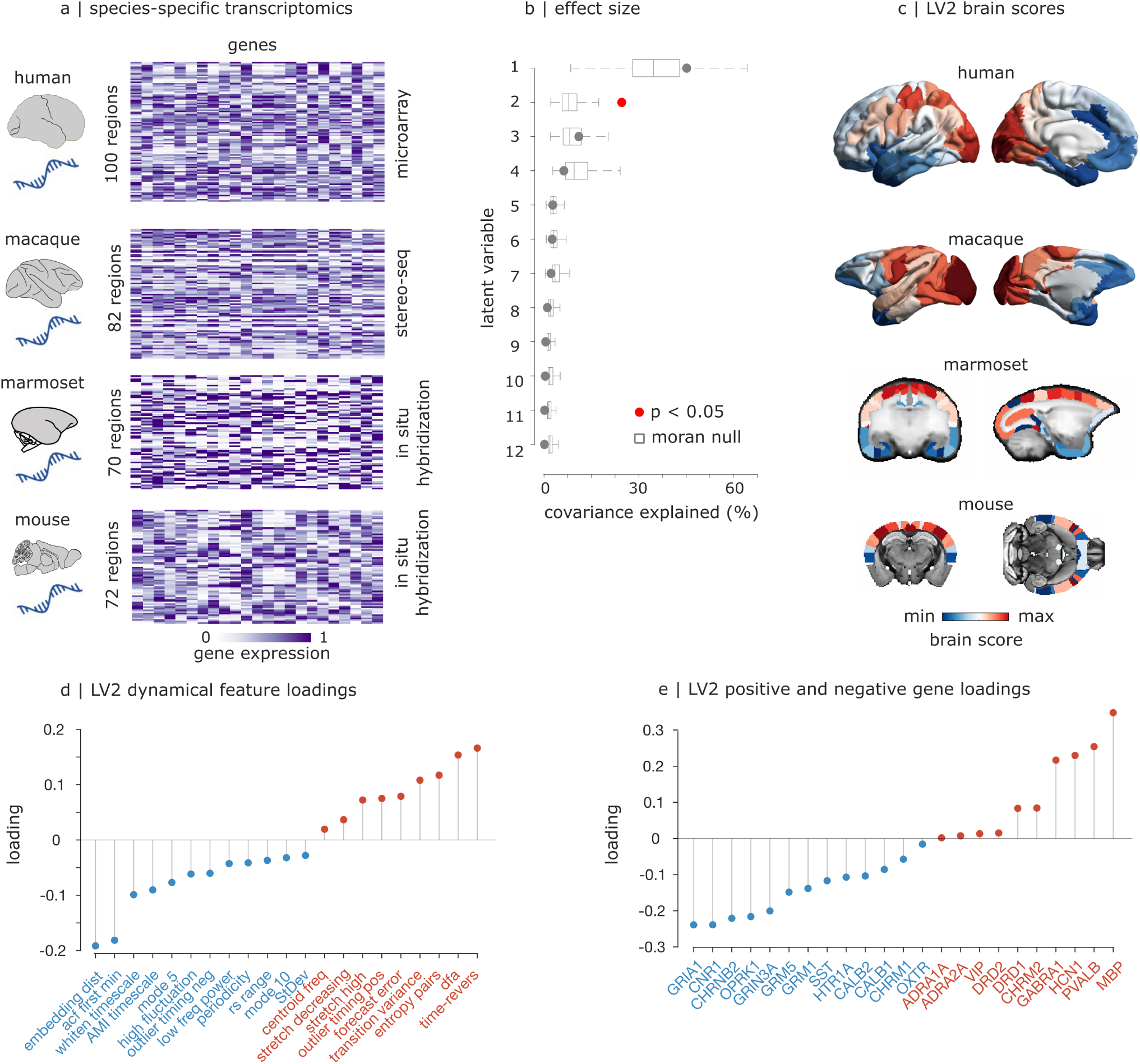
Anaesthetic-induced changes in neural dynamics map onto phylogenetically conserved dimensions of cortical gene expression. (**a**) We obtain species-specific cortical expression of key brain-related genes from microarray (human), stereo-seq (macaque) and in-situ hybridization (marmoset and mouse). Expression of each gene is sigmoid-normalised to lie in the interval between 0 and 1. (**b**) We find a statistically significant latent dimension of multivariate association between gene expression and anaesthetic-induced feature change (LV2). The first latent dimension is not significant beyond the effect of spatial autocorrelation.(**c**) Representation of the significant LV2 on the cortex of human, macaque, marmoset, and mouse, delineating a conserved anterior-ventral to posterior-dorsal gradient.(**d**) Loading of each representative dynamical feature onto the significant dimension of multivariate association with gene expression. (**e**) Loading of each gene onto the significant dimension of multivariate association with dynamical features. Red indicates positive loading, blue indicates negative loading.

We then use partial least squares correlation (PLS) to integrate cortical transcriptomics with dynamical features (each concatenated across species, after z-scoring within species). PLS is a data-driven technique that identifies multivariate patterns of maximum covariance between datasets—in this case, regional gene expression and regional changes in dynamical features, each z-scored within species and then concatenated across species (see *Methods*) [63, 64]. We assess significance against a distribution of null models generated using Moran spectral randomisation (see *Methods*), whereby brain maps of each species are randomised but their spatial autocorrelation is preserved, embodying the null hypothesis that local dynamics and gene expression are spatially correlated with each other purely because of inherent spatial autocorrelation [65, 66]. As a form of unbiased feature selection to aid interpretation, instead of the full set of consistent dynamical features we use a reduced set of representative time-series features (known as the CAnonical Time-series CHaracteristics or catch22 [67]). Lubba et al. [67] identified this set of minimally-redundant features in a data-driven manner, as providing similar performance to the full hctsa set across a large number of natural and human-made dynamical systems, while reducing feature dimensionality to simplify computation and interpretation. Since catch22 includes features pertaining to autocorrelation, forecasting, and temporal frequency, capturing many dynamical properties that we found to be consistently perturbed by anaesthesia, for our main PLS analysis we use the catch22 features to provide more interpretable results (Fig. 6; see *Methods* and Table S3) for the list of 21 included features and their explanations). Crucially, the same latent variables are recovered when using the full set of consistent dynamical features (see below; Fig. S14 and Fig. S15).

PLS reveals a statistically significant latent variable (linear weighted combination of the original variables) relating anaesthetic-induced changes in local dynamics, and cortical patterns of gene expression, across species. This latent variable (LV2) explains significantly more covariance than explained by spatial autocorrelation alone (25%; by contrast, LV1 is not significant, after taking into account spatial autocorrelation). This significant latent variable is associated with consistent cortical patterns in each species, which can be broadly characterised as anterior-ventral to posterior-dorsal (Fig. 6c). At one end, LV2 relates measures of time-reversibility and forecasting error with the expression of a key marker gene for inhibitory interneurons (*PVALB*), and the *α*1 subunit of the inhibitory GABA*_A_* receptor, target of propofol and many volatile anaesthetics [41, 55, 68], as well as the pace-maker channel *HCN1*, a shared target of both propofol and ketamine [20, 41, 55]; myelin-related gene *MBP*; and the inhibitory M2 muscarinic cholinergic receptor *CHRM2* (Fig. 6d,e). At the other end, LV2 relates changes in measures of autocorrelation with NMDA excitatory receptors *GRIN3A*, which are inhibited by anaesthetics including ketamine [55, 69, 70]; as well as other glutamatergic excitatory receptors, both ionotropic (*GRIA1* AMPA receptor) and metabotropic (*GRM5*, *GRM1*); excitatory nicotinic cholinergic receptors (*CHRNB2*), and cannabinoid and opioid neuromodulatory receptors (*CNR1*, *OPRK1*) (Fig. 6d,e). Importantly, highly consistent latent variables for the association between anaesthetic-induced changes in local neural dynamics and cortical transcriptomics are recovered, when the analysis is repeated using all dynamical features that are consistently perturbed by anaesthesia across species, instead of the reduced set from catch22 (Fig. S14). We observe the same significant LV2, with the same anterior-ventral to posterior-dorsal cortical gradient across species, alongside correlated gene loadings (Fig. S15; note that sign of PLS latent variables is arbitrary). Altogether, we find that conserved anaesthetic-induced changes in local dynamics are subtended by a transcriptomic gradient that is likewise conserved (at least across rodent and primate species), encompassing genes pertaining to regulation of excitation, inhibition, and neuronal rhythms—possibly indicating shared neurobiological circuitry on which anaesthesia operates.

### Computational modelling of anaesthetised neural dynamics from microscale to macroscale

Finally, having identified a consistent dynamical signature of anaesthesia across species and across anaesthetics, we seek to better understand its mechanistic origin through computational modelling. Computational models provide a way to link specific biophysical parameters to their functional consequences, by simulating how brain activity changes upon changing the parameters [71–76]. In this vein, recent work developed a computational approach to evaluate how molecular mechanisms impact large-scale brain activity, and used it to model anaesthesia [76]. The modelling framework consists of a whole-brain model of biophysically grounded mean-fields, which integrate cellular and molecular properties (membrane conductances and AMPA, NMDA, and GABA*_A_* synaptic receptors), thereby linking microscale properties to their macroscale consequences. At the microscale, the model simulates anaesthesia as biologically realistic increase in the time constant of inhibitory synaptic decay, which is empirically observed with GABA-ergic anaesthetics (such as propofol, sevoflurane, isoflurane, halothane) that act as positive allosteric modulators of the GABA*_A_* receptor [41, 68, 77, 78], potentiating GABA-mediated inhibition by prolonging the duration of inhibitory postsynaptic potentials [76]. Mean-field reductions of the microscale model are used to simulate regional BOLD activity. Once connected according to empirical structural connectivity of the human brain, the resulting whole-brain model links biophysically meaningful synaptic-level mechanisms to macroscale dynamics [76].

In other words, molecular mechanisms are propagated consistently from the single-cell synaptic scale to mesoscale mean-field regional dynamics and finally to macroscale structure-constrained whole-brain activity. Crucially, here we do not perform any additional fitting or tuning of the model by Sacha et al. [76], instead using the published parameters for awake and anaesthetised dynamics. In other words, we use the model to ask whether anaesthetic-induced prolongation of inhibitory postsynaptic potentials, a phenomenon experimentally observed at the microscale, is mechanistically sufficient to reproduce the dynamical phenotype of anaesthesia that we observed at the macroscale. To test this hypothesis, we apply comprehensive dynamical phenotyping to evaluate the model’s capacity to reproduce the empirical changes in local dynamics induced by anaesthesia. Remarkably, we find that 95% of the features that are consistent across all empirical contrasts (461/485; significantly more than expected by chance, *p <* 0.0001), are also successfully reproduced by this biophysical model that simulates anaesthesia as prolonged inhibitory postsynaptic current (Fig. 7). In other words, the model is not trained or tuned based on the hctsa features, but rather using a biologically realistic change in miscroscale postsynaptic potential. Nevertheless, it still manages to reproduce the vast majority of the macroscale changes that we observed empirically.

**Figure 7.**
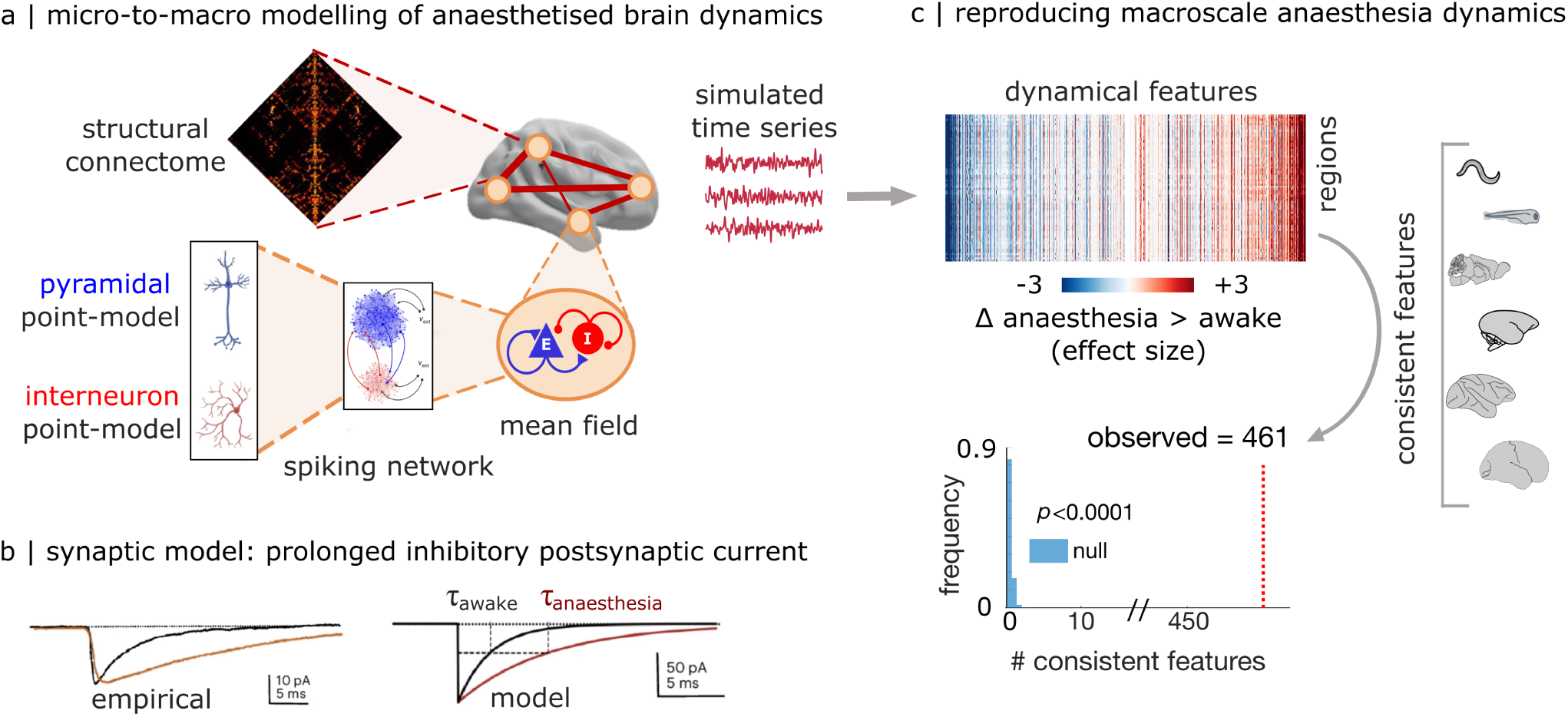
Computational modelling of anaesthetic-induced changes in macroscale local dynamics from microscale synaptic mechanisms. (**a**) Computational model to assess how microscale molecular mechanisms impact macroscale brain dynamics. The whole-brain model consists of biophysically grounded mean-field reductions (one for each brain region) of networks of excitatory and inhibitory neurons, integrating microscale cellular and molecular properties (membrane conductances and AMPA, NMDA, and GABA*_A_* synaptic receptors) to simulate regional fMRI signals. Once connected according to empirical anatomical connectivity of the human brain from in vivo diffusion tractography, the resulting whole-brain model links biophysically meaningful synaptic-level mechanisms to macroscale dynamics. (**b**) At the microscale, the model simulates the empirically-observed prolongation of inhibitory post-synaptic currents induced by GABA-ergic anaesthesia, as an increase in the time constant of inhibitory synaptic decay. (**c**) At the macroscale, we apply comprehensive dynamical phenotyping to the simulated BOLD signals from the model in the awake and anaesthetised regimes (N=19 simulations for each condition), following the same workflow as for our empirical data. Among the 485 features that had exhibited consistent changes across all empirical contrasts from Fig. 3, 461 features (>95%) also exhibit changes in the same direction in the computational model. This number is significantly greater than expected by chance, as assessed by a null model where each feature’s direction of change (increase or decrease) is assigned at random, repeated 10 000 times (*p <* 0.0001).

Though potentiation of inhibitory post-synaptic potentials is a prominent mechanisms for many widely used anaesthetics, including many of those employed in the present study (propofol, sevoflurane, halothane, isoflurane), other cellular- and molecular-level mechanisms may also be at play. Indeed Sacha et al. [76] also provide a model that implements the synaptic-level effects of a different class of anaesthetics, antagonists of the glutamatergic NMDA receptor (e.g., a high dose of ketamine). At the microscale, such drugs induce a shortening of the excitatory postsynaptic current [70, 79], which was implemented in the same multi-scale computational modelling framework developed by Sacha et al. [76], as a decrease in the excitatory synaptic decay time. To ensure robustness and convergence of our modelling results, we therefore also assess this second, distinct model of a biologically observed synaptic-level mechanism of anaesthesia, in terms of its ability to reproduce (without any additional fine-tuning or fitting) the changes in local neural dynamics induced by anaesthesia at the macroscale. We find that this alternative model also successfully reproduces >90% of anaesthetic-induced changes in macroscale local dynamics (Fig. S16). Hence, we demonstrate that the biophysical model’s ability to reproduce the empirical effects of anaesthesia on local dynamics is not critically dependent on the specific combination of model parameters. Instead, having found that anaesthetics with different molecular targets converge to produce convergent macroscale effects on brain dynamics (and ultimately behaviour); we can now provide a computational mechanism whereby different biologically plausible synaptic mechanisms can induce those same convergent effects at the macroscale.

### Sensitivity, robustness and validation

To ensure that results are not due to specifics of analytic choices, data selection, or processing, we conduct additional sensitivity and robustness analyses and validation. First, we ask: could the number of features exhibiting consistent change across every anaesthetic and species, occur purely due to chance alone, given the large number of features that we test? To test this null hypothesis, we construct a null distribution whereby the sign of each feature and contrast is assigned at random (+1 or -1). Repeating this process 10,000 times produces on average fewer than one feature exhibiting consistent sign across all 15 contrasts (*mean* = 0.41, in line with the analytic calculation that one should expect one feature for every 16,384 features). The number of consistent features observed under the null never approaches the empirically observed number of 485 consistent features, yielding a p-value of *p <* 0.0001 (Fig. S17).

Next, we consider the temporal resolution. Within each dataset, the temporal resolution is matched for awake and anaesthetised acquisition. Additionally, the datasets included here have broadly comparable temporal resolution (in the order of seconds; range 0.5-2.4s). Nevertheless, to further demonstrate robustness of our analyses to the differences in temporal resolution between datasets, we repeated the main analysis after harmonising all datasets by downsampling to the slowest temporal resolution among them (2.4s). We show that clear consistency across species is still observed (Fig. S18); in fact, the number of features that exhibit consistent anaesthetic-induced changes across all 15 contrasts is now approximately 11% greater (540; Fig. S19).

Having shown that the PLS analysis for gene-dynamics association is robust to use of all consistent features rather than the catch22 subset (Fig. S14 and Fig. S15), we also show that results of the PLS analysis also remain robust upon including additional contrasts for the human sevoflurane dataset, namely 2% vol and approximately 4.4% vol, i.e., both lower and higher than the dose used for the main analysis (Fig. S20). Likewise, results from the PLS analysis performed using all consistent features remain robust upon excluding the nematode and zebrafish data from the identification of consistent features, showing that inclusion of these non-mammalian, non-fMRI datasets does not bias the findings (Fig. S21). We also show that PLS results are not critically dependent on any individual species included. We perform a leave-one-species-out cross-validation analysis, whereby we systematically repeat the genes-features PLS analysis after excluding one species (human, macaque, marmoset, mouse) each time. We find that the gene- and feature-loadings for each of the main three latent variables are consistently significantly correlated between the 4-species PLS analysis, and each of the 3-species analyses (the only exception being LV3 for the analysis without macaque; Fig. S22). In other words, the latent variables of gene-dynamics association identified in any 3 species will generalise to the 4-species case.

To provide further biological validation, we show that the PLS brain scores obtained from human microarray transcriptomics are strongly and significantly correlated with the brain score obtained from human RNAseq gene expression data instead (Fig. S23). Likewise, the PLS brain scores obtained from the *in situ hybridization* mouse database can be reproduced when using mouse gene expression from two recently released alternative databases using MERFISH [80, 81] (Fig. S24). Additionally, one of the genes exhibiting the strongest spatial association with the significant latent variable is *PVALB/Pvalb*, which encodes the protein parvalbumin, a marker of inhibitory interneurons. We validate this finding in the macaque brain, for which immunohistochemically-derived parvalbumin density data are available for several cortical regions [82]. As expected, we find a significant negative spatial correlation between the PLS brain scores for macaque, and the regional protein density of parvalbumin, consistent with the negative loading of the *PVALB* gene (Fig. S25). Thus, we show that our gene-dynamics association is robust both to use of a broader set of dynamical features, and to use of a different modality for human and mouse gene expression.

## DISCUSSION

Processing information to guide engagement with the environment is a fundamental function of all nervous systems, which is consistently suppressed by anaesthesia across species [6]. Guided by the hypothesis that different anaesthesics may exert their shared effects by inducing similar changes in local neural dynamics as a final common pathway, here we systematically sampled over 6 000 features of local neural dynamics, to triangulate on a conserved dynamical signature of anaesthesia across imaging modalities, scales, species, and anaesthetics. This comprehensive strategy is intended to overcome a fundamental limitation of the existing literature (despite some notable exceptions [24–27]), whereby each study only picks a handful of ad-hoc measures for analysis, potentially leaving large gaps in our understanding of how anaesthetics reshape neural dynamics. We identified an evolutionarily conserved dynamical profile of anaesthesia, indicating that what is common across species and anaesthetics is not just the behavioural response to anaesthesia (isolation from the environment), but also anaesthesia’s effect on specific features of neural activity. Anaesthetics destabilise local dynamics, leading to reduced temporal autocorrelation, shorter intrinsic timescale, and faster decay of neural activity. This effect is reversed by deep-brain stimulation of the centromedian thalamus in the macaque, whereupon the animal re-awakens, providing a further link between neural and behavioural effects. The finding that anaesthetics’ suppression of sensation and action is consistently accompanied by specific changes in neural activity hints at the existence of evolutionarily conserved circuit dynamics that may be necessary for information processing.

Broadly, the present findings suggest that in the anaesthetised brain, intrinsic neural timescales become systematically shorter. Although the time-series features that characterise the dynamical profile of anaesthesia originate from different fields of science and perform different computations on the input time-series of neural activity, they are broadly related to reduced stability and greater unpredictability of the signal’s future from its past, coinciding with decreased measures of linear and nonlinear autocorrelation. In the awake mammalian brain, short timescales (whereby autocorrelation decays quickly) are thought to encode rapidly-changing sensory information. Conversely, long timescales reflect longer ‘temporal receptive windows’, supporting integrative processes and the encoding of contextual information [2, 4, 48–53, 83–88]. Indeed, recent applications of highly-comparative time-series analysis have highlighted the prominence of autocorrelation-related measures for both brain organisation [36] and human aging [39, 40]. In turn, shorter time-scale means that the brain is less capable of encoding contextual information in its ongoing dynamics, and less capable of integrating information over time [4, 85, 86, 89]. This faster decay of local activity, preventing integration, may explain why under anaesthesia incoming stimuli fail to escalate the processing hierarchy and achieve global relevance [8, 43, 90– 92], up to the point that the organism will fail to respond even to noxious stimuli.

The present finding of a shared dynamical signature of anaesthesia, obtained by formal comparison across several anaesthetics and six distinct species from human to nematode, dovetails with a growing number of individual reports that sought to identify signatures of anaesthesia in neural dynamics – with many returning similar results when applied in different species [9–16, 22–24, 93–95]. Consistent with the present work, shorter fMRI intrinsic timescale has been previously reported in humans under deep general anaesthesia (but not under mere sedation) and in patients with disorders of consciousness [96]; reduced fMRI autocorrelation was also reported in anaesthetised macaques [11]. Likewise, under anaesthesia, human fMRI exhibits a reduction in the power of slow frequencies [97]; a shift to faster neuronal activity was also reported in anaesthetised *c.elegans* with 2Hz calcium imaging [46], and medetomidine-isoflurane anaesthesia in mice is known to shift the spectral components of fMRI signal fluctuations towards higher frequencies [98, 99]). At first blush, the shorter intrinsic timescale in the BOLD signal during anaesthesia identified in the present work and in these previous reports may appear at odds with results from electrophysiology. Anaesthesia is often associated with slowing down of fast cortical electrodynamics, as measured by increased prevalence of electrophysiological slow-waves, slower response time to perturbations in macaque local field potentials (LFPs), and longer autocorrelation of the EEG signal [4, 21, 100–104]. However, both the haemodynamics and calcium imaging used here measure neural activity at several orders of magnitude slower than EEG and LFPs (ranging from 500ms for our GCamP data in *c.elegans* to 2400ms for the fMRI, as opposed to the millisecond-level resolution of electrophysiology). Shorter fMRI/calcium imaging timescale is therefore not incompatible with slowing down of EEG activity. To provide practical support for this theoretical argument, we also provide a toy example (Fig. S26), showing that the same time-series can appear slower and more synchronised or faster and less synchronised, depending on whether they are sampled at a fast or slow rate. Together with the findings in the present report, these observations in fMRI and electrophysiology collectively suggest convergence towards a narrower, less diverse range of timescales during anaesthesia, with infraslow fMRI becoming faster, and fast electrophysiology becoming slower.

In addition to reshaping local dynamics, we find that anaesthesia desynchronises regions with similar dynamical profiles, such that they no longer engage in timelocked activity even though they have similar ongoing dynamics. Corroborating the present results, reduced BOLD signal co-fluctuations were previously identified with a variety of anaesthetics in humans [96, 105–108], macaques [11], marmosets [109], and rats [110], but decorrelations have also been reported at the neuronal level in flies and mice [111, 112]. Notably, Huang and colleagues reported lower synchrony and reduced autocorrelation as shared fMRI signatures of unconsciousness between anaesthesia and disorders of consciousness, hinting at an even more fundamental link with consciousness [96]. These results are also in excellent agreement with recordings from local field potentials and intracranial electrocorticograms in humans, which showed that under propofol anaesthesia, neuronal activity becomes desynchronised and functionally fragmented across both time and space [113]. Here we confirm that desynchronisation of slow neural dynamics generalises even more broadly across species and across scales, from neuronal calcium imaging to mammalian haemodynamics. The present results therefore lend support to theoretical accounts wherein functional uncoupling/unbinding/failure of integration was pinpointed as potential ‘final common mechanism’ or common endpoint of how diverse anaesthetics exert their shared behavioural effects: a hypothesis that has stood the test of time for over two decades [6–8, 32, 33]. Anaesthetic-induced breakdown of the relationship between local and global activity is also consistent with work showing that under anaesthesia and sleep, spontaneous and stimulus-evoked activity fail to propagate globally: a potential account for the concomitant loss of behavioural responsiveness [43, 92, 114, 115]. Indeed here we found that when the effects of anaesthesia on the brain’s dynamical phenotype are reversed upon centromedian thalamic deep-brain stimulation in the macaque, the animal also reawakens from anaesthesia [43]. Putting the region- and network-level perspectives together, the present findings suggest that under anaesthesia, local neural activity becomes functionally isolated in space and time: less able to influence processing at other regions (reduced functional synchrony) and also less able to persist in time and influence future activity (reduced autocorrelation) – generating a dynamical regime that may be altogether ‘inhospitable to the information exchange […] critical to the coordination of events that allows both the sensation and response that is characteristic of a bidirectional relationship with the environment’ [6].

If the effects of anaesthesia on local dynamics are stereotyped across species, do they also have a common molecular origin? We find that the conserved changes in local neuronal dynamics induced by anaesthesia covary spatially with corresponding conserved patterns of gene expression across four mammalian species, from rodents to humans. Broadly, two main themes emerge from the pattern of molecular underpinnings identified by multivariate analysis of genes-dynamics relationships. On one hand, the involvement of biological substrates for brain rhythms and timescales: *PVALB* is the marker gene for a key class of fast-spiking inhibitory interneurons involved in generation of brain rhythms [116]; *MBP* is fundamental for myelination, enhancing speed of signal transmission [117]; and the hyperpolarization-activated cyclic-nucleotide (HCN) gated channels, which are present from invertebrates to mammalians, play a pace-maker role in the brain [118]. These results lend support to the growing appreciation for the role of genetic and molecular contributors to neural dynamics and timescales [1, 2, 9, 31, 37, 119–121], and dovetail with the prevalence of features pertaining to signal timescale and autocorrelation in our unbiased screening of anaesthetic effects. Indeed, the pace-maker HCN channels are sensitive to many volatile and intravenous anaesthetics [6, 41, 55, 122–124], and *HCN1*-mediated inhibition was recently identified as a shared mechanism of propofol and ketamine anaesthesia in humans [20]. On the other hand, we found prominent involvement of excitation and inhibition: genes coding for inhibitory receptors and interneurons (*GABRA1, HCN1, CHRM2, PVALB*) predominantly cluster at one end of the multivariate gene-dynamics gradient, with excitatory receptors (*GRIA1, GRIN3A, GRM1, GRM5, CHRNB2*) predominantly clustering at the other end–with both ends including genes pertaining to prominent anaesthetic targets, such as the inhibitory GABA*_A_* and excitatory NMDA receptors.

Our biophysical computational model offers further mechanistic support for key roles of both excitation and inhibition, and local timescales. The model links synaptic-level biophysical changes in local activity, based on microscale experimental findings, to their macroscale dynamical consequences [76]. Anaesthesia is implemented in two distinct ways, corresponding to the main molecular mechanisms of GABA-ergic anaesthetics (such as propofol, sevoflurane) and ketamine anaesthesia [41, 55]. In one case, the time constant of inhibitory synaptic decay is increased, effectively potentiating the inhibitory postsynaptic current. In the other case, the excitatory synaptic decay time is decreased. Hence both mechanisms involve an interaction between timescale and excitation or inhibition. Intuitively, both have the effect of reducing the postsynaptic neuron’s ability to integrate incoming signals into a response: either by shrinking the temporal window over which an excitatory signal can facilitate subsequent postsynaptic spiking (which in turn makes it less likely that enough such signals will be received together); or by increasing the time over which an inhibitory signal can keep the postsynaptic neuron silenced. Altogether, our multi-scale biophysical model demonstrates that a shorter timescale of postsynaptic excitation (resp., prolonged inhibitory timescale) at the microscale, mechanistically translates into shortening of the timescale of neural activity at the macroscale: not only at the level of individual neurons, but also entire brain regions – recapitulating our empirical results.

The present modelling strategy was driven by our interest in shared mechanisms, rather than any specific drug: following [76], we simulated two prominent synaptic-level mechanisms of anaesthesia that are shared by many individual drugs, each representing a broad class of anaesthetics [41, 68, 70, 76–79]. However, alternative modelling strategy are also possible. For example, each anaesthetic has a unique profile of molecular affinities [55]. Hence researchers interested in simulating a specific drug may choose to further increase the biological realism (and complexity) of the model by enriching it with additional receptor- and molecular-level mechanisms, or additional cell types and their regionally heterogeneous distribution [76]. Indeed, propofol and ketamine also share additional mechanisms that were not implemented in the present model, such as *HCN1*-mediated inhibition [20]. Other relevant biological mechanisms shared by many anaesthetics that could be implemented in the model include the “synaptic power failure” account of anaesthesia, whereby volatile anaesthetics hinder excitatory synaptic function by decreasing the availability of presynaptic ATP [125, 126]. When individual differences are of interest, such as for precision medicine, individualised connectomes from a patient’s own diffusion MRI tractography data could also be used to develop personalised models [127]. Ultimately, as the saying goes, ‘all models are wrong, but some are useful’, and the present one is no different.

Here the model parameters for wakefulness and anaesthesia were fixed as pre-specified in [76], to match experimental observations. Our modelling strategy was to start from well-characterised biological phenomena at the microscale (changes in excitatory and inhibitory time constants). We then used our model to test the mechanistic hypothesis that such changes in microscale biophysics could account for the empirical changes in macroscale dynamics induced by anaesthesia. In other words, we used the dynamical phenotype of anaesthesia identified by our empirical analysis, as a read-out of the model’s effectiveness. Indeed it is noteworthy that a single synaptic-level change can reproduce to such an extent the rich dynamical signature of anaesthesia observed across diverse species and anaesthetics. An alternative strategy could involve explicitly optimising the model parameters to reproduce this dynamical phenotype of anaesthesia: a much richer model-fitting target than the one used by existing computational models of BOLD signal changes under anaesthesia, which are typically tuned to reproduce a small number of experimentally observed features of brain activity (often just one) [128–133]. Here we show that it is possible to capture a richer readout of dynamical features. This alternative approach of using dynamical features as optimisation targets would effectively use the model, not to test a mechanistic hypothesis, but as a tool for hypothesis-generation about potential biological mechanisms. When validated or refuted *in vivo*, such predictions can then guide the development of more refined models *in silico*, progressively converging on the underlying biological truth [75, 134]. As the present work demonstrates, dynamical phenotyping is widely applicable across imaging modalities and species, and to experimental manipulations ranging from pharmacology to brain stimulation: hence once a dynamical phenotype of interest has been identified, the same framework for model-based hypothesis generation about its mechanistic origins could be applied to the effects of any drug, disease, or other manipulation.

Indeed, rich phenotyping of local dynamics provides deeper insight into the effects of additional experimental manipulations. Concretely, we found that dynamical signatures of anaesthesia are reversed upon centromedian thalamic deep-brain stimulation in the macaque, whereupon the animal also reawakens from anaesthesia despite continuous presence of the anaesthetic – [43]. We demonstrated this bi-directional association by contrasting propofol anaesthesia with centromedian thalamic DBS, which restores arousal and responsiveness to the environment. We also included an even more stringent contrast between centromedian thalamic DBS, and DBS in a control site (ventrolateral thalamus), which fails to reawaken the animal from anaesthesia despite equivalent level of stimulation [43]. Methodologicaly, these findings demonstrate that the dynamical signatures identified here do not merely track the presence of anaesthetic agents in the system; rather, they reflect whether the anaesthetic successfully suppresses the brain’s ability to interact with the environment. More broadly, our evidence from macaque DBS corroborate the recent report from targeted striatal manipulation in mouse, indicating that local manipulations can affect distal BOLD dynamics at the macroscale beyond the targeted region [38]. We note that the electrical stimulation of CT-DBS was not limited to centromedian thalamus but spread beyond this target to neighboring central-thalamic nuclei. As reported in the original study, modelling the volume of activated tissue using the LEAD-DBS macaque toolbox showed that electrical stimulation went above the electrode contact centered in the centro-median thalamus to encompass intralaminar CT nuclei [43]. Thus, the fact that electrical current was delivered to the region of CT is consistent with the literature in rodents, macaque and human primates, where there is convergent evidence from multiple sources that the intralaminar nuclei (including centeromedian-parafasicular group and CL nucleus) have both critical integrative roles in general, and a central role in consciousness, specifically [135–137]. The intralaminar thalamus has strong reciprocal connections with several regions of the frontal cortex, posterior cortical association areas that support poly-sensory integration, and the basal ganglia [135, 138]. Indeed, a recent systematic review of thalamic nuclei and consciousness by Cacciatore et al. [136] reported that across 167 articles encompassing both humans and other species, results highlighted the centromedian-parafascicular complex of the intralaminar thalamus as the nucleus most related to the generation, modulation, and maintenance of the level of consciousness. This evidence therefore converges with our own present findings about the role of the central thalamus in reversing behavioural and neural effects of anaesthesia. To further discriminate between the individual roles of specific nuclei within the intralaminar thalamus, future work may employ alternative stimulation tools than DBS, such as microstimulation and optogenetic tools that are increasingly becoming available in nonhuman primates. Altogether, we show how the brain’s dynamical regime can be modulated bi-directionally by focal thalamic stimulation and global pharmacology, in a way that tracks behavioural outcome.

The consistency of dynamical effects across species suggests that dynamical phenotypes may provide a common frame of reference for cross-species comparison. Identifying mappings between the brains of humans and other species is a perennial question of translational neuroscience. The traditional approach has focused on anatomy, whereby homologous brain areas are identified according to shared morphological or cytoarchitectonic properties, or similar connectional patterns [139–144]. More recently, gene expression has also been used as a high-dimensional common space to embed the cortical architecture of different species, such as human and mouse, without the need to explicitly match regions [62, 145, 146]. A complementary approach involves finding homologs by identifying which brain regions exhibit consistent responses to the same stimuli [147–151], or matching recurrent patterns of whole-brain activity [5, 152, 153]. Here, we combined these two approaches: we used the high-dimensional common space of local dynamics (i.e., same time-series features in every species) to enable inter-species comparison in response to the same intervention (anaesthesia), which does not require direct one-to-one mapping between regions of different species. We then combined the availability of the same features with the availability of transcriptomic maps for the same genes across the same species, to identify consistent mappings between features and genes. Since dynamics are ubiquitous across nervous systems, our findings show that dynamical phenotyping may provide a common reference frame to embed a wide and diverse range of neuroscientific datasets, from human functional MRI to nematode calcium imaging.

Our results indicate that across diverse species and scales, anaesthesia is characterised by consistent deviations from the dynamical regime of the awake brain. The convergence of diverse anaesthetic agents, and our computational modelling, further indicate that distinct synaptic mechanisms can induce such convergent shifts in dynamical regime. It is also likely that further mechanisms may be able to exert equivalent effects. Nonetheless, the question remains: why should such disparate species converge on a similar dynamical state? Several potential answers may be considered, which are not mutually exclusive. First, the neurotransmitter systems that are targets for anaesthesia are evolutionarily ancient, probably having evolved 700-800 MYA, and the parsimony of evolutionary processes has conserved these for increasingly complex functional purposes [154]. Since anaesthetics affect synaptic machinery subserving these systems (including the present PLS analysis identifying key excitatory and inhibitory systems in mammals), this may contribute to explaining the commonality we observe. For species that are phylogenetically closer, such as mammals and especially primates, an additional potential reason for the observed convergence is their anatomical similarity: the evolutionarily conserved neurotransmitter systems that form the targets of anaesthetic agents are also largely conserved in their anatomical distribution, and present at critical sites which provide strategic roles as modulators or gatekeepers. Such fundamental functions may be especially likely to represent a conserved part of the neural blueprint. Affecting function at these strategic locations may in turn represent a critical vulnerability to anaesthetic agents that is shared across several of the species included here. Second, consciousness is perhaps one of the most complex emergent characteristics of neural systems, as argued on multiple empirical and theoretical grounds [16, 155, 156] and this complexity may in turn lead to fragility and vulnerability. Perturbations of the complex polysynaptic systems subserving consciousness and the capacity to process information and adaptively respond to the environment may therefore be more likely to result in failure than less complex functional pathways. Indeed our computational model indicates that distinct synaptic-level mechanisms can converge to produce similar macroscale dynamical changes, dovetailing with observations that several mechanistic pathways may converge to produce similar anaesthetic states [54]. More broadly, most if not all known animal species (including invertebrates) periodically fall into a spontaneous sleep-like state of reduced responsiveness [157, 158]. The reasons behind the need for sleep and its conservation across phylogeny are topics of intense research in their own right [157, 158], and it is possible that some of them may overlap with the reasons behind the evolutionary conservation of response to anaesthetic agents [6], and the general fragility of the dynamical regime required for wakefulness. Our comprehensive characterisation of the awake and anaesthetised dynamical regimes opens an exciting opportunity for future work to address this question, by identifying relevant features of brain dynamics that such explanations should focus on.

It is especially noteworthy that our results are observed across both regional haemodynamics (from BOLD fMRI and fMRI with monocrystalline iron oxide nanoparticle contrast agent) but also single-neuron data from calcium imaging in nematodes and zebrafish. This means that any of our results that are shared across species cannot logically be attributed to putative physiological confounds of the haemodynamic signal such as breathing, cardiac, or vasodilation/vasoconstriction differences. First, because the different anaesthetics included here do not all influence such physiological parameters in the same way, and some are even opposites; for example, volatile anaesthetics are vasodilators, whereas propofol is a vasoconstrictor [6–8, 102, 159]. Second, such extraneous physiological effects are simply not present in the immobilised nematode worm, unlike the behavioural effects of anaesthesia, and unlike our results about neural dynamics, which are shared across species. Third, our contrasts include reawakening induced by thalamic DBS during which a high dose of propofol is still present [43], so that any physiological changes induced by the drug beyond its responsiveness-suppressing effects would be equally present in both conditions, and cannot drive the difference observed between the two conditions. We also included an even more stringent contrast between centromedian thalamic DBS, and DBS in a control site (ventrolateral thalamus), which fails to reawaken the animal from anaesthesia despite equivalent level of stimulation, demonstrating that the location of stimulation matters, not just its intensity [43]. These results demonstrate that the dynamical signatures identified here do not merely track the presence of anaesthetic agents in the system; rather, they reflect whether the anaesthetic successfully suppresses the brain’s ability to interact with the environment. Additionally, except for the human dataset all data were acquired from head-fixed or otherwise immobilised animals, minimising any influences of motion. Altogether, generalisation of our results to nematode and zebrafish calcium imaging represents a strong control that our results cannot be attributed to any fMRI-specific confounds.

The present work should be considered in light of inevitable methodological limitations. Our focus here was on anaesthesia’s effect on the brain’s capacity to process environmental information, which is indexed by loss of behavioural responsiveness—a marker that is widely shared across species and anaesthetics [6, 8, 160]. Although the generalisation of our results across different species is a key strength of the present work, combining datasets from different species and different modalities is challenging due to methodological heterogeneity in acquisition modality and preprocessing. Notably, we did not have the same anaesthetic in every species, introducing heterogeneity of molecular and cellular mechanisms. Although the datasets included here have broadly comparable temporal resolution (in the order of seconds; range 0.5-2.4s), we repeated the main analysis after harmonising all datasets by downsampling to the slowest temporal resolution among them (2.4s), showing similarly consistent results. Likewise, regional gene expression is quantified in different ways for each species: we sought to mitigate this limitation through replication with alternative transcriptomic databases. Another limitation is the small sample size of each individual dataset, due to the combination of technical challenges of performing anaesthesia in the scanner, and the additional ethical and technical challenges of research in nonhuman animals. However, anaesthesia is a powerful perturbation, making its effects evident even in a relatively small sample. Indeed, the present sample sizes are consistent with current publication standards [14, 15, 42– 45]. Additionally, we sought to compensate for the limitation of individual datasets’ sample size by combining multiple datasets and multiple species, and then only focusing on effects that are highly generalisable across species and datasets. Here we believe lies one of the key strengths of the present work.

Although here we were interested in what is similar across species and anaesthetics, differences between species and anaesthetics were also evident: understanding their origins and behavioural relevance may represent an interesting avenue for future pursuit – which our present datasets are not well suited for addressing. Some of these differences are likely biological, due to the different molecular mechanisms of specific anaesthetics, different doses used in different species, or interspecies and inter-individual differences in neural architecture and other biological phenotypes. However, interindividual and inter-species biological variability may also be confounded with methodological differences (for example, all our fMRI data are from mammalian species; some anaesthetics are only used in one species). Disentangling their respective contributions to the differences in dynamical phenotypes will therefore likely require dedicated data, with standardised protocols for neuroimaging acquisition and anaesthetic administration – especially if cross-species differences are to be the focus of interest. Indeed, such differences in anaesthetics and acquisition methodologies across datasets could represent a limitation, if the present study were focused on inter-species differences. Crucially, however, the driving question of the present work was about the shared effects of anaesthesia on local dynamics. Therefore, here we focused only on those effects that are the same across anaesthetics, species, and imaging modalities. Such consistent effects therefore cannot be explained by the differences between drugs, species, or data modalities. In this context, differences in how the various datasets were acquired and processed (e.g., temporal and spatial resolution; with vs without global signal regression) may even be seen as an asset: combining multiple species and anaesthetics strengthens our ability to triangulate on a ‘final common endpoint’ of anaesthesia at the neural level to match the common behavioural endpoint, while convergence of results across extraneous methodological variations can be interpreted as an additional support for the robustness of our findings.

In conclusion, through comprehensive phenotyping of brain dynamics and systematic comparison across species, we have identified a dynamical regime that characterises anaesthetic-induced breakdown of information processing. Under anaesthesia, local neural activity becomes spatio-temporally isolated: both at the level of single neurons, and at the level of brain regions. Deepbrain stimulation of the central thalamus has the opposite effect, demonstrating bi-directional control over brain dynamics. Biophysical modelling provides a mechanistic link between the macroscale dynamical phenotype of anaesthesia, and microscale effects of key molecular targets on the timescales of synaptic excitation and inhibition.

## METHODS

### Human fMRI anaesthesia dataset

The human sevoflurane data included in this study have been published before [42, 161, 162]. For clarity and consistency of reporting, we use the same wording as in those previous publications, and we refer the reader to the original publication for details [42].

#### Participants and ethics

The ethics committee of the medical school of the Technische Universitat Munchen (Munchen, Germany) approved the current study, which was conducted in accordance with the Declaration of Helsinki. Written informed consent was obtained from volunteers at least 48 h before the study session. Twenty healthy adult men (20 to 36 years of age; mean, 26 years) were recruited through campus notices and personal contact, and compensated for their participation in the study. Before inclusion in the study, detailed information was provided about the protocol and risks, and medical history was reviewed to assess any previous neurologic or psychiatric disorder. A focused physical examination was performed, and a resting electrocardiogram was recorded. Further exclusion criteria were the following: physical status other than American Society of Anesthesiologists physical status I, chronic intake of medication or drugs, hardness of hearing or deafness, absence of fluency in German, known or suspected disposition to malignant hyperthermia, acute hepatic porphyria, history of halothane hepatitis, obesity with a body mass index more than 30 kg/m2, gastrointestinal disorders with a disposition for gastroesophageal regurgitation, known or suspected difficult airway, and presence of metal implants. Data acquisition took place between June and December 2013.

#### Anaesthesia protocol

Sevoflurane concentrations were chosen so that participants tolerated artificial ventilation (reached at 2.0 vol%) and that burst-suppression (BS) was reached in all participants (around 4.4 vol%). To make group comparisons feasible, an intermediate concentration of 3.0 vol% was also used. In the MRI scanner, participants were in a resting state with eyes closed for 700s. Since EEG data were simultaneously acquired during MRI scanning [42] (though they are not analysed in the present study), visual online inspection of the EEG was used to verify that participants did not fall asleep during the pre-anaesthesia baseline scan. Sevoflurane mixed with oxygen was administered via a tight-fitting facemask using an fMRI-compatible anaesthesia machine (Fabius Tiro, Drager, Germany). Standard American Society of Anesthesiologists monitoring was performed: concentrations of sevoflurane, oxygen and carbon dioxide, were monitored using a cardiorespiratory monitor (DatexaS, General electric, USA). After administering an end-tidal sevoflurane concentration (etSev) of 0.4 vol% for 5 min, sevoflurane concentration was increased in a stepwise fashion by 0.2 vol% every 3 min until the participant became unconscious, as judged by the loss of responsiveness (LOR) to the repeatedly spoken command “squeeze my hand” two consecutive times. Sevoflurane concentration was then increased to reach an end-tidal concentration of approximately 3 vol%. When clinically indicated, ventilation was managed by the physician and a laryngeal mask suitable for fMRI (I-gel, Intersurgical, United Kingdom) was inserted. The fraction of inspired oxygen was then set at 0.8, and mechanical ventilation was adjusted to maintain end-tidal carbon dioxide at steady concentrations of 33 ± 1.71 mmHg during BS, 34 ± 1.12 mmHg during 3 vol%, and 33±1.49 mmHg during 2 vol% (throughout this article, mean ± SD). Norepinephrine was given by continuous infusion (0.1±0.01*µg* kg-1 min-1) through an intravenous catheter in a vein on the dorsum of the hand, to maintain the mean arterial blood pressure close to baseline values (baseline, 96 ± 9.36 mmHg; BS, 88 ± 7.55 mmHg; 3 vol%, 88 ± 8.4 mmHg; 2 vol%, 89±9.37 mmHg; follow-up, 98±9.41 mmHg). After insertion of the laryngeal mask airway, sevoflurane concentration was gradually increased until the EEG showed burst-suppression with suppression periods of at least 1,000 ms and about 50% suppression of electrical activity (reached at 4.34 ± 0.22 vol%), which is characteristic of deep anaesthesia. At that point, another 700s of electroencephalogram and fMRI was recorded. Further 700s of data were acquired at steady end-tidal sevoflurane concentrations of 3 and 2 vol%, respectively (corresponding to Ramsay scale level 6, the deepest), each after an equilibration time of 15 min. In a final step, etSev was reduced to two times the concentration at LOR. However, most of the participants moved or did not tolerate the laryngeal mask any more under this condition: therefore, this stage was not included in the analysis. Sevoflurane administration was then terminated, and the scanner table was slid out of the MRI scanner to monitor post-anaesthetic recovery. The participants was manually ventilated until spontaneous ventilation returned. The laryngeal mask was removed as soon as the patient opened his mouth on command. The physician regularly asked the participant to squeeze their hand: recovery of responsiveness was noted to occur as soon as the command was followed. Fifteen minutes after the time of recovery of responsiveness, the Brice interview was administered to assess for awareness during sevoflurane exposure; the interview was repeated on the phone the next day. After a total of 45 min of recovery time, another resting-state combined fMRI-EEG scan was acquired (with eyes closed, as for the baseline scan). When participants were alert, oriented, cooperative, and physiologically stable, they were taken home by a family member or a friend appointed in advance.

#### MRI data acquisition

Although the original study acquired both functional MRI (fMRI) and electroencephalographic (EEG) data, in the present work we only considered the fMRI data. Data acquisition was carried out on a 3-Tesla magnetic resonance imaging scanner (Achieva Quasar Dual 3.0T 16CH, The Netherlands) with an eight-channel, phased-array head coil. The data were collected using a gradient echo planar imaging sequence (echo time = 30 ms, repetition time (TR) = 1.838 s, flip angle = 75 ^◦^, field of view = 220 220 mm2, matrix = 72 × 72, 32 slices, slice thickness = 3 mm, and 1 mm interslice gap; 700-s acquisition time, resulting in 350 functional volumes). The anatomical scan was acquired before the functional scan using a T1-weighted MPRAGE sequence with 240 240 170 voxels (1 1 1 mm voxel size) covering the whole brain. A total of 16 volunteers completed the full protocol and were included in our analyses; one participant was excluded due to high motion, leaving N=15 for analysis [42].

#### Functional MRI preprocessing and denoising

We applied a standard preprocessing pipeline in accordance with our previous publications with anaesthesia data [94, 161]. Preprocessing was performed using the *CONN* toolbox, version 17f (CONN; http://www.nitrc.org/projects/conn) [163], implemented in MATLAB 2016a. The pipeline involved the following steps: removal of the first 10s, to achieve steady-state magnetization; motion correction; slice-timing correction; identification of outlier volumes for subsequent scrubbing by means of the quality assurance/artifact rejection software *art* (http://www.nitrc.org/projects/ artifact_detect); normalisation to Montreal Neurological Institute (MNI-152) standard space (2 mm isotropic resampling resolution), using the segmented grey matter image from each participant’s T1-weighted anatomical image, together with an a priori grey matter template.

Denoising was also performed using the CONN toolbox, using the same approach as in our previous publications with pharmaco-MRI datasets [94, 161]. Pharmacological agents can induce alterations in physiological parameters (heart rate, breathing rate, motion) or neurovascular coupling. The anatomical CompCor (aCompCor) method removes physiological fluctuations by extracting principal components from regions unlikely to be modulated by neural activity; these components are then included as nuisance regressors [164]. Following this approach, five principal components were extracted from white matter and cerebrospinal fluid signals (using individual tissue masks obtained from the T1-weighted structural MRI images) [163]; and regressed out from the functional data together with six individual-specific realignment parameters (three translations and three rotations) as well as their first-order temporal derivatives; followed by scrubbing of outliers identified by ART, using Ordinary Least Squares regression [163]. Finally, the denoised BOLD signal time-series were linearly detrended and band-pass filtered to eliminate both low-frequency drift effects and high-frequency noise, thus retaining frequencies between 0.008 and 0.09 Hz.

The step of global signal regression (GSR) has received substantial attention in the literature as a denoising method [165–167]. However, recent work has demonstrated that the global signal contains behaviourally relevant information [168] and, crucially, information about states of consciousness, across pharmacological and pathological perturbations [24]. Therefore, in line with ours and others’ previous studies, here we avoided using GSR to denoise our human fMRI data in favour of the aCompCor denoising procedure, which is among those recommended.

Finally, denoised BOLD signals were parcellated into 100 cortical regions-of-interest (ROIs) from the Schaefer atlas [169].

#### Functional MRI preprocessing and denoising

HCP-minimally preprocessed data [170] were used for all acquisitions. The minimal preprocessing pipeline includes bias field correction, functional realignment, motion correction, and spatial normalisation to Montreal Neurological Institute (MNI-152) standard space with 2mm isotropic resampling resolution [170].

We used the same denoising procedure as for the human anaesthesia dataset, based on the aCompCor method implemented in CONN [163].

### Macaque fMRI anaesthesia datasets

The macaque fMRI datasets used here have been previously reported; for clarity and consistency of reporting, where possible we use the same wording as in these previous reports, and we refer the reader to them for details [15, 22, 43].

#### Animals and ethics

For the macaque Multi-Anaesthesia dataset, N=5 rhesus macaques were included for analyses (*Macaca mulatta*, one male, monkey J, and four females, monkey A, K, Ki, and R, 5-8 kg, 8-12 yr of age), in a total of six different arousal conditions: awake state, deep ketamine, light propofol, deep propofol, light sevoflurane, and deep sevoflurane anaesthesia. Here, we used the awake, deep propofol, deep sevoflurane, and ketamine data. Three monkeys were used for each condition: awake state (monkeys A, K, and J), ketamine (monkeys K, R and Ki), propofol (monkeys K, R, and J), sevoflurane (monkeys Ki, R, and J). Each monkey had fMRI restingstate acquisitions on different days and several monkeys were scanned in more than one experimental condition. Sex was not considered in this study. Because of the small sample sizes, the sex balance per group could not be secured. All procedures are in agreement with the European Convention for the Protection of Vertebrate Animals used for Experimental and Other Scientific Purposes (Directive 2010/63/EU) and the National Institutes of Health’s Guide for the Care and Use of Laboratory Animals. Animal studies were approved by the institutional Ethical Committee (Commissariat a l’Energie atomique et aux Énergies alternatives; France; protocols CETEA 10-003 and 12-086). Additional details of the acquisitions can be found in the original publications [11, 15].

For the macaque DBS dataset, N=5 male rhesus macaques (*Macaca mulatta*, 9 to 17 years and 7.5 to 9.1 kg) were included, three for the awake (non-DBS) experiments (monkeys B, J, and Y) and two for the DBS experiments (monkeys N and T). Only males were included in the DBS dataset in order to avoid the menstrual cycle and hormone variations. All procedures are in agreement with 2010/63/UE, 86-406, 12-086 and 16-040. For additional details, we refer the reader to the original publication [43].

#### Anaesthesia protocol

For the Multi-Anaesthesia dataset, monkeys received anaesthesia either with ketamine, propofol, or sevoflurane [11, 15], with two different levels of anaesthesia for propofol and sevoflurane anaesthesia (light and deep). The anaesthesia levels were defined according to the monkey sedation scale (Table S5), based on spontaneous movements and the response to external stimuli (presentation, shaking or prodding, toe pinch), and corneal reflex [15] and EEG. EEG data were simultaneously acquired during MRI scanning (though they are not analysed in the present study). For each scanning session, the clinical score was determined at the beginning and end of the scanning session, together with continuous visual monitoring of electroencephalography monitoring. Monkeys were intubated and ventilated [11, 15]. Heart rate, noninvasive blood pressure, oxygen saturation, respiratory rate, end-tidal carbon dioxide, and cutaneous temperature were monitored (Maglife, Schiller, France) and recorded online (Schiller).

During deep ketamine, deep propofol, and deep sevoflurane anaesthesia, monkeys stopped responding to all stimuli, reaching a state of general anaesthesia. For ketamine anaesthesia, ketamine was injected intramuscular (20 mg/kg; Virbac, France) for induction of anaesthesia, followed by a continuous intravenous infusion of ketamine (15 to 16 mg · kg^−1^ · h^−1^) to maintain anaesthesia. Atropine (0.02 mg/kg intramuscularly; Aguettant, France) was injected 10 min before induction, to reduce salivary and bronchial secretions. For propofol anaesthesia, monkeys were trained to be injected an intravenous propofol bolus (5 to 7.5 mg/kg; Fresenius Kabi, France), followed by a target-controlled infusion (Alaris PK Syringe pump, CareFusion, USA) of propofol (light propofol sedation, 3.7 to 4.0 *µ*g/ml; deep propofol anaesthesia, 5.6 to 7.2 *µ*g/ml) based on the Paedfusor pharmacokinetic model. During sevoflurane anaesthesia, monkeys received first an intramuscular injection of ketamine (20 mg/kg; Virbac) for induction, followed by sevoflurane anaesthesia (light sevoflurane, sevoflurane inspiratory/expiratory, 2.2/2.1 volume percent; deep sevoflurane, sevoflurane inspiratory/expiratory, 4.4/4.0 volume percent; Abbott, France). Only 80 minutes after the induction, the scanning sessions started for the sevoflurane acquisitions to get a washout of the initial ketamine injection. To avoid artefacts related to potential movements throughout magnetic resonance imaging acquisition, a muscle-blocking agent was coadministered (cisatracurium, 0.15 mg/kg bolus intravenously, followed by continuous intravenous infusion at a rate of 0.18 mg · *kg*^−1^ · *h*^−1^; GlaxoSmithKline, France) during the ketamine and light propofol sessions. Here we include the ketamine, deep propofol, and deep sevoflurane data.

For the DBS dataset [43], anaesthesia was induced with an intramuscular injection of ketamine (10 mg/kg; Virbac, France) and dexmedetomidine (20 *µ*g/kg; Ovion Pharma, USA) and then the same method as reported above for deep propofol sedation was used (Monkey T: TCI, 4.6 to 4.8 *µ*g/ml; monkey N: TCI, 4.0 to 4.2 *µ*g/ml). Awake scanning data were obtained from the remaining three animals, who did not provide anaesthesia data.

#### Deep Brain Stimulation protocol

Two monkeys (N and T) were implanted with a clinical DBS electrode (Medtronic, Minneapolis, MN, USA, lead model 3389). The DBS lead had four active contacts for electrical stimulation (1.5-mm contact length, 0.5-mm spacing, and 1.27-mm diameter). We performed stereotactic surgery, targeting the right centromedian thalamus using a neuronavigation system (Brain-Sight, Rogue, Canada), guided by the rhesus macaque atlases [171, 172] and a preoperative and intraoperative anatomical MRI [MPRAGE (magnetization prepared - rapid gradient echo), T1-weighted, repetition time (TR) = 2200 ms, inversion time (TI) = 900 ms, 0.80-mm isotropic voxel size, and sagittal orientation]. The electrode was stabilized with the Stimloc lead anchoring device (Medtronic, Minneapolis, MN, USA). The extracranial part of the DBS lead was hosted using a homemade three-dimensional (3D) printed MRI-compatible chamber. We waited at least 20 days after implantation before starting the DBS-fMRI experiments. Two methods were used to ensure for the anatomical localization of the DBS lead and the DBS contacts. First, a reconstruction method based on in vivo brain imaging. Second, a histology study in one of the implanted monkeys (for more details, see [43]).

For stimulation, the DBS electrode was plugged to an external stimulator (DS8000, World Precision Instrument, USA), and all the parameters were tuned to a fixed value of frequency (f = 130.208 Hz, T = 7.68 ms), waveform (monopolar signal), and length of width pulse (monkey N, w = 320 *µ*s; monkey T, w = 140 *µ*s). The absolute voltage amplitude was set to 3V (‘low’ DBS) or 5V (‘high’ DBS). The resting-state fMRI experiments were acquired either in the awake state or under propofol anaesthesia without (‘off’ condition) or with a low or high DBS during the entire run on either the CT or VT thalamic nuclei. The DBS started a few seconds before the beginning of the fMRI sequence and stopped just after the end of the MRI sequence [43]. Here, we included the awake and no-DBS conditions, as well as 5V CT and 5V VT stimulation conditions.

As reported by Tasserie et al., [43], ‘DBS significantly affected the general physiology parameters of the anaesthetized monkeys such as the mean heart rate (*P* = 3.38 × 10^−26^) and mean blood pressure (*P* = 5.78 × 10^−17^). For example, for monkey T, high CT-DBS significantly increased mean heart rate (*P* = 1.23 × 10^−23^, compared to anaesthesia; *P* = 6.76 × 10^−19^, compared to low CT-DBS condition) and mean blood pressure (*P* = 8.66 × 10^−14^, compared to anaesthesia condition; *P* = 6.67 × 10^−11^, compared to low CT-DBS condition)’.

#### Behavioural assessment of arousal

We used a preclinical behavioural scale to assess the arousal levels of the monkeys [173]. This scale, based on the Human Observers Assessment of Alertness and Sedation Scale [174] and previously utilised in non-human primate (NHP) research [175], was used consistently across all experimental conditions, in both datasets. The arousal testing occurred outside the MRI environment and was conducted at the beginning and end of each scanning session, for each condition, once the animals were no longer under paralysis. The assessment encompassed six criteria as described in S5. The behavioural score ranged from 0 to 11, where 11 represented the maximum score achievable and 0 the lowest.

In all cases, and for both datasets, we observed no differences in arousal scores between different animals in the same condition. For both datasets, the behavioural score during wakefulness was the maximum of 11/11 for all the animals (Monkey A, Monkey K, and Monkey J from the Multi-Anaesthesia dataset, and Monkeys B, J and Y from the DBS dataset): exploration of the surrounding world = 2; spontaneous movements = 2; shaking/prodding = 2; toe pinch = 2; eyes opening = 2; corneal reflex = 1.

For results pertaining to the different anaesthesia conditions of the Multi-Anaesthesia dataset, see Table S25 of [22]. As a summary, deep anaesthesia with ketamine, propofol, or sevoflurane induced an arousal score of 0, consistently in all animals. In contrast, light anaesthesia induced an arousal score of 3 for sevoflurane, and 4 for propofol.

In the anaesthesia without DBS (‘off’) condition, monkeys N and T displayed the minimum behavioral score of 0 over 11, same as the deep anaesthesia from the Multi-Anaesthesia dataset: exploration of the surrounding world = 0; spontaneous movements = 0; shaking/prodding = 0; toe pinch = 0; eyes opening = 0; corneal reflex = 0. When the CT electrical stimulation amplitude was increased to 5V (high-amplitude CT DBS), animals reached a total score of 9 over 11 (exploration of the surrounding world = 1; spontaneous movements = 1; shaking/prodding = 2; toe pinch = 2; eyes opening = 2; corneal reflex = 1). For VT DBS, both low (3V) and high (5V) amplitude stimulation led to a clinical score of 0, identical to what is observed in the absence of any stimulation.

#### MRI data acquisition

To minimise head motion, monkeys were head-fixed using an implanted with magnetic resonance compatible head post. For the awake scanning sessions, monkeys were trained to sit in the sphinx position in a primate chair inside the dark magnetic resonance imaging scanner and fixate without any task, and the eye position was monitored at 120 Hz (Iscan Inc., USA). The eyetracking was performed to make sure that the monkeys were awake during the whole scanning session and not sleeping [176]. The eye movements were not regressed out from rfMRI data. For the anaesthesia sessions, animals were positioned in a sphinx position, mechanically ventilated, and their physiologic parameters were monitored. No eye-tracking was performed in anaesthetic conditions.

For the Multi-Anaesthesia dataset, before each scanning session, a contrast agent, monocrystalline iron oxide (MION) nanoparticle (Feraheme, AMAG Pharmaceuticals, USA; 10 mg/kg, intravenous), was injected into the monkey’s saphenous vein [177]. Monkeys were scanned at rest on a 3-Tesla horizontal scanner (Siemens Tim Trio, Germany) with a single transmit-receive surface coil customised to monkeys. Each functional scan consisted of gradient-echo planar whole-brain images (repetition time = 2,400 ms; echo time = 20 ms; 1.5-mm3 voxel size; 500 brain volumes per run).

For the DBS dataset, absence of motion was ensured by paralising the animals with curare. Monkeys were scanned at rest on a 3-Tesla horizontal scanner (Siemens, Prisma Fit, Erlanger Germany) with a customised eightchannel phasedarray surface coil (KU Leuven, Belgium). The parameters of the functional MRI sequences were: echo planar imaging (EPI), TR = 1250 ms, echo time (TE) = 14.20 ms, 1.25-mm isotropic voxel size and 500 brain volumes per run. Event-related data pertaining to auditory stimulation were also acquired and are reported in [43], but here we only used the resting-state fMRI data, and will not discuss the event-related data further. Scalp EEG data were also acquired using an MR-compatible system and custom-built caps (EasyCap, 13 channels), an MR amplifier (BrainAmp, Brain Products, Germany), and the Vision Recorder software (Brain Products). These results are reported in [43], but here we did not consider the EEG data and will not discuss them further.

#### Macaque functional MRI preprocessing, denoising, and time-series extraction

For the Multi-Anaesthesia dataset, a total of 157 functional magnetic imaging runs were acquired [15]: Awake, 31 runs (monkey A, 4 runs; monkey J, 18 runs; monkey K, 9 runs), Ketamine, 25 runs (monkey K, 8 runs; monkey Ki, 7 runs; monkey R, 10 runs), Light Propofol, 25 runs (monkey J, 2 runs; monkey K, 11 runs; monkey R, 12 runs), Deep Propofol, 31 runs (monkey J, 9 runs; monkey K, 10 runs; monkey R, 12 runs), Light Sevoflurane, 25 runs (monkey J, 5 runs; monkey Ki, 10 runs; monkey R, 10 runs), Deep Sevoflurane anaesthesia, 20 runs (monkey J, 2 runs; monkey Ki, 8 runs; monkey R, 10 runs). Additional details are available from the original publications [11, 15, 92].

Functional images were reoriented, realigned, and rigidly coregistered to the anatomical template of the monkey Montreal Neurologic Institute (Montreal, Canada) space with the use of Python programming language and FMRIB Software Library (FSL) software (http://www.fmrib.ox.ac.uk/fsl/; accessed February 4, 2018) [15]. From the images, the global signal was regressed out to remove confounding effect due to physiologic changes (e.g., respiratory or cardiac changes).

For the DBS dataset, a total of 199 Resting State functional MRI runs were acquired: Awake 47 runs (monkey B: 18 runs; monkey J: 13 runs; monkey Y: 16 runs), anaesthesia (DBS-off) 38 runs (monkey N: 16 runs,; monkey T: 22 runs), low amplitude centro-median thalamic DBS 36 runs (monkey N: 18 runs; monkey T: 18 runs), low amplitude ventro-lateral thalamic DBS 20 runs (monkey T), high amplitude centro-median thalamic DBS 38 runs (monkey N: 17 runs; monkey T: 21 runs), and high amplitude ventro-lateral thalamic DBS 20 runs (monkey T: 20 runs) [43].

Images were preprocessed using Pypreclin (Python preclinical pipeline) [10]. Functional images were corrected for slice timing and B0 inhomogeneities, reoriented, realigned, resampled (1.0 mm isotropic), masked, coregistered to the MNI macaque brain template [178], and smoothed (3.0-mm Gaussian kernel). Anatomical images were corrected for B1 inhomogeneities, normalised to the anatomical MNI macaque brain template, and masked.

For both datasets, data were parcellated according to the Regional Map parcellation [144]. This parcellation comprises 82 cortical ROIs (41 per hemisphere). Voxel time series were filtered with low-pass (0.05-Hz cutoff) and high-pass (0.0025-Hz cutoff) filters and a zero-phase fast-Fourier notch filter (0.03 Hz) to remove an artifactual pure frequency present in all the data [11, 15, 43].

Furthermore, an extra quality control (QC) procedure was performed to ensure the quality of the data after time-series extraction [92]. This quality control procedure is based on trial-by-trial visual inspection by an expert neuroimager (C.M.S.), and it is the same as was previously implemented in [22, 92]. Its adoption ensures that we employ consistent criteria across our two macaque datasets, by adopting the more stringent of the two. We plotted the time series of each region, as well as the static functional connectivity matrix (FC), the dynamic connectivity (dFC) and a Fourier analysis to detect unconventional spikes of activity. For each dataset, visual inspection was first used to become familiar with the characteristics of the entire dataset: how the amplitude spectrum, time-series, FC and dynamic FC look. Subsequently, each trial was inspected again with particular focus on two main types of potential artefacts. The first one may correspond to issues with the acquisition and is given by stereotyped sinusoidal oscillatory patterns without variation. The second one may correspond to a head or other movement not corrected properly by our preprocessing procedure. This last artefact can be sometimes recognized by bursts or peaks of activity. Sinusoidal activity generates artificially high functional correlation and peak of frequencies in the Amplitude spectrum plot. Uncorrected movements generate peaks of activity with high functional correlation and sections of high functional correlations in the dynamical FC matrix. If we observed any of these anomalies we rejected the trial, opting to adopt a conservative policy. See Figures S19-S21 from [22] for examples of artifact-free and rejected trials.

As a result, for the Multi-Anaesthesia data set a total of 119 runs are analysed in subsequent sections (the same as used in [92]): awake state 24 runs, ketamine anaesthesia 22 runs, light propofol anaesthesia 21 runs, deep propofol anaesthesia 23 runs, light sevoflurane anaesthesia 18 runs, deep sevoflurane anaesthesia 11 runs. For the DBS data set, a total of 156 runs are analysed in subsequent sections: awake state 36 runs, Off condition (propofol anaesthesia without stimulation) 28 runs, low-amplitude CT stimulation 31 runs, low-amplitude VT stimulation 18 runs, high-amplitude CT stimulation 25 runs, high-amplitude VT stimulation 18 runs.

### Marmoset fMRI anaesthesia dataset

The marmoset data included here have been published before [45]. For clarity and consistency of reporting, we use the same wording as in the original publication where possible, and we refer the reader to the original work for details [45].

#### Animals and ethics

This study was approved by the Animal Experiment Committees at the RIKEN Center for Brain Science (CBS) and was conducted per the guidelines for Conducting Animal Experiments of RIKEN CBS. Three male and one female healthy common marmosets (*C. jacchus*) between 3 and 6 years of age were included. All marmosets were examined 8 times to collect functional MRI data in all conditions. After awake data were firstly collected, and sedate/anesthetic data were done in a random order for sedate/anesthetic condition with an interval of 1 month between each examination in each individual. For further details we refer the reader to the original publication [179].

#### Anesthesia protocol

We used data from awake scans, and from scans under general anaesthesia induced with isoflurane, sevoflurane, or propofol. Essential details are reported below, and for additional detail we refer to the original publication [45].

##### Isoflurane

Three percent isoflurane with 100% O2 as carrier gas was administered through a facial mask to the marmosets retained with leather gloves. Once sufficient sedation was achieved, an 8 Fr catheter (Atom multi-use tube, Atom Medical Corp., Tokyo, Japan) was inserted into the trachea as an intratracheal tube. The isoflurane concentration was reduced to 2.5%, and 50 *µ*g/kg of atropine and 3.0 mL of physiological saline were administered subcutaneously to prevent intratracheal secretion and dehydration, respectively. After intratracheal intubation, general anesthesia was maintained with 1.8% isoflurane. The marmosets were connected to artificial ventilation for small animals (SN-480-7, Shinano Seisakusho, Tokyo, Japan), and mechanical ventilation was performed under the following conditions: 50% inspiratory oxygen, 8 mL of tidal volume, and 30 breaths per min RR. PR, SpO2, RR, EtCO2, inspiratory and endtidal isoflurane concentrations, and rectal temperature were measured with a vital sign monitor.

##### Sevoflurane

All procedures were performed in the same way as isoflurane, except for dosages. The induction of general anesthesia, intratracheal intubation, and maintenance of general anesthesia were performed with 5.0 and 3.0% of sevoflurane (Pfizer Japan Inc., Tokyo, Japan), respectively.

##### Propofol

Propofol (12 mg/kg) was administered as induction of general anesthesia over 3 min via an indwelling needle. Atropine (50 *µ*g/kg) was administered subcutaneously to prevent intratracheal secretion. Immediately after bolus administration, the predicted plasma concentration of propofol was titrated to 7–9 *µ*g/mL for intratracheal intubation with continuous infusion. The administration dose and protocol were calculated beforehand using a pharmacokinetic parameter reported by [179] and pharmacokinetic analysis software, NONMEM ver. VII (GloboMax ICON Development Solutions, Ellicott City, MD, USA). Once a sufficient plasma concentration was obtained, the propofol dose was controlled to maintain that concentration. Intratracheal intubation, respiratory management, and vital sign monitoring were performed in the same manner as for isoflurane.

#### MRI data acquisition

All marmosets underwent a surgical procedure to attach a headpost to the cranial bones to prevent head movement during data collection [45]. During imaging, the marmosets were placed on a custom-made imaging table (Takashima Seisakusho Co., Ltd, Tokyo, Japan) and immobilized by fixing the head post using a head post fixing tool attached at a custom-made imaging table in all conditions. The marmosets were fitted with earplugs. A hot water circulator was used during imaging to maintain body temperature 36–38 ^◦^C under all conditions [45].

In Awake condition, data were collected in the dark and monitored with an infrared camera to prevent the marmosets from falling asleep. If the marmosets were observed closing their eyes during a scan, they were awakened with a loud noise before the next scan was started. They were rewarded with a highly palatable food at the end of each imaging. In anesthetic condition, the marmosets were monitored in the same way to observe spontaneous movement or coughing for intratracheal tube. If spontaneous movement or coughing was observed, additional sedatives/anesthetics were administrated and infusion rate or concentration of inhalational anesthesia was increased.

An ultra-high field MRI system with a static magnetic field strength of 9.4 T (Bruker BioSpin, Ettlingen, Germany), a custom-made 8-channel receiver coil for the marmoset head (Takashima Seisakusho Co., Ltd, Tokyo, Japan), and a 154 mm inner diameter transmitter coil (Bruker BioSpin, Ettlingen, Germany) were used to collect structural and functional data. Structural data and T2-weighted images were imaged using rapid acquisition with relaxation enhancement (RARE) sequence with the following conditions and parameters: time repetition (TR)=4331 ms, time echo (TE) = 15.0 ms, FOV = 42.0 28.0 36.0 mm, matrix size = 120 × 80 voxels, resolution = 0.35 0.35 mm, slice thickness = 0.7 mm, number of slices = 52, scan time = 1 min and 26 s, RARE factor = 4. Functional images were captured using a gradient recalled echo-planar imaging (EPI) sequence with the following conditions and parameters: TR = 2,000 ms, TE = 16.0 mm, FOV = 42.0 28.0 36.0,mm matrix size = 60 × 40 voxels, resolution = 0.7 × 0.7 mm, slice thickness = 0.7 mm, number of slices = 52, repetition = 155, scan time = 310s. Functional imaging was performed 12 times per animal, per condition [45].

#### Marmoset functional MRI preprocessing and denoising

After the acquired data were converted to Neuro Informatics Technology Initiative format (NIfTI), the voxel size was changed from 0.7 mm isotropic to 3.5 mm isotropic using SPM (Wellcome Trust Center for Neuroimaging, London, UK). Estimation and correction of geometric distortions induced by magnetic susceptibility were performed with the top-up tool of the FMRIB Software Library (FSL) software (FMRIB, Oxford, UK) because all cross-sections were imaged with a single excitation in EPI. Slice timing correction was performed to correct for signal acquisition timing discrepancies in each section. Realignment was applied to compensate for head movements caused by body movements. The deviations in 6 directions were obtained: x (left/right), y (front/back), z (up/down), pitch (rotational direction of nodding and looking up), roll (rotational direction of moving the ear closer to the shoulder), and yaw (rotational direction of looking left/right). For each measurement time point (TR), the deviation from the reference time point, and the first functional brain image, was determined; and the image was moved and rotated by the rigid body model based on this deviation. The method of finding the parameters of the linear transformation was used to minimize the difference between the first functional brain image and the affine transformation of the series of functional brain images to be corrected, by calculating convergence using the method of least squares. After correcting the spatial scale error between the structural and functional images with co-registration, segmentation was performed to provide information on the tissue to which each voxel belongs in terms of brain tissue classification. The voxels were spatially standardized by normalization, which aligns the voxels to the standard brain image to correct for structural differences between individuals. Smoothing was applied to suppress excessive voxel value fluctuations within individuals and apply normal probability field theory. Functional data were smoothed using spatial convolution with a Gaussian kernel of 2 voxels (7 mm). Then, physiological noise was denoised using ordinary least squares regression with cerebrospinal fluid pulsation, heart rate, and respiratory artifacts as regressors. Temporal band pass filtering was performed by frequency filtering (0.01–0.1 Hz) using the fMRI denoising pipeline of CONN [163]. Finally, preprocessed functional data were parcellated into 70 cortical regions from the marmoset MBM-vM atlas [180].

### Mouse fMRI anaesthesia dataset

The mouse fMRI data used here have been published before [14]. For clarity and consistency of reporting, where possible we use the same wording as in the original publication [14].

#### Animals and ethics

In vivo experiments were conducted in accordance with the Italian law (DL 26/214, EU 63/2010, Ministero della Sanita, Roma) and with the National Institute of Health recommendations for the care and use of laboratory animals [14]. The animal research protocols for this study were reviewed and approved by the Italian Ministry of Health and the animal care committee of Istituto Italiano di Tecnologia (IIT). All surgeries were performed under anesthesia.

Adult (*<* 6 months old) male C57BL/6J mice were used throughout the study. Mice were group housed in a 12:12 hours light-dark cycle in individually ventilated cages with access to food and water ad libitum and with temperature maintained at 21 1 degrees centigrade and humidity at 60 ± 10%. All the imaged mice were bred in the same vivarium and scanned with the same MRI scanner and imaging protocol employed for the awake scans (see below).

A first group of mice (n = 10, awake dataset) underwent head-post surgery, scanner habituation and fMRI image acquisitions as described below. See [14] for the full surgical, habituation, and scanner protocol. The scans so obtained constitute the awake rsfMRI mouse dataset we used throughout our study. Two additional groups of age matched male C57BL/6J mice were used as reference rsfMRI scans under anesthesia.

#### Anaesthesia protocol

The first group of animals (n = 19, halothane dataset) was previously scanned under shallow halothane anaesthesia, 0.75% [98]. The employed anaesthesia regimen is well characterized [14, 181]; it is representative of the network architecture observed with different anaesthesia regimens in rodents [99] and it exhibits rich spatiotemporal dynamics by preserving spectral properties of fMRI signal fluctuations [98].

A second, separate group (n = 14) of mice were imaged under medetomidine-isoflurane anesthesia (0.05 mg/kg bolus and 0.1 mg/kg/h IV infusion, plus 0.5% isoflurane) [14]. While this anesthetic combination is known to shift the spectral components of fMRI signal fluctuations towards higher frequencies (hence departing from the characteristic 1/f power law distribution that characterizes awake and halothane rsfMRI datasets [98, 99]), it nonetheless represents the most-widely anesthetic mixture used in the rodent imaging community [99].

#### MRI data acquisition

To prevent motion, each mouse was secured using an implanted headpost in the custom-made MRI-compatible animal cradle and the body of the mouse was gently restrained (for details of the headpost implantation and habituation protocol, see the original publication [14]). For scanning under anesthesia, mice were first deeply anesthetized with isoflurane (4% induction), intubated and artificially ventilated (90 BPM). In one group, anesthesia was then switched to halothane (0.75%). In a second group, a bolus of medetomidine (0.05 mg/kg) was given via tail vein cannulation before waiting 5 minutes and starting an infusion of medetomidine (0.1 mg/kg/h) with isoflurane reduced to 0.5%. In both cases, the rsfMRI acquisition started 30 minutes after the switch to light anesthesia [14].

All scans were acquired at the IIT laboratory in Rovereto (Italy) on a 7.0 Tesla MRI scanner (Bruker Biospin, Ettlingen) with a BGA-9 gradient set, a 72 mm birdcage transmit coil, and a four-channel (awake, halothane) or three-channel (medetomidine-isoflurane) solenoid receive coil. Awake and medetomidine-isoflurane rsfMRI scans were acquired using a singleshot echo planar imaging (EPI) sequence with the following parameters: TR/TE=1000/15 ms, flip angle=60 degrees, matrix=100 x 100, FOV=2.3 × 2.3 cm, 18 coronal slices (voxel-size 230 230 600 mm), slice thickness=600 mm and 1920 time points, for a total time of 32 minutes. Mice under halothane anesthesia (n = 19) were scanned with a TR/TE=1200/15ms, flip angle=60 degrees, matrix=100 100, 24 coronal slices (voxel-size 200 200 500 mm), for a total of 1600 time points, total acquisition time of 32 minutes as described in [14].

#### Mouse functional MRI preprocessing and denoising

Preprocessing of fMRI images was carried out as described in previous work [14]. Briefly, the first 2 minutes of the time series were removed to account for thermal gradient equilibration. Functional MRI time-series were then time despiked (3dDespike, AFNI), motion corrected (MCFLIRT, FSL), skull stripped (FAST, FSL) and spatially registered (ANTs registration suite) to an inhouse mouse brain template with a spatial resolution of 0.23 0.23 0.6 mm^3^. Denoising involved the regression of 25 nuisance parameters. These were: average cerebral spinal fluid signal plus 24 motion parameters determined from the 3 translation and rotation parameters estimated during motion correction, their temporal derivatives and corresponding squared regressors. No global signal regression was employed. In-scanner head motion was quantified via calculations of frame-wise displacement (FD). Average FD levels in awake conditions were comparable to those obtained in anesthetized animals (halothane) under artificial ventilation (p = 0.13, Student t-test) [14]. To rule out a contribution of residual head-motion, we further introduced frame-wise fMRI scrubbing (FD > 0.075 mm). The resulting time series were band-pass filtered (0.01-0.1 Hz band) and then spatially smoothed with a Gaussian kernel of 0.5 mm full width at half maximum. Finally, the time-series were trimmed to ensure that the same number of timepoints were included for all animals, resulting in 1414 volumes per animal. Finally data were parcellated into 72 cortical symmetric regions from the Allen Mouse Brain Atlas (CCFv3).

### Larval zebrafish calcium imaging dataset

#### Animals and ethics

All experiments were conducted on Tg(elavl3:H2BGCaMP6s) zebrafish larvae (*Danio rerio*) at 6 or 7 days post-fertilization (dpf). Sex cannot be determined at the larval stage. This transgenic line expresses a nuclear-localized calcium sensor pan-neuronally [182]. Larvae were raised in embryo medium in an incubator at 28oC on a 14-hour/10-hour day/night cycle. From 5 dpf onward, the medium was replaced daily and larvae were fed live Tetrahymena thermophila CU428.2 (Cornell University). All protocols were approved by the animal care committee of Université Laval (CPAUL protocol 2022-1012).

#### Anaesthesia protocol

Larvae were immobilized in 2% low-melting point agarose (Invitrogen 16520100) in a 30-mm glass-bottom petri dish (Mattek P35G-1.0-14-C). Once solidified, agarose was carefully removed around the tail below the swim bladder using a small scalpel (Fisher 35-205) to allow tail monitoring (Flir BFS-U3-28S5M-C camera with Navitar Zoom 7000 lens) while keeping the head completely immobilized during brain imaging experiments. The imaging chamber was submerged in standard embryo medium, then larvae were brought under the microscope and imaged under baseline awake conditions for 10 minutes, with constant red illumination projected under the chamber (AAXA Technologies KP-750-00 DLP Projector with a Kodak dark red filter). After baseline imaging, a tricaine methanesulfonate solution (MS222, Sigma E10521, sodium channel blocker, 750 mg/L dissolved in embryo medium with pH adjusted using NaHCO3 Fisher, S233-500) [183] was introduced into the imaging chamber using a gravity perfusion system (ALA Scientific Instruments). Drug perfusion was sustained for at least 5 minutes to clear out the chamber volume several times and reach a stable incubation concentration. Full anaesthesia was confirmed in each animal through the complete loss of tail and ocular movements, assessed using both a behavioral monitoring camera or using autofluorescence from the eyes in brain-wide recordings. Imaging experiments lasted between 20 and 30 minutes, with variable recording lengths during the anaesthesia period due to variations in perfusion velocity. The drug concentration used was within established safety margins [183], and full behavioral recovery was confirmed after the drug washout in subjects that were kept in separate petri dishes with fresh embryo medium.

#### Calcium imaging of neuronal activity

Imaging experiments were conducted using equipment and protocols that were detailed in a previous publication [184]. Briefly, brain-wide calcium activity was captured at single-cell resolution using a resonant-scanning two-photon microscope (Scientifica SliceScope with SciScan LabView software) equipped with a piezo-driven water-dipping objective (Nikon, 16x, 0.8 NA). Volumetric stacks of 21 imaging planes (zoom 1.4x, 512x730 pixels, 12 microns in z-spacing) were acquired at a volume rate of 0.986 Hz for a total recording length of 20-30 minutes. Imaging planes intersected roughly 50,000 neurons spanning most brain regions from dorsal to ventral anatomical locations [184]. GCaMP6s fluorescence was excited at a 920 nm wavelength using a Spectra Physics Insight X3 tunable laser and detected using a GaAsP photomultiplicator tube. Laser power after the objective was fixed below 20 mW. Brain imaging ran continuously throughout the drug perfusion phase, which caused flow-induced z-drift in the preparation. Full recordings were thus truncated into two different segments during which there was no solution flow and samples had fully stabilized. Experiments in which z-drift persisted after perfusion were discarded entirely. After functional imaging, anatomical stacks (2 micron z-spacing, 24x frame averaging) were acquired near the isosbestic (calcium-independent) excitation wavelength (860 nm) of the fluorescent sensor, yielding homogeneous nuclear intensities used for anatomical registration in subsequent steps, as per [184].

#### Processing of neuronal activity traces

Calcium imaging planes were motion corrected using the NoRMCorre algorithm [185] implemented within the CaImAn package [186]. Fluorescent nuclei were identified using local intensity maxima in the temporal averages of each calcium imaging plane [187], and fluorescence traces were extracted from each nucleus within a small disk (3-pixel radius). Each trace was detrended from slow baseline drifts and converted to relative Δ*F/F*_0_ units using a smoothed minimum filter applied within a 60-second temporal window. Large fluorescence transients observed at the onset of the drug delivery phase were excluded from anesthetized traces to capture steady-state activity. To situate each neuron in anatomical coordinates, calcium imaging volumes were registered to the mapZebrain larval brain atlas [188] using a step-wise ANTs registration procedure described previously [184, 189]. Following registration, native neuron coordinates were transformed to brain atlas coordinates, then each neuron was assigned to one of 200 anatomical locations (3D regions of interest, ROIs) per brain hemisphere. ROIs were established by subdividing 65 brain region masks uniformly using an iterative clustering approach. Briefly, for each brain region and within each hemisphere, a number of clusters was first determined based on the region’s volume (larger regions requiring more clusters). K-means clustering was then applied 30 times on the voxel coordinates of a region’s 3D mask, yielding a set of cluster centroids that was then re-clustered iteratively to yield consensus centroids across all animals. Mask voxels were finally assigned to the consensus clusters based on their nearest neighboring centroid, effectively partitioning each 3D mask into a Voronoi diagram. This approach was used to obtain a larger set of ROIs (groups of neighbouring neurons) with comparable volumes, thereby providing a finer anatomical resolution and reducing neuron sampling biases. Following this parcellation procedure, the fluorescence signals of individual neurons were averaged within each ROI, and 48 ROIs were excluded from further analysis as they did not contain sampled neurons in at least one animal. This resulted in 352 coarse-grained and anatomically comparable activity traces per larva. Time series were finally truncated to 250 time steps in both awake and anesthetized conditions to obtain comparable lengths across all conditions and animals.

### Nematode calcium imaging dataset

The *C. elegans* data included in this study have been published before. For clarity and consistency of reporting, where possible we use the same wording as in the original publications [44, 46].

#### C. elegans strains

All experiments were on young adult hermaphrodites of the transgenic strain QW1217 (zfIs124[Prgef1::GCaMP6s]; otIs355[Prab-3::NLS::tagRFP]). GCaMP6s, a fluorescent calcium reporter, and nuclear-localized red fluorescent protein are both expressed pan-neuronally in this strain (gift of M. Alkema, University of Massachusetts, Worcester, Massachusetts). *C. elegans* were cultivated at 20 ^◦^C on nematode growth medium seeded with Escherichia coli OP50 [46].

#### Anaesthesia protocol

*C. elegans* become anesthetized on exposure to isoflurane with a minimum alveolar concentration (MAC) value of approximately 3% at room temperature. At 17 ◦C, isoflurane is 2.3 times more soluble in tissues such as muscle and fat than at 37 ^◦^C and consequently absorption of a larger quantity of isoflurane is required to produce a similar chemical potential gradient at cooler temperatures. A concentration of 3% isoflurane at 17 ^◦^C is thus pharmacodynamically similar, with respect to its physical chemistry, to a concentration of 1.3% at 37 ^◦^C. Alterations in *C. elegans* neuronal activity in response to isoflurane exposure were assessed using two experimental regimes: stepwise anesthetization and emergence [46]. Here we include data from the stepwise anesthetisation regime.

Three 5-min-long neuronal activity recordings were taken from each animal (n = 10) after progressive equilibration to 0%, 4%, and 8% atmospheric isoflurane [46]. These concentrations correspond to 1.3 and 2.6 MAC, respectively. For the main analysis we include data from the 0% and 4% isoflurane conditions. Imaging was performed at time t = 30 min, 150 min, and 270 min from the start of the experiment at isoflurane levels of 0%, 4%, and 8%. Isoflurane levels were stepped by exchanging the immersion medium for fresh S-basal buffer with the addition of 13 or 26 *µ*l of pipetted isoflurane for 4% and 8% isoflurane, respectively. The atmosphere within the covered Petri dish was then equilibrated, during the time between imaging sequences, to a concentration of 4% or 8%, respectively, using continuous monitoring with an infrared spectrometer (Ohmeda 5250 RGM; GE Healthcare, USA) and instillation of isoflurane via syringe pump as necessary to maintain the targeted concentration [46].

#### Calcium imaging of neuronal activity

To prevent any motion, the animals were immobilized for imaging by encapsulation in a pad of permeable hydrogel consisting of 13.3% polyethylene glycol diacrylate (Advanced BioMatrix, USA) with 0.1% Irgacure (Sigma-Aldrich, USA) [46]. Hydrogel pads containing animals to be imaged were cured with ultraviolet light onto silanated glass coverslips, which were affixed to the bottom of 50-mm Petri dishes with vacuum grease. Petri dishes were then filled with 50 mL of S-basal buffer (100 mM NaCl, 50 mM KPO4 buffer, and 5 *µ*g/ml cholesterol) as the immersion medium. Tetramisole was added to this buffer at 5 mM to further immobilize the animals.

GCaMP6s and red fluorescent protein fluorescence in the *C. elegans* head ganglia were captured in volumetric stacks using a dual inverted selective plane illumination microscope (Applied Scientific Instrumentation, USA) and water-immersed 0.8 NA 40x objective (Nikon, USA). GCaMP6s and red fluorescent protein were respectively excited using 5-mW 488-nm and 561-nm lasers (Vortan Laser Technology, USA). Volumes for each fluorescent channel (GCaMP6s and red fluorescent protein) were obtained at rate of 2 Hz.

For each animal imaged, 120 neurons (ordered from anterior to posterior in each animal) were tracked in the head region using the nuclear-localized red fluorescent protein fluorophore, and their activity was extracted using the fluctuations in cytoplasmic GCaMP6s neurofluorescence. This tracking and extraction procedure was performed as a massively parallel computation executed at the Massachusetts Green High Performance Computing Center using computational techniques as previously detailed by [44].

#### Preprocessing of neuronal activity traces

In recordings from the stepwise equilibration regime, neuronal activity intensity for all three recordings in each animal (0%, 4%, and 8%) was normalized against the average intensity across all neurons and time points in the 0% isoflurane recording. The first 100 time-points were excluded to remove large transients, leaving 500 for analysis. Because the animals are immobilized and encapsulated in a hydrogel for imaging, it is not possible to normalize to an alternative behavioral endpoint [46].

### Comprehensive dynamical phenotyping

#### Highly-comparative time-series analysis

To extract the dynamical phenotype of each brain region or neuron, we performed massive time-series feature extraction using the *highly comparative timeseries analysis* toolbox, hctsa [29, 30]. For each region/neuron, in each individual, under each condition, the hctsa toolbox extracted >7 200 univariate dynamical features, derived from diverse fields including neuroscience, physics, ecology and economics [29, 30]. Features range from basic statistics of the distribution of time-points, linear correlations among time-points, and stationarity, to measures of entropy, time-delay embeddings, and signal complexity, among others. Each individual feature is the implementation of a computation (termed ‘master operation’) on the input time-series, using specific parameters. For example, sample entropy (SampEn) computes the probability that similar sequences of observations in a time-series will remain similar as their size increases. Its computation therefore requires a threshold *r* for deciding when two sequences will be considered similar; and an embedding dimension *m* that determines the size of the sequences. Multiple individual features are obtained from this master operation as different combinations of *r* and *m*.

We performed an initial pre-filtering, and did not compute any features that had returned *NaN* across all regions, or that had displayed no variance, in an independent dataset of human functional MRI (Human Connectome Project; [190]). The values of different features can vary across several orders of magnitude. Unless otherwise specified, here we do not normalise features across regions, since this could obscure differences across conditions. Instead, we use effect sizes computed on the original features. Effect sizes are expressed in units of standard deviation, and therefore they are readily comparable across features and also across regions and datasets. Despite our initial pre-filtering, not all of the remaining time-series features could be extracted successfully from all our datasets. We therefore performed an additional post-filtering. Features that failed to be extracted were excluded. To ensure consistency, features were only included in the final analysis if they could be included for each dataset. Features that failed to produce a finite effect size for a region, had their effect size for that region set to zero. Following pre-filtering and post-filtering, a total of 6 827 dynamical features were retained across datasets.

To guide and aid interpretation, we also stratified dynamical features into 10 broad categories, based on associated keywords [29, 30] and further inspection (Table S2). These include: (1) distribution (e.g. mean, variance, outliers, and tests for distribution family); (2) autocorrelation (and its nonlinear and lagged versions); (3) periodic (e.g. power spectrum, seasonality, wavelet); (4) entropy and related quantities; (5) symbolic (discretisation of the signal and resulting motifs and transitions between them); (6) forecasting and model fit from past to future; (7) transformations of the data (e.g. scaling, surrogates); (8) variability in time (e.g., change-points, stationarity); (9) measures from the complex systems literature (e.g., time-reversal, fractality, embeddings); and (10) miscellaneous others (e.g., visibility graph). Note that this is not a formal taxonomy: some features may plausibly belong to more than one category, and some categories are only loosely defined. Rather, the goal is to provide a broad overview.

#### Dynamical profile similarity

To quantify the similarity between the temporal profiles of each pair of brain regions (neurons in the nematode), each dynamical feature is z-scored across regions. The vectors of z-scored regional features are then correlated for each pair of brain regions, producing a matrix of ‘dynamical profile similarity’ (also termed ‘temporal profile similarity’ [31]) that represents the strength of the similarity of the local dynamical fingerprints of brain areas. This procedure is performed separately within each individual and condition of each dataset.

The coupling between DPS and functional connectivity is in turn obtained by correlating the vectorised DPS and FC matrices. FC is computed as the zero-lag correlation between pairs of regional time-series.

#### catch22 subset of representative features

Among all hctsa dynamical features, Lubba and colleagues [67] identified a reduced set of 22 features (23 after adding standard deviation) that captures a diverse range of interpretable time-series properties from the broader literature on dynamical systems (S3). The features in this subset (known as the CAnonical Time-series CHaracteristics or catch22) were identified as a way of reducing the dimensionality of the full hctsa feature-set to simplify computation and interpretation. The catch22 set consists of time-series features that (i) exhibit strong classification performance across 93 natural and artificial time-series classification datasets, comparable to the performance of the full hctsa set; and (ii) are minimally redundant on those same datasets [67]. We use this reduced set for our multivariate association analysis with regional gene expression. Note that one of these 23 features, *CO_HistogramAMI_even_2_5*, was excluded by our pre-filtering and therefore is not included among the features that we extract. A second feature, ‘acf timescale’ (first1e_acf_tau), occasionally failed to produce finite effect sizes for every region of every dataset due to no variance, and therefore had to be excluded from the PLS analysis, leaving 21 for the final analysis.

### Cortical gene expression

#### Human brain gene expression from microarray

Regional human gene expression profiles were obtained using microarray data from the Allen Human Brain Atlas (AHBA) [56], with preprocessing as recently described [191]. The Allen Human Brain Atlas (AHBA) is a publicly available transcriptional atlas containing gene expression data measured with DNA microarrays and sampled from hundreds of histologically validated neuroanatomical structures across normal postmortem human brains from six donors (five male and one female; age 24–55 years). We extracted and mapped gene expression data to the 100 cortical ROIs of the Schaefer parcellation using the abagen toolbox https://abagen.readthedocs.io/ [192]. Data were pooled between homologous cortical regions to ensure adequate coverage of both left (data from six donors) and right hemisphere (data from two donors). Distances between samples were evaluated on the cortical surface with a 2mm distance threshold. Only probes where expression measures were above a background threshold in more than 50% of samples were selected. A representative probe for a gene was selected based on highest intensity. Gene expression data were normalised across the cortex using scaled, outlier-robust sigmoid normalisation. 15, 633 genes survived these preprocessing and quality assurance steps. We also replicate the main results using human gene expression data from an alternative modality, RNA-seq, which was available from 2/6 AHBA donors [56].

#### Macaque cortical gene expression from stereo-seq

We used cortex-wide macaque gene expression data recently made available by [57], who combined single-nucleus RNA sequencing (“snRNA-seq”) with high-resolution, large-field-of view spatial transcriptomics from spatiotemporal enhanced resolution omics-sequencing (“stereo-seq”) [57]. Specifically, the authors made available (https://macaque.digital-brain.cn/ spatial-omics) post-mortem gene expression data covering 143 regions of the left cortical hemisphere of one 6yo male cynomolgus macaque (*Macaca fascicularis*). We refer the reader to [57] for details. The animal protocol was approved by the Biomedical Research Ethics Committee of CAS Center for Excellence in Brain Science and Intelligence Technology, Chinese Academy of Sciences (ION-2019011). Animal care complied with the guideline of this committee [57].

Briefly, Chen and colleagues obtained 119 coronal sections at 500-*µ*m spacing, covering the entire cortex of the left hemisphere, which were used for stereo-seq transcriptomics [57]. Adjacent 50-*µ*m thick sections were also acquired for regional microdissection and snRNAseq analysis, as well as 10-*µ*m sections adjacent to each stereo-seq section, which were used for the anatomical parcellation of brain regions via immunostaining [57]. Stereo-seq is a DNA nanoball (DNB) barcoded solid-phase RNA capture method [57]. It involves reverse transcription of RNAs released from frozen tissue sections fixated onto the stereo-seq chip, and subsequent PCR amplification. The resulting “amplified-barcoded complementary DNA (cDNA) is used as template for library preparation, and sequenced” to obtain high-resolution spatially resolved transcriptomics [57].

Gene expression data were made available for 143 cortical regions of the left hemisphere, including prefrontal, frontal, cingulate, somatosensory, insular, auditory, temporal, parietal, occipital and piriform areas. As reported in [57], for each coronal section, the cortical region and layer parcellation were manually delineated on Stereoseq data background, based on cytoarchitectual pattern (e.g. cell density, cell size) revealed by total mRNA expression, nucleic acid staining, and NeuN staing of adjacent sections. To make the gene expression data compatible with our macaque functional MRI datasets, the aggregated gene expression across layers was manually mapped onto the cortical regions of the “regional mapping” macaque atlas of Kötter and Wanke [144], mirroring data between hemispheres [61].

#### Mouse brain gene expression from in situ hybridization

Mouse gene expression profiles were obtained using in situ hybridization data from the Allen Mouse Brain Atlas [58]. We followed the same preprocessing as recently described [193]. Briefly, the Allen Mouse Brain Atlas consists of data acquired from a pipeline that includes semi-automated riboprobe generation, tissue preparation and sectioning, in-situ hybridization (ISH), imaging, and data post-processing. These data were acquired both sagittally and (for a smaller set of genes) coronally, and were further processed, aligned by the Allen Institute to their Common Coordinate Framework version 3 (CCFv3) reference atlas [194] and summarized voxelwise through a measure termed gene expression energy (defined as the sum of expressing pixel intensity divided by the sum of all pixels in a division), resulting in 3D gene expression images at a 200 *µ*m isotropic resolution. This gene expression energy increases in regions of high expression, and is bounded by zero in regions of no expression. For our study, we used gene expression energy data from the coronal dataset (4345 gene expression images corresponding to 4082 unique genes), because of its whole-brain coverage and data quality. Fernandes and colleagues [195] provide tools to work with this gene expression data; these tools are available online (github.com/DJFernandes/ABIgeneRMINC). Voxelwise gene expression data were further summarized as normalized mean expression within regions of interest as defined by the CCFv3 reference atlas. For each ROI and each gene, voxelwise expression energy data were averaged over voxels containing valid expression signal. Finally, independently for each region, gene expression data were normalized by regressing out mean gene expression across the brain.

We also replicate the main results using mouse gene expression data from two recently released alternative databases from MERFISH, as reported in [80] and [81]. Briefly, the database from [80] is a whole mouse brain spatial transcriptomics dataset containing 3.9 million segmented cells passing quality control (MERFISH-C57BL6J-638850). It was acquired using multiplexed error-robust fluorescence in situ hybridization (MERFISH) technique with a 500 gene panel. The MERFISH data was registered to the Allen CCFv3 standard template, acquired CCF coordinates for each segmented cell, and assigned into pre-defined brain structures. A separate single-cell RNA sequencing (scRNA-seq) dataset was acquired using 10Xv3 technique and imputed (projected) into MERFISH space, deriving 8,460 marker genes for each MERFISH cell. A region-by-gene matrix was derived from the cell-by-gene tabular data by aggregating within the assigned Allen CCFv3 atlas regions and taking the average hierarchically following a pre-defined simplified anatomical hierarchy. The [81] database is a whole mouse brain spatial transcriptomics dataset acquired using MERFISH technique with a 1122 gene panel, containing 9.3 million segmented cells passing quality control (Zhuang-ABCA-1/2/3/4). The MERFISH data was registered to the Allen CCFv3 standard template, and coordinates were acquired for 5.4 million cells, which were then assigned into pre-defined brain structures. We derived the region-by-gene matrix similarly as above. Out of 23 brain-related genes, 11 are available in the Yao database, and 13 are available in the Zhang database.

#### Marmoset brain gene expression from in situ hybridization

Gene expression profiles for the common marmoset (*Callithrix jacchus*) were also obtained from the Marmoset Gene Atlas, a recent database of in situ hybridization transcriptomics across the entire marmoset brain (https://gene-atlas.bminds.brain.riken.jp/) available within the Brain/MINDs data portal (https://dataportal.brainminds.jp/). Full details are provided in the original publications [59, 60]. All procedures were performed in accordance with a protocol approved by RIKEN Institutional Animal Care [59, 60]. Briefly, brains were sectioned on a freezing microtome at 28 *µ*m. ISH staining for each probe consisted of 60 coronal sections evenly spaced at 196 *µ*m intervals. Whole section images were scanned and digitised. Converted imaged were aligned with Nissl stained reference atlas. The Marmoset Gene Atlas database provides gene expression data for marmosets from newborn to 1 year old. The available age of each gene expression data varies by gene; transcriptomic data from 1-year-old animals were used when available. For genes lacking 1-year-old data (*HCN1, HTR1A, ADRA1A, ADRA2A, CHRNB2, GRIN3A, CALB1, GABRA1, OXTR, CNR1, OPRK1, VIP, GRM1, GRM5, MOBP*) the available neonatal brain data were used instead. This database allows investigation of gene expression levels using anatomical location information referenced from HE-stained histological images. Although each ISH section is complemented by an adjacent neuroanatomically stained section, enabling users to compare gene expression against a brain atlas, mapping gene expression patterns over the entire brain is both time and labor intensive. From the database, 300×300 pixel images (approximately 750 x 750 *µ*m) were extracted using ImageJ (https://imagej.net/ij/) at 3-5 locations per region across 70 cortical regions from the marmoset MBM-vM atlas [180] corresponding to our MRI data. Since each gene expresses in different layers of cortex, images from the layer showing the highest gene expression were selected [60]. The extracted images were converted to grayscale, and the pixel values for all images were obtained for each gene. The bottom 1 percentile of the overall values for each gene was used as the threshold for binarization of the image. The number of black pixels was calculated for each binarized image, representing the expression level for that image. The expression levels of the images extracted for each region were averaged, and this value was used as the expression level for that region.

#### Macaque parvalbumin density from immunohistochemistry

Burt and colleagues [82] assembled data on the immunohistochemically measured densities of parvalbumin-expressing inhibitory interneurons for several macaque brain areas, from multiple immunohistochemistry studies [196–199]. We used these data as provided by [61], who mapped parvalbumin density onto the Regional Mapping macaque atlas used in the present study.

### Biophysical computational model

We provide here a compact synthesis of the multiscale biophysical AdEx whole-brain model from [76]: we refer the reader to the original publication for full details of the implementation. For the present work, all parameters were kept the same as in Sacha et al. [76]. Note that only the BOLD signals from the final whole-brain scale realised through the integration in The Virtual Brain simulator [200, 201]. were used in the present analysis. However, for the purpose of illustrating our modelling framework, we briefly introduce each of the multi-scale components that were developed and analysed in detail by Sacha et al. [76], progressing from single-neuron dynamics to mesoscale mean-field formulations and largescale whole-brain models.

At the microscopic level, neuronal activity is described using the adaptive exponential integrate-and-fire (AdEx) model [76]. Synaptic currents arise from excitatory and inhibitory conductances defined by quantal synaptic events, while adaptation is controlled by voltage-dependent and spike-triggered mechanisms. Mesoscopically, a Master Equation formalism yields population-averaged firing rates, covariances, and adaptation variables. The population transfer function maps the statistics of membrane potential fluctuations to output firing rates through a semi-analytic expression involving mean, variance, and correlation timescales of voltage dynamics. A first-order reduction leads to a compact system for excitatory and inhibitory mean-field populations driven by external input composed of deterministic and Ornstein– Uhlenbeck noise terms, representing the activity of an individual brain region [76]. At the whole-brain scale, multiple mean-field nodes (corresponding to the 68 regions of the Desikan-Killiany anatomical parcellation [202], as in [76]) are coupled via anatomically defined connectivity from diffusion MRI tractography and associated propagation delays. Here we used the same anatomical data, atlas, and parameters as in [76]. Finally, regional BOLD signals are generated through a Balloon– Windkessel hemodynamic model [203]. Below, equations are followed by a short explanatory note. For the present work, all parameters were kept the same as in Sacha et al. [76].

#### Single-cell AdEx model

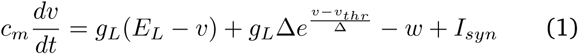

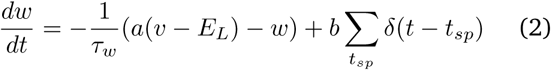

These equations describe the membrane voltage and adaptation current of an AdEx neuron, driven by leak, exponential spike-initiation currents, synaptic input, and spike-triggered adaptation.

#### Synaptic conductance

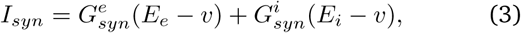

This expresses synaptic current as the sum of excitatory and inhibitory conductance-driven components.

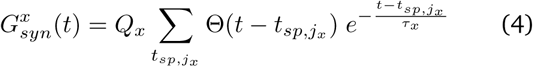

Conductances follow exponential decays triggered by presynaptic spikes of type *x* ∈ {*e*, *i*}.

#### Mean-field equations

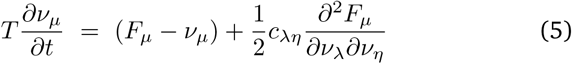

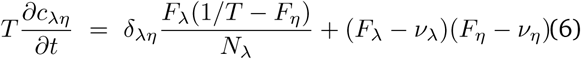

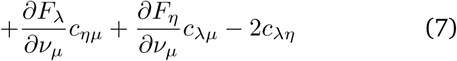

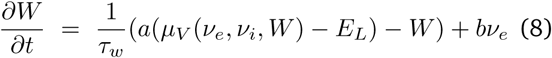

These equations evolve the mean firing rates, covariances, and population adaptation based on the transfer function *F*_μ_.

#### Transfer function

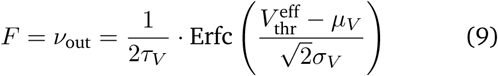

This semi-analytic expression converts voltage statistics to output firing rate.

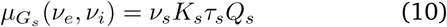

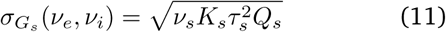

These formulas give mean and variance of synaptic conductances assuming Poisson presynaptic firing.

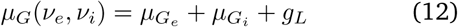

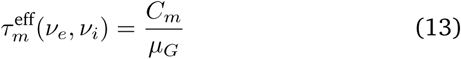

These define total conductance and effective membrane time constant.

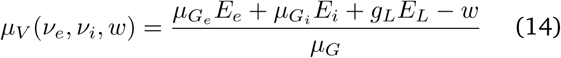

This is the mean membrane potential under fluctuating conductances.

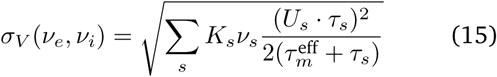

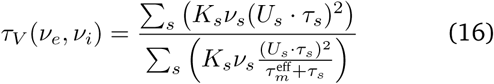

where *s* = {*e*, *i*} and U*_s_* = *Q_s_*μ*_G_*(*E_s_* − μ*_V_*). These give the variance and correlation time of membrane voltage fluctuations.

The 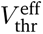 in Eq. 9 is the phenomenological spike threshold voltage (see [204] for details).

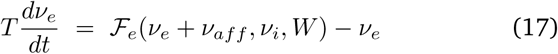

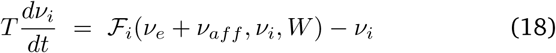

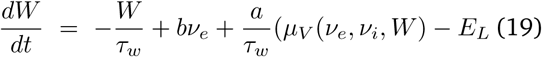

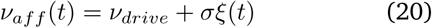

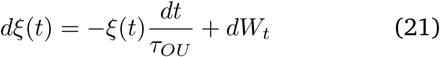

These define the population dynamics under noisy external input.

#### Whole-brain coupling

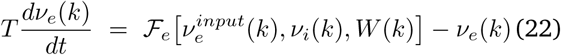

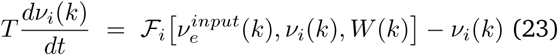

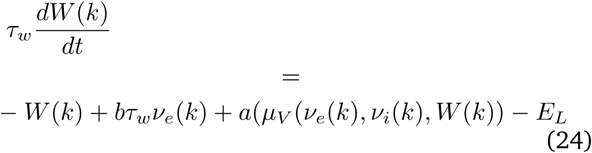

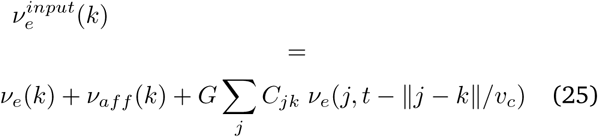

These equations specify the long-range excitatory coupling through structural connectivity, scaled by global coupling *G* (here kept fixed as in [76]) and delayed by propagation over tract lengths.

#### Simulated BOLD signals

The simulated BOLD signal was generated through the dedicated *BOLD* monitor provided by The Virtual Brain software. The simulated mean field time series of the excitatory population (down-sampled at 250Hz) of each node were convolved with the hemodynamic response function (described by the first order Volterra kernel of the Ballon Windekessel model [205]). This signal was finally sampled to the fMRI repetition time (TR) of 2s, as in [76].

### Statistical analyses

#### Anaesthetic effect size

For each contrast (awake vs anaesthesia) in each dataset, we consider each region/neuron and feature in turn. We use the measures of effect size toolbox for MATLAB https://github.com/hhentschke/ measures-of-effect-size-toolbox [206] to estimate an effect size for the difference between that feature’s values during wakefulness and during anaesthesia using Hedge’s measure of the standardized mean difference, *g*, which is interpreted in the same way as Cohen’s *d*, but more appropriate for small sample sizes [47]. Note that features were not normalised across regions before analysis, as doing so could obscure differences in magnitude. Although hctsa features span several orders of magnitude, such that raw differences cannot be meaningfully compared across features, effect sizes are expressed in units of standard deviation, and therefore they are readily comparable across features and also across regions and datasets. For each contrast in each dataset, this procedure produces a matrix of awake-vs-anaesthesia effect sizes, with dimensions *n* regions/neurons × *f* features (with *f* = 6827 for all species after post-filtering).

#### Significance of the number of consistent features

To establish whether the empirically observed number of consistent dynamical features is significantly greater than the number of features that would be expected to exhibit consistency (all increases or all decreases) across all contrasts, just by chance alone, we construct a null distribution with the same number of features and contrasts (here, 15) where the sign of each feature and contrast is assigned at random (+1 or -1). We then compute the number of features having the same sign across all 15 contrasts. Repeating this process 10,000 times produces a null distribution against which we test the empirical number of consistent features to obtain a p-value for our null hypothesis.

#### Multivariate association with Partial Least Squares

Partial Least Squares (PLS) analysis was used to relate regional gene expression to anaesthetic-induced changes in local dynamics in a multivariate fashion. PLS analysis is an unsupervised multivariate statistical technique that decomposes relationships between two datasets *X_n_*_×_*_g_* and *Y_n_*_×_*_f_* into orthogonal sets of latent variables with maximum covariance, which are linear combinations of the original data [63, 64]. In the present case, *X_n_*_×_*_g_* is regional gene expression across *n* regions (100 for the human; 82 for the macaque; 70 for the marmoset; and 72 for the mouse) and *g* genes (the same set of 23 genes for each species). *Y_n_*_×_*_f_* is the matrix of regional anaesthetic-induced changes in dynamical features, across *f* features.

To ensure consistent gene-dynamics relationships across species, we perform this analysis on matrices *X_n_*_×_*_g_* and *Y_n_*_×_*_f_* obtained by vertically concatenating the species-specific matrices for human, macaque, and mouse. Within each species, we separately z-score the *X_n_*_×_*_g_* and *Y_n_*_×_*_f_* matrices column-wise. We then concatenate vertically the species-specific *X* matrices of z-scores, and we likewise concatenate vertically the species-specific *Y* matrices of z-scores. This is possible because the columns are the same for each across all three species: the same 23 genes, and the same dynamical features. This results in a single *X_n_*_×_*_g_* matrix and a single *Y_n_*_×_*_f_* matrix, each with *n* = 100+82+70+72 = 324 rows.

PLS finds components from the predictor variables (regional gene expression) that have maximum covariance with the response variables (regional changes in dynamical features). The PLS components (i.e., linear combinations of the weighted variables) are ranked by the covariance between predictor and response variables so that the first few PLS components provide a low-dimensional representation of the covariance between the higher-dimensional data matrices. Concretely, this is achieved by performing singular value decomposition (SVD) on the matrix *Y* ^′^*X*, such that:

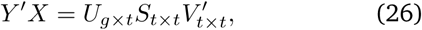

where *U_g_*_×_*_t_* and *V_t_*_×_*_t_* are orthonormal matrices consisting of left and right singular vectors, and *S_t_*_×_*_t_* is a diagonal matrix of singular values. The *i*th columns of *U* and *V* constitute a latent variable, and the *i*th singular value in *S* represents the covariance between singular vectors. The *i*th singular value is proportional to the amount of covariance between gene expression and anaesthetic-induced dynamical changes captured by the *i*th latent variable, where the effect size can be estimated as the ratio of the squared singular value to the sum of all squared singular values.

For the main analysis, we use the subset of *catch22* representative features (excluding *acf_timescale* which did not return finite values for all regions; and *ami2* which did not pass our pre-filtering step, leaving the same 21 dynamical features available for use in all species). As a further validation, we also repeat the analysis using all 485 consistent features. Significance of the latent variables is assessed against a null distribution of null maps with preserved spatial autocorrelation, as described below.

#### Null model

The statistical significance of the covariance explained by each PLS model is tested by permuting the response variables 10 000 times while considering the spatial dependency of the data by using spatial autocorrelation-preserving permutation tests, to control for the spatial autocorrelation inherent in neuroimaging data, which can induce an inflated rate of false positives [65, 207].

For each species, we generate null maps using Moran spectral randomisation based on on the inverse Euclidean distances between parcel centroids for that species, as implemented in the BrainSpace toolbox (https://brainspace.readthedocs.io/en/latest/) [66]. Moran spectral randomisation quantifies the spatial autocorrelation in the data in terms of Moran’s *I* coefficient, by computing spatial eigenvectors known as Moran eigenvector maps. The Moran eigenvectors are then used to generate null maps by imposing the spatial structure of the empirical data on randomised surrogate data [66, 207]. As for the empirical data, each null map is z-scored within-species and the corresponding null maps are then concatenated vertically across species, to obtain a null *Y* matrix. PLS analysis is then applied to the original *X* matrix and this null *Y* matrix. This procedure is repeated 10 000 times, to obtain a null distribution of the singular values associated with each latent variable. This test embodies the null hypothesis that regional gene expression and regional dynamical changes induced by anaesthesia are spatially correlated with each other only because of inherent spatial autocorrelation. The p-value is computed as the proportion of null singular values that are greater in magnitude than the empirical singular values. Thus, these p-values represent the probability that the observed spatial correspondence between genes and dynamical features could occur by randomly correlating maps with comparable spatial autocorrelation.

## Data and code availability

The original pharmacological fMRI data are available from the corresponding authors of the original publications referenced herein. Human gene expression data from the Allen Human Brain Atlas [56] are available at https://human.brain-map.org/. Macaque cortical gene expression data from [57] are available at https://macaque.digital-brain.cn/spatial-omics. The dataset is provided by Brain Science Data Center, Chinese Academy of Sciences (https://braindatacenter.cn/). Mouse gene expression data from in situ hybridization are available from the Allen Mouse Brain Atlas [58] at https://mouse.brain-map.org/. The Marmoset Gene Atlas is available at https://gene-atlas.bminds. brain.riken.jp/. The MERFISH mouse transcriptomic data are available through the Neuroscience Multi-omic Data Archive (NeMO, https://nemoarchive.org/). Human Connectome Project data in DSI Studio-compatible format are available at http://brain.labsolver.org/ diffusion-mri-templates/hcp-842-hcp-1021.

The Highly Comparative Time-Series Analysis (hctsa) toolbox is freely available at https://github.com/ benfulcher/hctsa. The abagen toolbox for processing of the AHBA human transcriptomic dataset is available at https://abagen.readthedocs.io/. The BrainSpace toolbox for Moran spectral randomisation is available at https://brainspace.readthedocs. io/en/latest/. The Measures of Effect Size toolbox is available at https://github.com/hhentschke/ measures-of-effect-size-toolbox. The Virtual Brain code for the biophysical model is available at https://www.thevirtualbrain.org/tvb/zwei/home(version 2.9).

## Author contributions

A.I.L., B.M. conceived the analysis; A.I.L. carried out analysis and visualisation; L.U., J.T., and B.J. designed the macaque experiments and collected and curated the macaque data; A.L., P.D., and P.D.K. designed the zebrafish experiments; A.L. collected and curated the zebrafish data; KM performed regional quantification of marmoset gene expression K.M., J.H., H.O. designed the marmoset experiments and collected and curated the marmoset data; D.G., A.R., R.I., D.J., designed the human experiments and collected and curated the human data; S.G. and A.Go. designed the mouse experiments and collected and curated the mouse data, and provided the mouse connectome; C.W.C. designed the nematode experiments and collected and curated the nematode data; C.M.S. and R.C. contributed to macaque data processing; G.S., Z-Q. L. contributed to data analysis; Y.Y., Z-Q. L. contributed mouse gene expression data; B.F., G.S., A.D., D.K.M., A.Go., B.J., E.A.S. contributed to interpretation; B.J., B.M. supervised the project; A.I.L. and B.M. wrote the manuscript with feedback from all co-authors. All authors approved the manuscript.

## Funding

AIL acknowledges the support of St John’s College, Cambridge; the Natural Sciences and Engineering Research Council of Canada (NSERC), [funding reference number 202209BPF-489453-401636, Banting Postdoctoral Fellowship] and FRQNT Strategic Clusters Program (2020-RS4-265502 - Centre UNIQUE - Union Neuroscience & Artificial Intelligence - Quebec) via the UNIQUE Neuro-AI Excellence Award; and a Wellcome Early Career Award (grant number 226924/Z/23/Z). BM acknowledges support from the Natural Sciences and Engineering Research Council of Canada (NSERC), Canadian Institutes of Health Research (CIHR), Brain Canada Foundation Future Leaders Fund, the Canada Research Chairs Program, the Michael J. Fox Foundation, and the Healthy Brains for Healthy Lives initiative. ZQL acknowledges support from the Fonds de Recherche du Québec – Nature et Technologies (FRQNT). GS was supported by a postdoctoral fellowship from the Canadian Institutes of Health Research (CIHR). CWC is supported by NIH grant R35 GM145319 - "Pan-neuronal functional imaging and anesthesia". This work was also supported by the visitor program of the European Institute for Theoretical Neuroscience [to A.I.L.]; the Canadian Institute for Advanced Research (CIFAR; grant RCZB/072 RG93193) [to DKM and EAS]; Cambridge Biomedical Research Centre and NIHR Senior Investigator Awards and the British Oxygen Professorship of the Royal College of Anaesthetists [to DKM]; the European Research Council (ERC) under the European Union’s Horizon 2020 research and innovation program (DISCONN; no. 802371 to A.Go.; and no. 101125054 - BRAINAMICS to A.Go.); the Stephen Erskine Fellowship of Queens College Cambridge [to EAS]. RC and AD were supported by CNRS and the European Union (Human Brain Project, H2020-945539). This work was also supported by the Fondation Bettencourt Schueller (to B.J.); Fondation de France (to B.J.), Human Brain Project (Corticity project FLAG-ERA JTC2017, to B.J.); Institut National de la Sante et de la Recherche Medicale (to B.J.), UVSQ (to B.J.), Commissariat a l’Energie Atomique (to B.J.), College de France (to B.J.); the Fondation pour la Recherche Medicale (FRM grant number ECO20160736100, to J.T.); FNRS Belgium, project MIS/VA - F.4512.21 (to C.M.S.) and grant Embodied-Time – 40011405 (to C.M.S.). YY acknowledges the support of the Quebec Bio-Imaging Network Postdoctoral Recruitment Scholarship, and David T.W. Lin Fellowship from the McGill University Faculty of Medicine and Health Sciences. This work was also supported by the program for Brain Mapping by Integrated Neurotechnologies for Disease Studies (Brain/MINDS) from the Japan Agency for Medical Research and Development (AMED) (Grant Number JP23wm0625001 to HO), JSPS KAKENHI (Grant Number JP20H03630 to JH), and by “MRI platform” as a program of Project for Promoting public Utilization of Advanced Research Infrastructure of the Ministry of Education, Culture, Sports, Science and Technology (MEXT), Japan (Grant Number JPMXS0450400622 to JH). A.L. was supported by PhD scholarships from the Natural Sciences and Engineering Research Council of Canada (NSERC), Fonds de recherche du Québec–Nature et technologies (FRQNT), and Unifying Neuroscience and Artificial Intelligence–Québec (UNIQUE). The zebrafish work was funded by the NSERC (RGPIN-2019-06887 and RGPIN-2024-06492 to P.D. and RGPIN-2023-05980 to P.D.K.), the Sentinel North program of Université Laval (Canada First Research Excellence Fund), the Northern Contaminants Program of Canada, and the Next Generation Networks for Neuroscience (Neuronex)/FRQS (295823). Acquisition of the human sevoflurane dataset was supported from institutional funds by the Technical University of Munich. For the purpose of open access, the authors have applied a Creative Commons Attribution (CC BY) licence to any Author Accepted Manuscript version arising from this submission. Any opinions, findings, and conclusions or recommendations expressed in this material are those of the authors and do not reflect the views of the funders.

## Competing interests

The authors declare no competing interests.

Supplementary Figures

## Notes

### Competing Interest Statement

The authors have declared no competing interest.

### Summary of Updates

Added zebrafish calcium imaging and marmoset gene expression; new biophysical model

