## Supplementary Information for "Comprehensive profiling of anaesthetised brain dynamics across phylogeny"

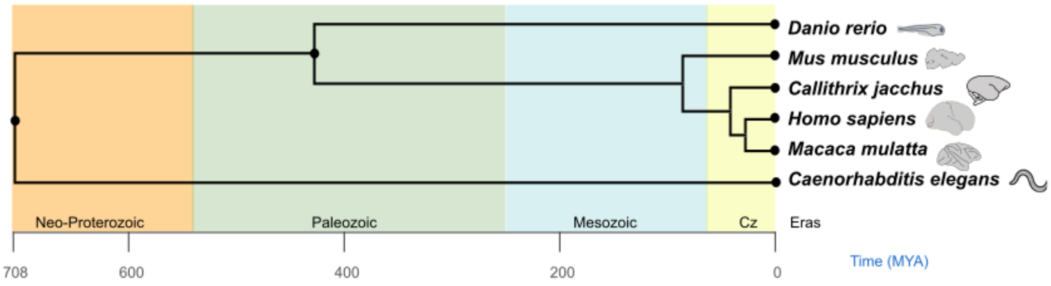


Figure S1. **Phylogenetic tree for the six species included in the present study |** The tree originates (far left) from a common ancestral organism >700 million years ago (MYA). Geological eras are also shown. Figure generated using `TimeTree` (https://timetree.org/) [34].

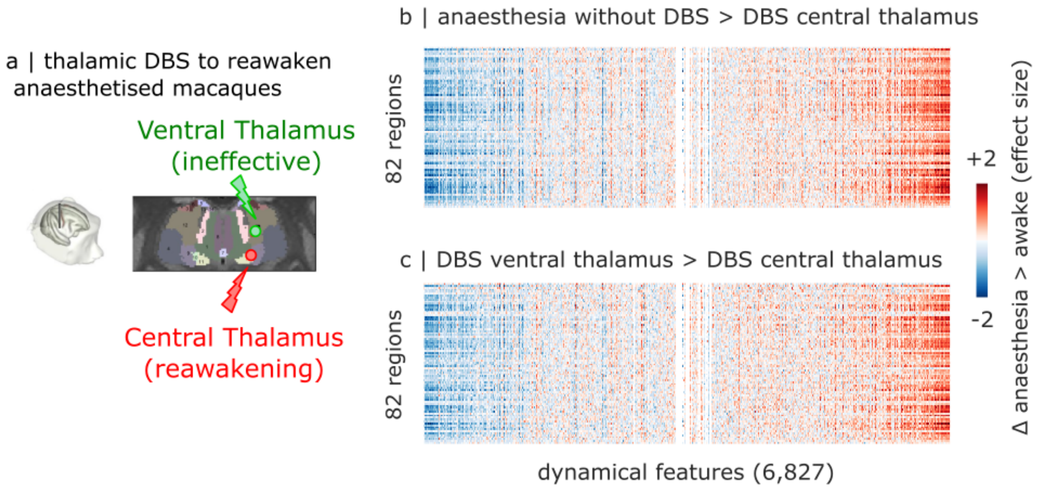


Figure S2. **DBS-induced restoration of neural dynamics in the anaesthetised macaque |** (**a**) Illustration of deep-brain stimulation targets in the macaque thalamus: centromedian thalamus, and ventral thalamus (adapted from Figure 1 of [43], published under CC-BY license). (**b**) Reawakening from anaesthesia induced by CT DBS, versus propofol anaesthesia without DBS. (**c**) Reawakening from anaesthesia induced by CT DBS, vs VT DBS at the same intensity (which does not reawaken the animal). Both effects resemble the difference between anaesthesia and baseline wakefulness, even though here wakefulness is achieved through stimulation while propofol is still present. For visualisation purposes, the color range is capped at [-2, 2].

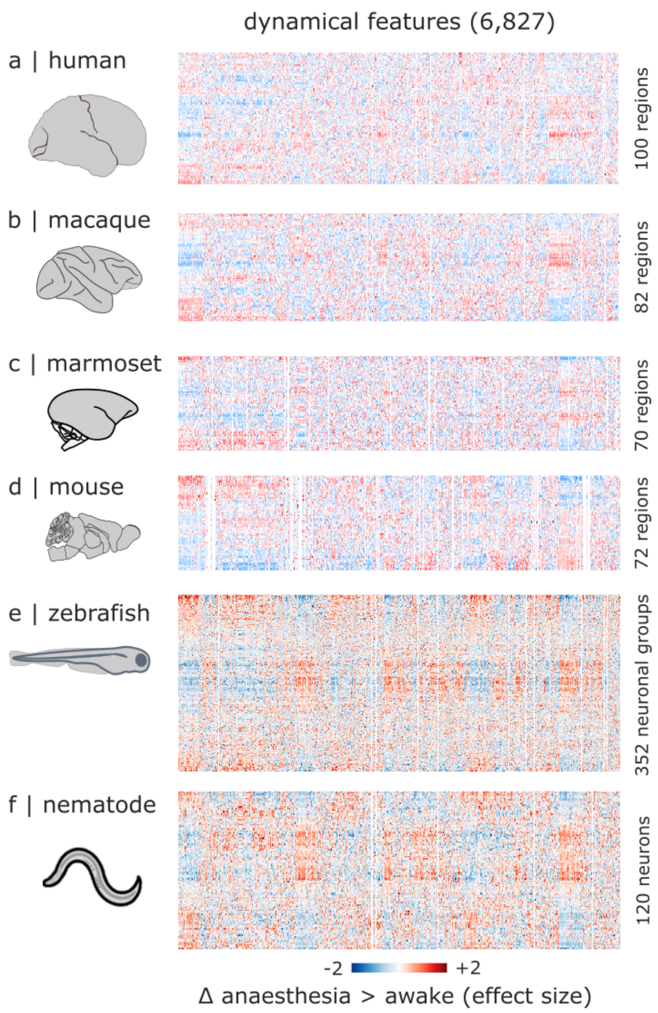


Figure S3. **Z-scored anaesthetic-induced changes in neural dynamics across species |** (**a**) Human: z-scored mean effect sizes across awake vs vol 3% sevoflurane, and recovery vs vol 3% sevoflurane. (**b**) Macaque: z-scored mean effect sizes across awake vs sevoflurane; awake vs propofol; awake vs ketamine (for the Multi-anaesthesia dataset); awake vs propofol (no DBS); CT DBS versus propofol; and CT DBS vs VT DBS. (**c**) Marmoset: z-scored mean effect sizes across awake vs sevoflurane; awake vs propofol; and awake vs isoflurane. (**d**) Mouse: z-scored mean effect sizes across awake vs halothane; and awake vs medetomidine-isoflurane. (**e**) Larval zebrafish: z-scored effect size for awake vs tricaine. (**f**) Nematode: z-scored effect size for awake vs isoflurane. Z-scoring is performed separately for each column (feature) in each species. Rows are then reordered to highlight consistent patterns of regional variation. Features are provided in the same order across species.

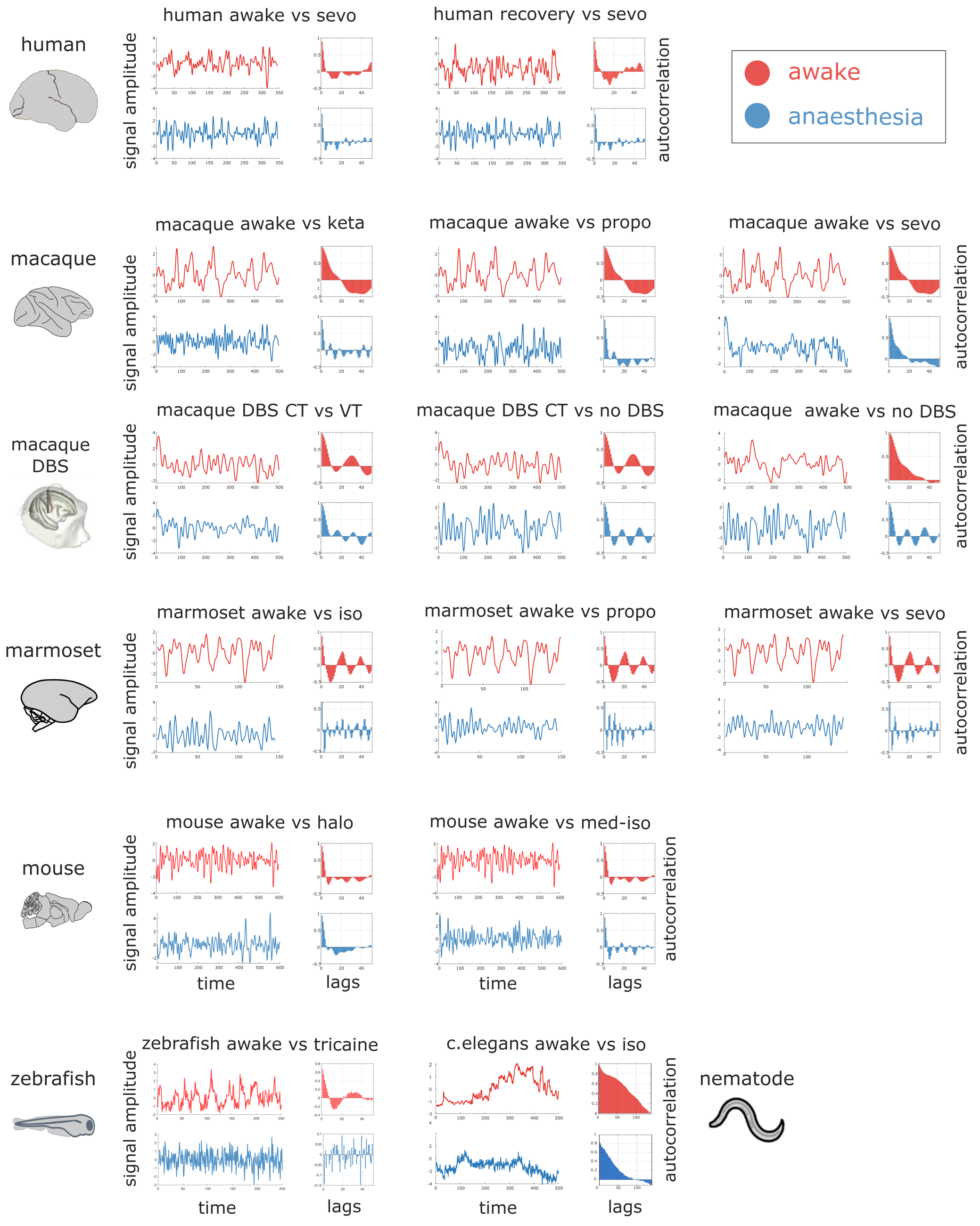


Figure S4. **Examples of anaesthetic effect on neural time-series across species |** Example time-series and their autocorrelation function (up to lag-50) during wakefulness/recovery (top) and anaesthesia (bottom), for each contrast. Red and blue indicate wakefulness and anaesthesia, respectively.

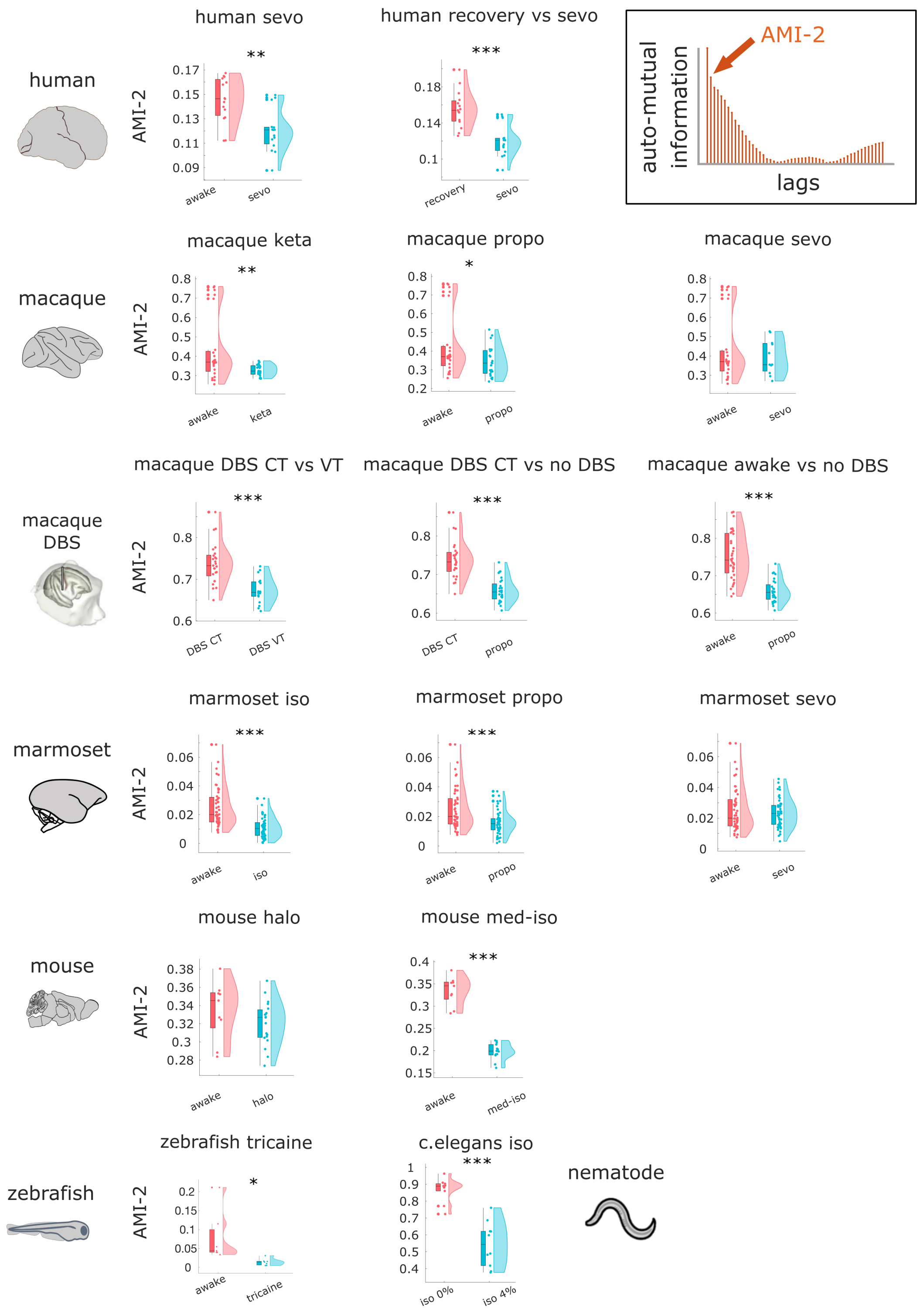


Figure S5. **Individual-level anaesthetic effect on lag-2 automutual information |** Ordinate represents the brain's mean value of lag-2 automutual information, across all regions. Each data-point represents one scan. Box-plots: center line, median; box limits, upper and lower quartiles; whiskers, $1.5\times$ interquartile range. *, $p < 0.05$; **, $p < 0.01$; ***, $p < 0.001$, from non-parametric permutation-based t-test (dependent samples for human, marmoset and zebrafish; independent samples for macaque, mouse and nematode).

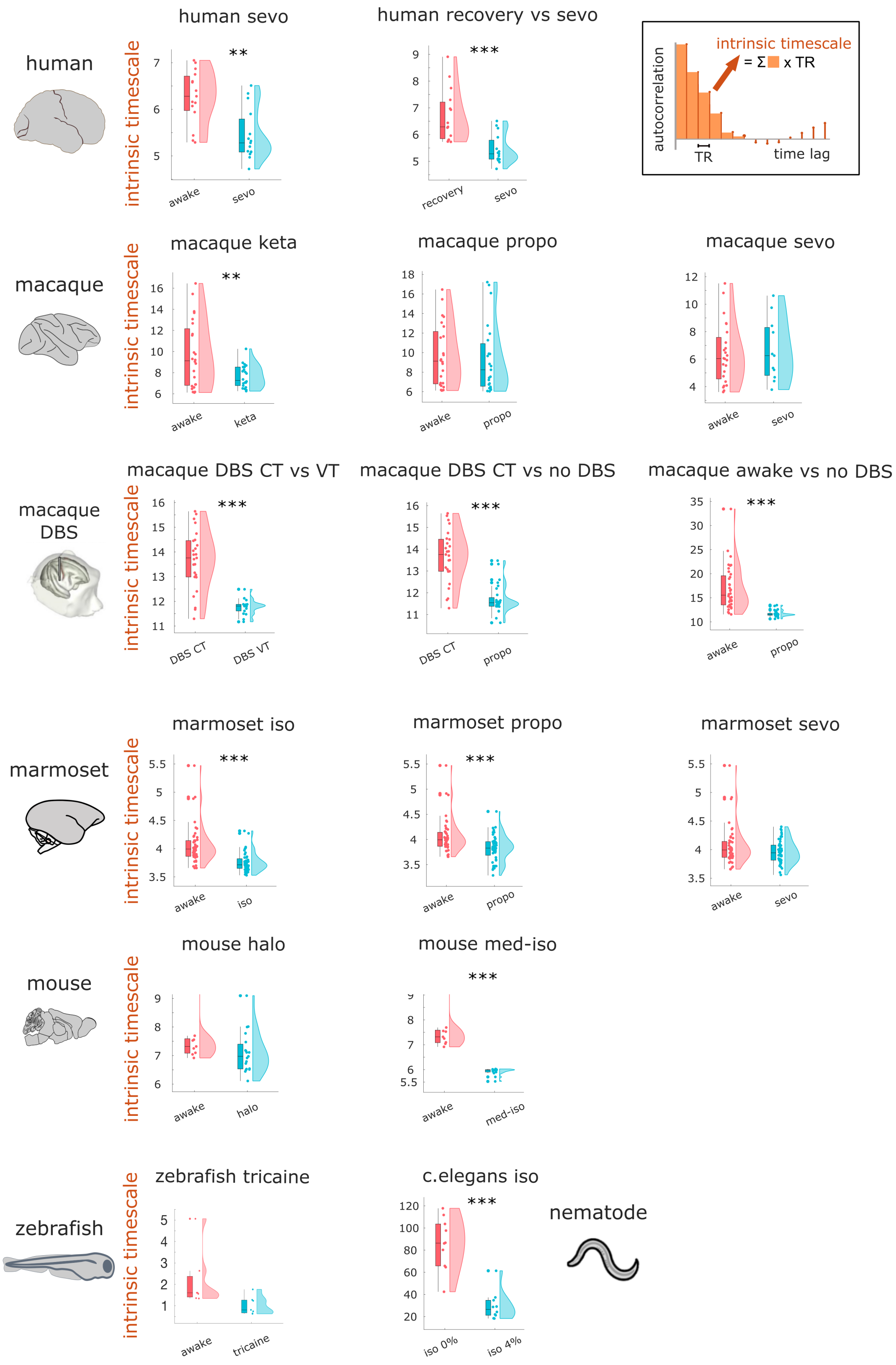


Figure S6. **Individual-level anaesthetic effect on brain-wide intrinsic neural timescales across species |** Ordinate represents the brain's mean value of intrinsic neural timescale, across all regions. Each data-point represents one scan. Box-plots: center line, median; box limits, upper and lower quartiles; whiskers, 1.5× interquartile range. *, $p < 0.05$; **, $p < 0.01$; ***, $p < 0.001$, from non-parametric permutation-based t-test (dependent samples for human, marmoset and zebrafish; ; independent samples for macaque, mouse and nematode).

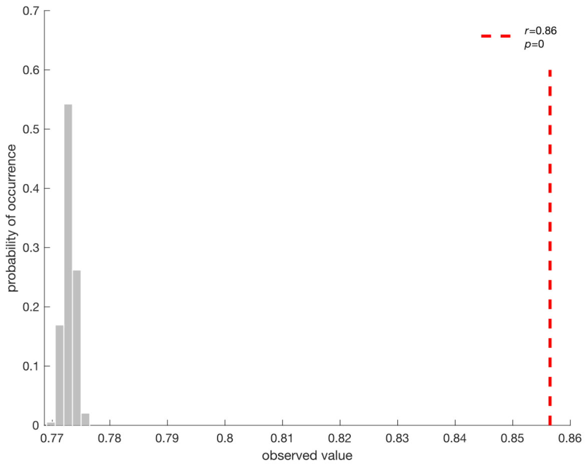


Figure S7. **Similarity of anaesthetic-induced dynamical changes is significant beyond the effect of shared sign |** The empirically observed mean correlation between the consistent profiles of anaesthetic-induced contrasts is significantly greater than the level of correlation that would be expected by chance, purely based on the fact that features have the same sign ($p < 0.0001$). We quantify this with a null distribution of 10,000 surrogate datasets, generated by shuffling the consistent features within each contrast while preserving their sign (i.e., positive and negative features are reshuffled separately).

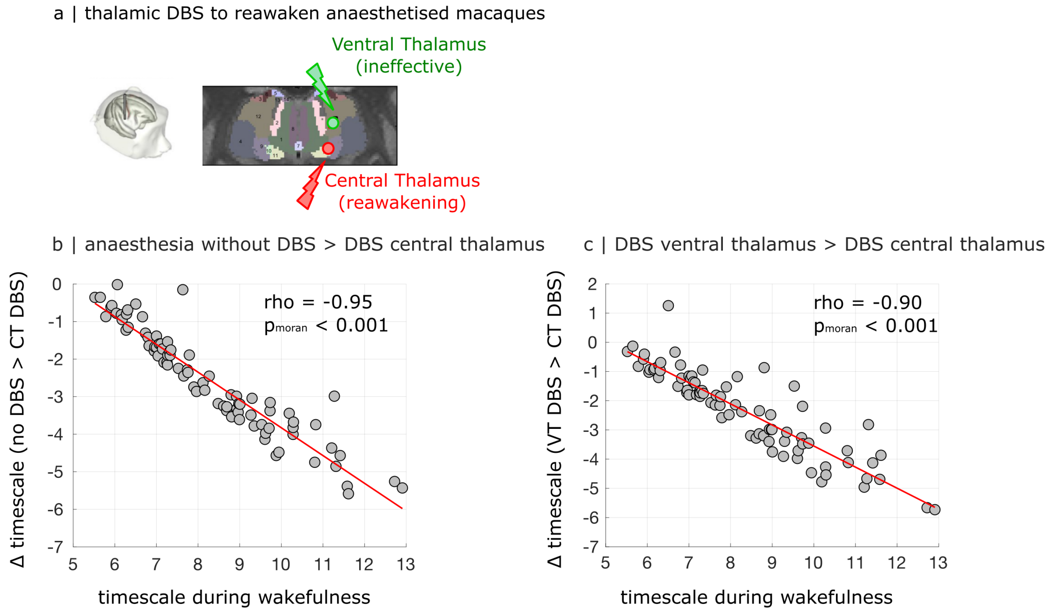


Figure S8. **DBS-induced restoration of intrinsic neural timescales in the anaesthetised macaque |** (**a**) Illustration of deep-brain stimulation targets in the macaque thalamus: centromedian thalamus, and ventral thalamus (adapted from Figure 1 of [43], published under CC-BY license). (**b**) Regional intrinsic neural timescales are increased during reawakening from anaesthesia induced by CT DBS, than under propofol anaesthesia without DBS, and the change is proportional to the value at wakefulness. (**c**) Regional intrinsic neural timescales are increased during reawakening from anaesthesia induced by CT DBS, than during VT DBS at the same intensity (which does not reawaken the animal), and the change is proportional to the value at wakefulness. Both effects resemble the difference between anaesthesia and baseline wakefulness, even though here wakefulness is achieved through stimulation while propofol is still present.

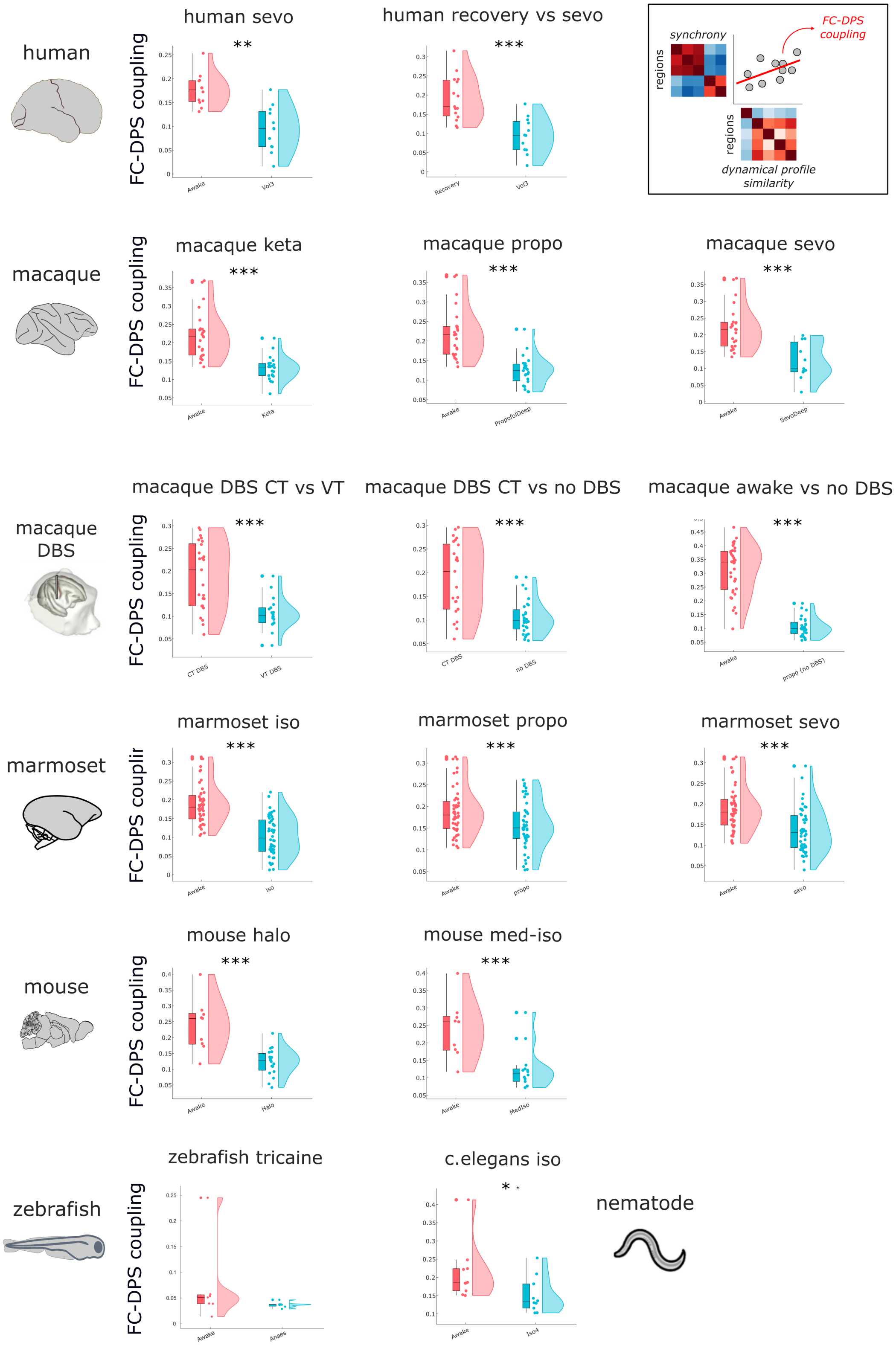


Figure S9. **Anaesthetic effect on the coupling between synchrony and dynamical profile similarity |** Ordinate represents the correlation between inter-regional synchrony and dynamical profile similarity. Each data-point represents one scan. Box-plots: center line, median; box limits, upper and lower quartiles; whiskers, $1.5\times$ interquartile range. *, $p < 0.05$; **, $p < 0.01$; ***, $p < 0.001$, from non-parametric permutation-based t-test (dependent samples for human, marmoset and zebrafish; independent samples for macaque, mouse and nematode).

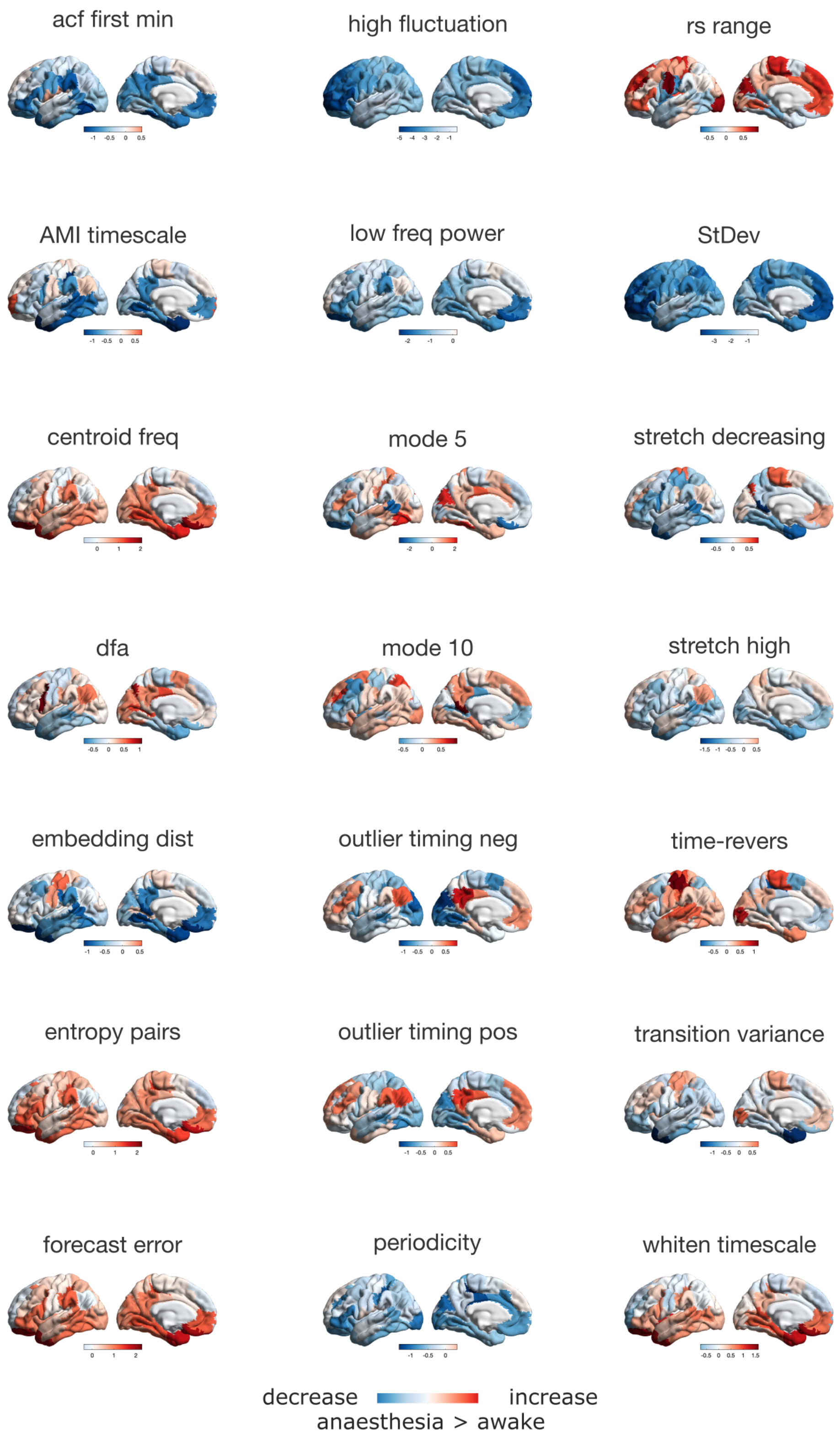


Figure S10. **Regional anaesthetic-induced changes in dynamical features for the** `catch22` **set, for the human brain |** For each region, effect sizes are averaged across all human anaesthesia > awake contrasts.

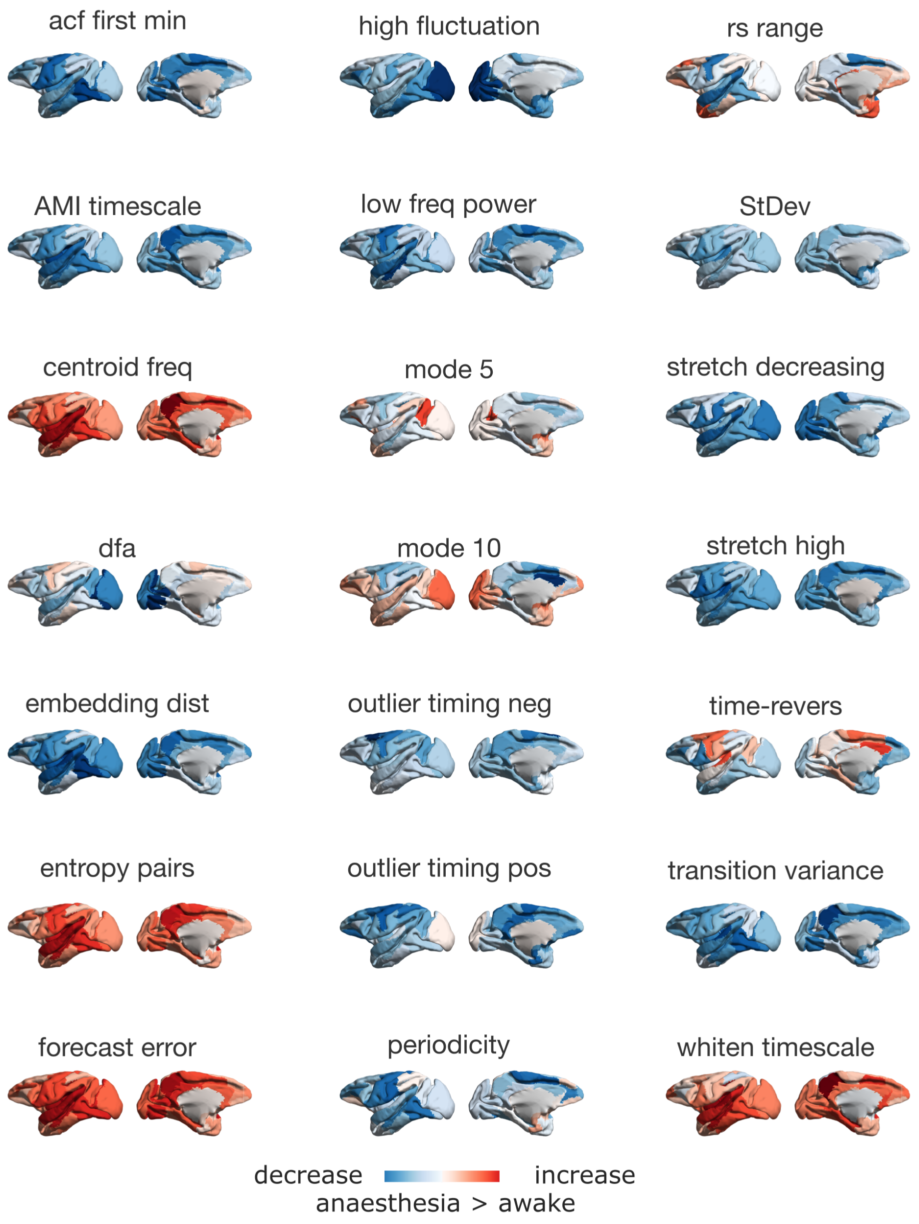


Figure S11. **Regional anaesthetic-induced changes in dynamical features for the** `catch22` **set, for the macaque brain |** For each region, effect sizes are averaged across all macaque anaesthesia > awake contrasts.

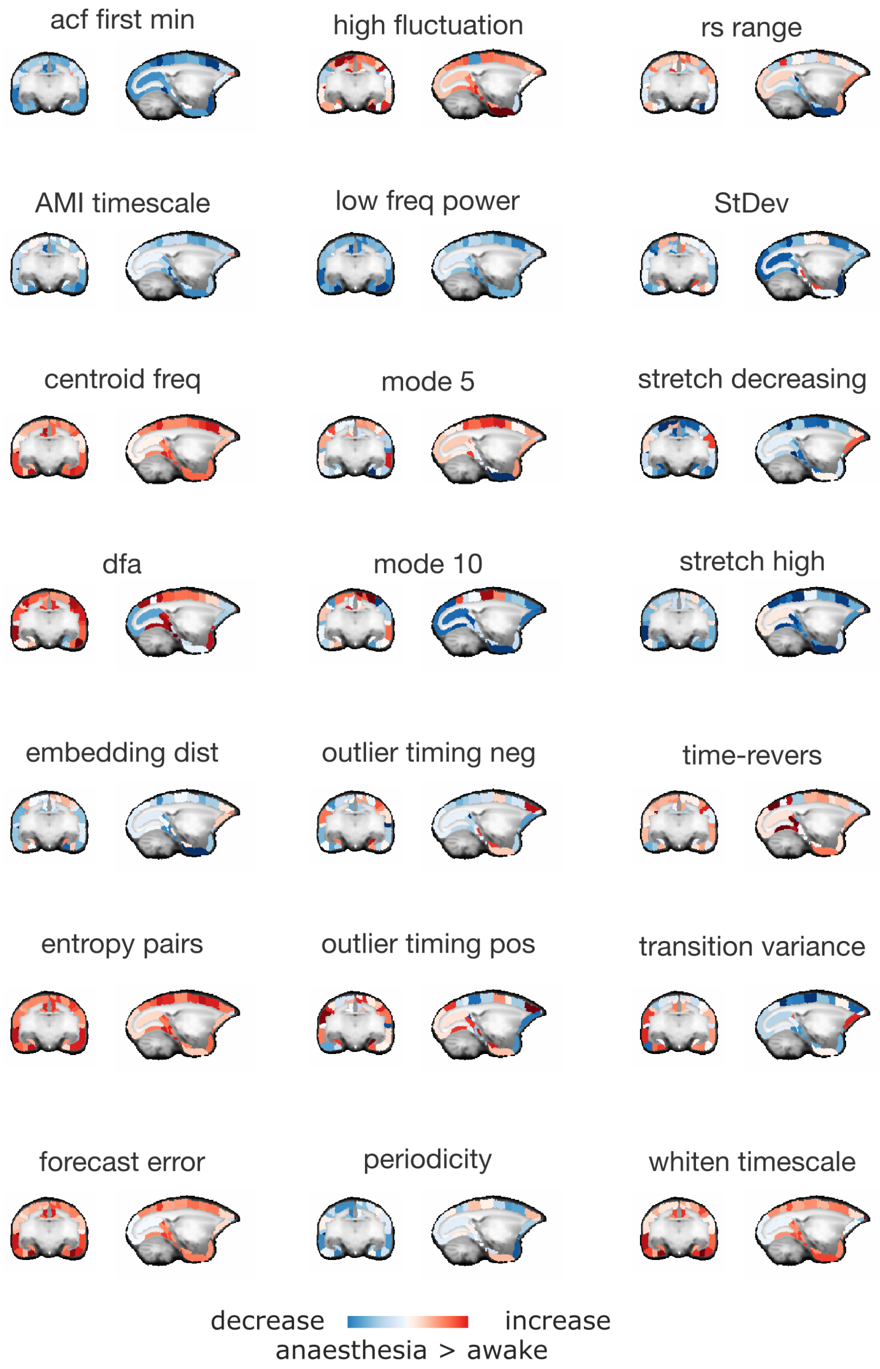


Figure S12. **Regional anaesthetic-induced changes in dynamical features for the `catch22` set, for the marmoset brain |** For each region, effect sizes are ma across all marmoset anaesthesia > awake contrasts.

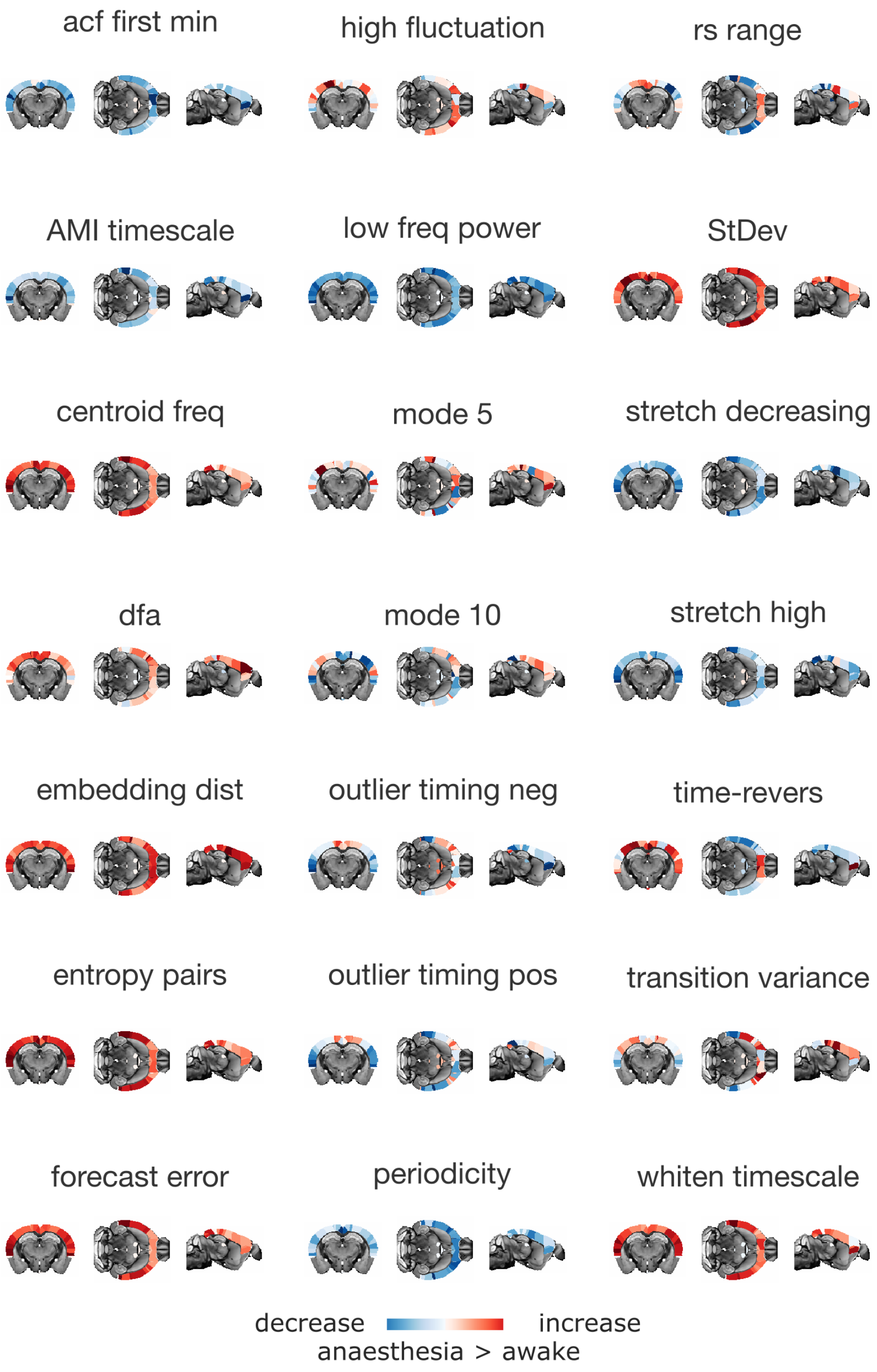


Figure S13. **Regional anaesthetic-induced changes in dynamical features for the `catch22` set, for the mouse brain |** For each region, effect sizes are ma across all mouse anaesthesia > awake contrasts.

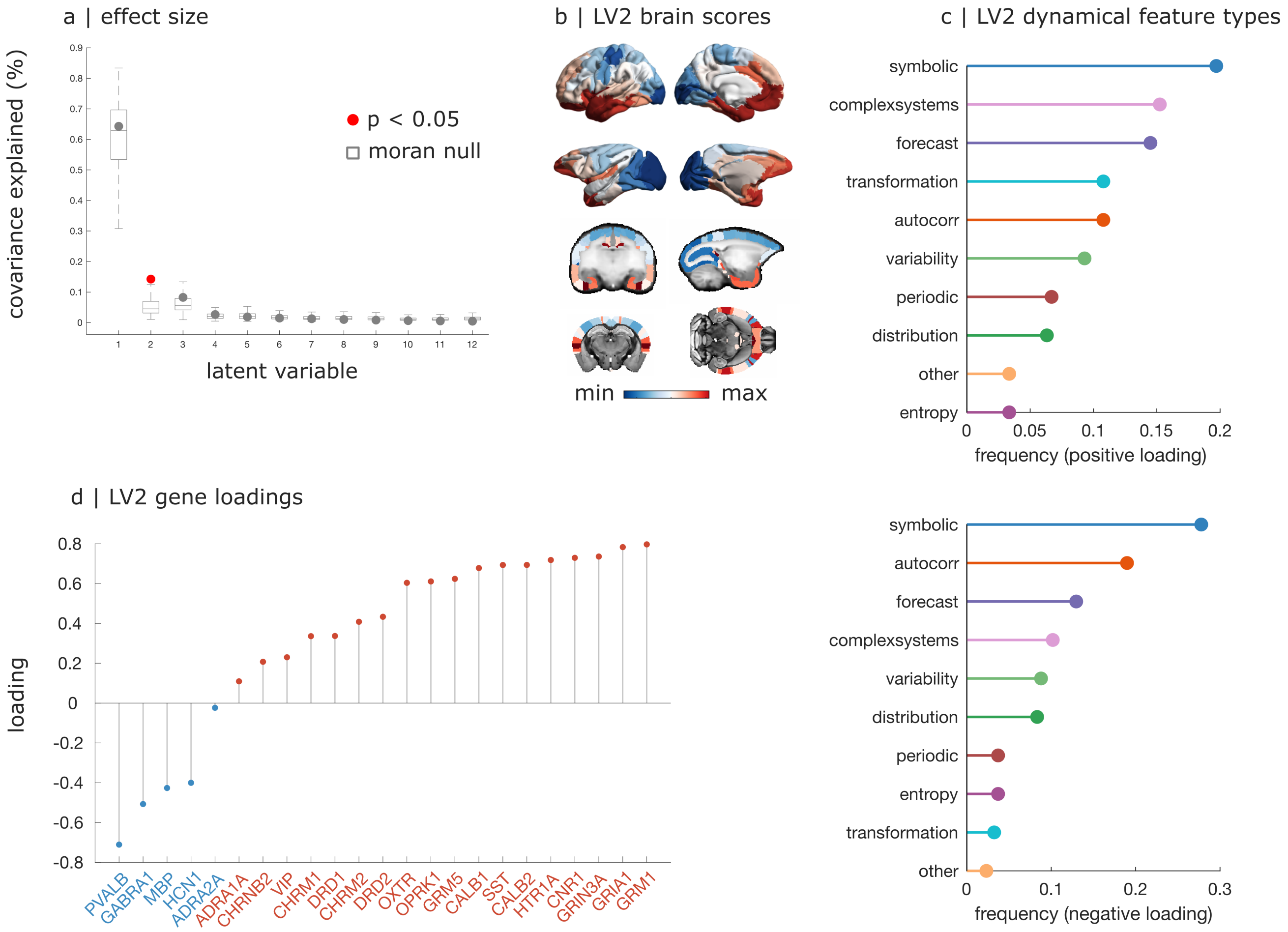


Figure S14. **Gene-dynamics association across the full set of dynamical features that are consistently perturbed by anaesthesia |** (**a**) We find a statistically significant latent dimension of multivariate association between gene expression and anaesthetic-induced feature change (LV2). The first latent dimension is not significant beyond the effect of spatial autocorrelation. (**b**) Representation of the significant LV2 on the cortex of each species, delineating an evolutionarily conserved posterior-dorsal to anterior-ventral gradient. (**c**) Proportion of features belonging to each of 10 broad categories, among those with positive LV2 loading (top), and among those with negative LV2 loading (bottom). (**d**) Loading of each gene onto the significant LV2 dimension of multivariate association with anaesthetic-induced changes in local dynamical features. Red indicates positive loading, blue indicates negative loading.

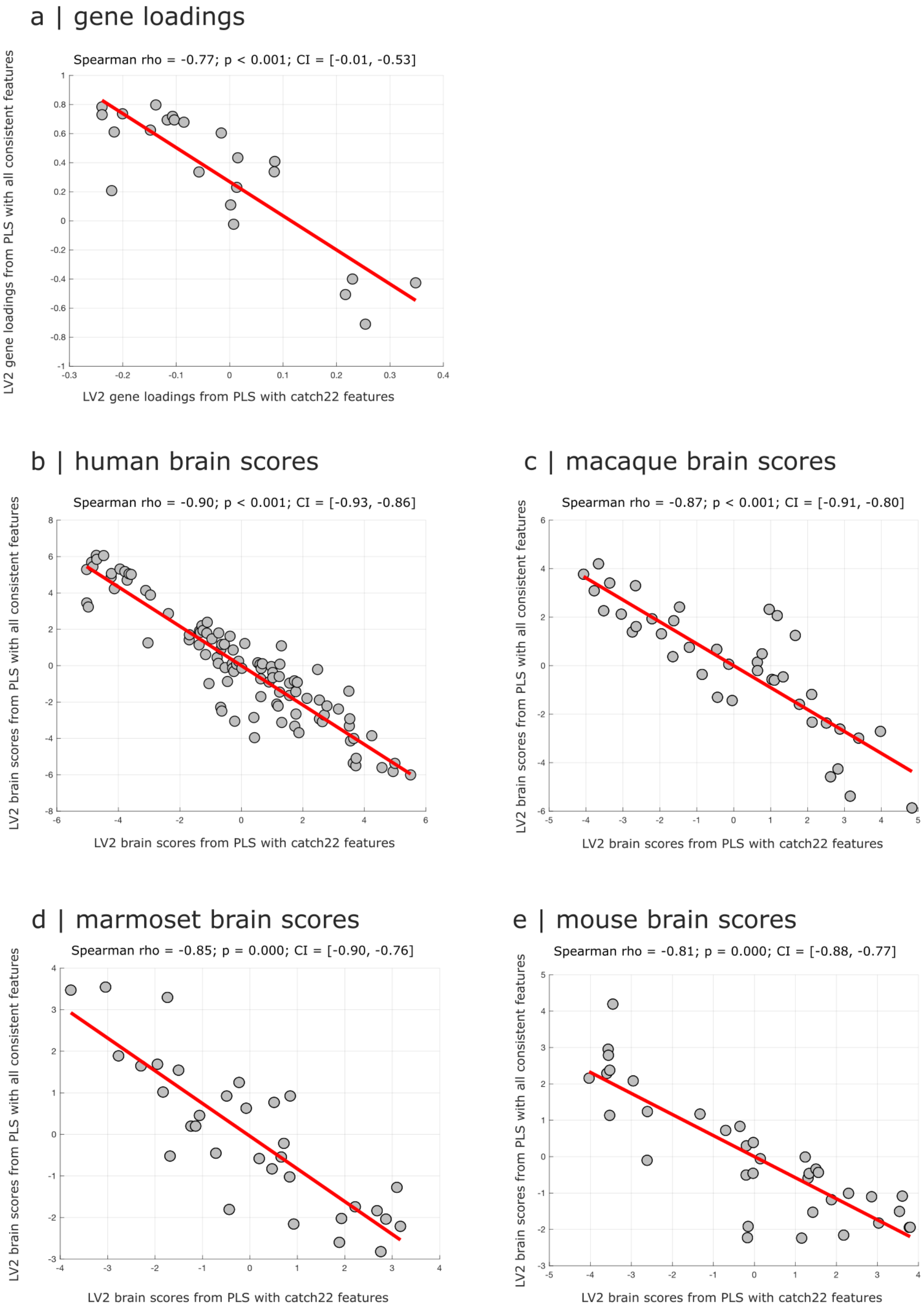


Figure S15. **Replicating conserved significant dimensions of gene-dynamics association with all dynamical features that are consistently perturbed by anaesthesia |** (**a**) There is a significant correlation between LV2 gene loadings obtained with the `catch22` subset of dynamical features, and obtained with the full set of 485 consistent features from `hctsa`. Note that sign is arbitrary with PLS so what matters is only the magnitude of correlation. (**b-e**) Significant correlations between LV2 brain scores obtained with the `catch22` subset of dynamical features (Fig. 6c), and with the full set of 485 consistent features from `hctsa` for human (b), macaque (c) marmoset (d) and mouse (e). Note that sign is arbitrary with PLS so what matters is only the magnitude of correlation.

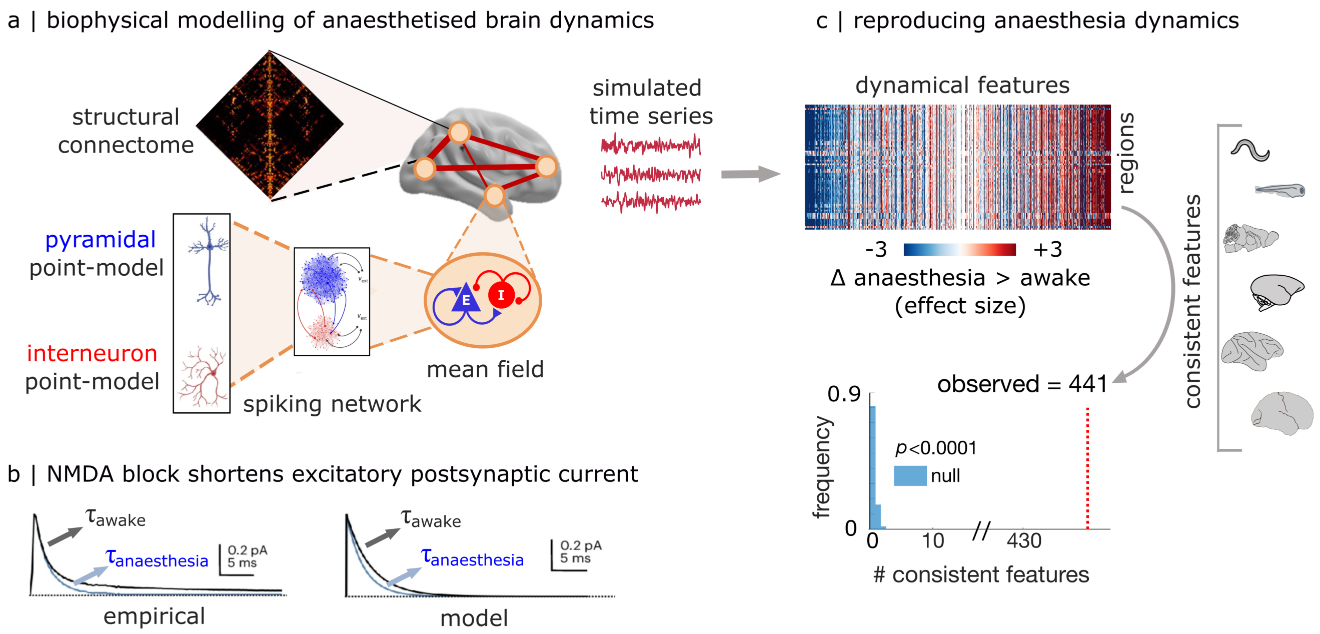


Figure S16. **Computational modelling of ketamine-induced changes in macroscale local dynamics from microscale synaptic mechanisms |** (**a**) Computational model to assess how microscale molecular mechanisms impact macroscale brain dynamics. The whole-brain model consists of biophysically grounded mean-field reductions (one for each brain region) of networks of excitatory and inhibitory neurons, integrating microscale cellular and molecular properties (membrane conductances and AMPA, NMDA, and $GABA_A$ synaptic receptors) to simulate regional fMRI signals. Once connected according to empirical anatomical connectivity of the human brain from in vivo diffusion tractography, the resulting whole-brain model links biophysically meaningful synaptic-level mechanisms to macroscale dynamics. (**b**) At the microscale, the model simulates the empirically-observed shortening of excitatory post-synaptic currents induced by NMDA receptor antagonists such as ketamine, as a decrease in the time constant of excitatory synaptic decay. (**c**) At the macroscale, we apply comprehensive dynamical phenotyping to the simulated BOLD signals from the model in the awake and anaesthetised regimes (N=19 simulations for each condition), following the same workflow as for our empirical data. Among the 485 features that had exhibited consistent changes across all empirical contrasts from Fig. 3, 441 features (>90%) also exhibit changes in the same direction in the computational model. This number is significantly greater than expected by chance, as assessed by a null model where each feature's direction of change (increase or decrease) is assigned at random, repeated 10 000 times ($p < 0.0001$).

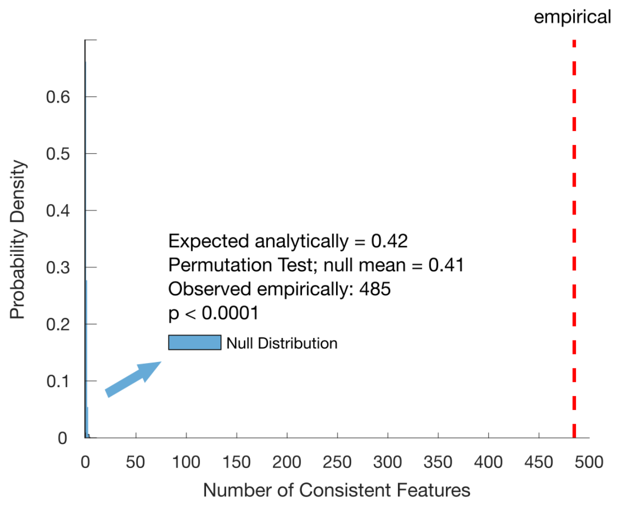


Figure S17. **The empirically observed number of consistent dynamical features is significantly greater than the number expected by chance alone |** We construct a null distribution whereby the sign of each feature and contrast is assigned at random (+1 or -1). Repeating this process 10,000 times produces on average fewer than 1 feature exhibiting consistent sign across all 15 contrasts, whereas we observe 485 ($p < 0.0001$).

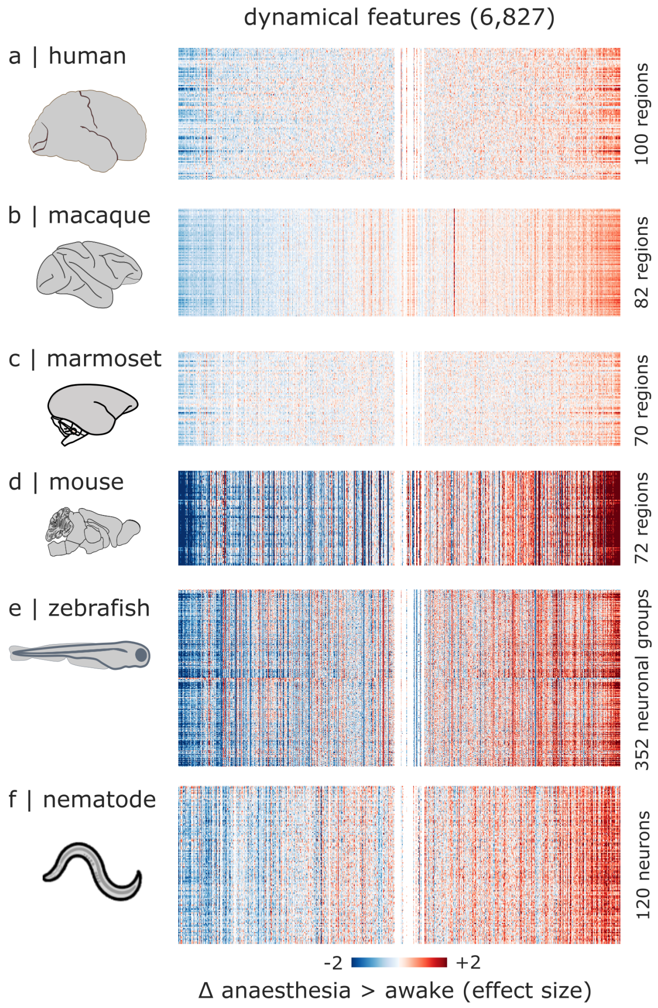


Figure S18. **Anaesthetic-induced changes in neural dynamics across species, after harmonising all datasets by downsampling to TR = 2.4s |** (**a**) Human: mean effect sizes across awake vs vol 3% sevoflurane, and recovery vs vol 3% sevoflurane. (**b**) Macaque: mean effect sizes across awake vs sevoflurane; awake vs propofol; awake vs ketamine (for the Multi-anaesthesia dataset); awake vs propofol (no DBS); CT DBS versus propofol; and CT DBS vs VT DBS. (**c**) Marmoset: mean effect sizes across awake vs sevoflurane; awake vs propofol; and awake vs isoflurane. (**d**) Mouse: mean effect sizes across awake vs halothane; and awake vs medetomidine-isoflurane. (**e**) Larval zebrafish: effect size for awake vs tricaine. (**f**) Nematode: effect size for awake vs isoflurane. The order of features (columns) is the same in each species, sorted to highlight common patterns. For visualisation purposes, the color range is capped at [-2, 2].

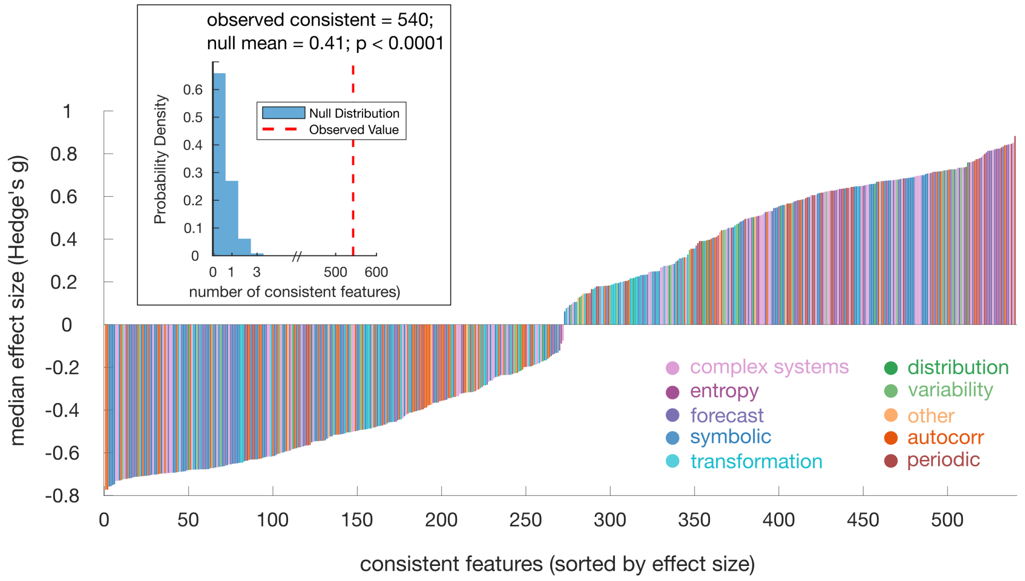


Figure S19. **Consistent effects of anaesthesia on neural dynamics after harmonising all datasets by downsampling to TR = 2.4s |** We find 540 features that change consistently across all contrasts between wakefulness and anaesthesia. The median effect sizes (Hedge's *g*) are shown for each of the consistent features, color-coded according to membership of 10 broad categories of dynamics (feature types; see *Methods* and Table S2). (**Inset**) The empirically observed number of consistent dynamical features is significantly greater than the number that would be expected to exhibit consistency by chance alone (approximately 1 for every 16,000). We construct a null distribution whereby the sign of each feature and contrast is assigned at random (+1 or -1). Repeating this process 10,000 times produces on average fewer than 1 feature exhibiting consistent sign across all 15 contrasts, whereas we observe 540 ($p < 0.0001$). Note that x-axis is interrupted to better show the null distribution.

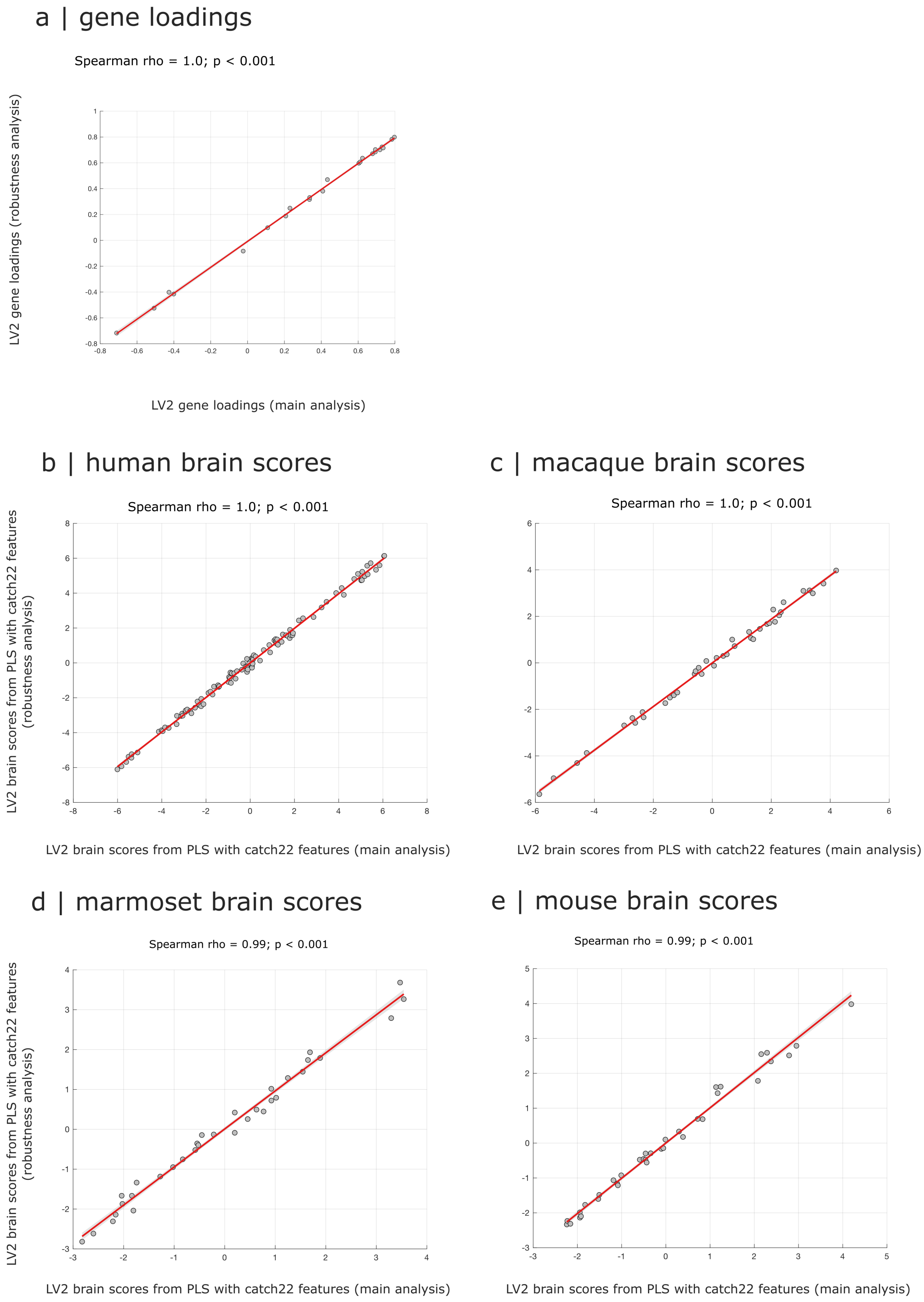


Figure S20. **Replicating conserved significant dimensions of gene-dynamics association with additional contrasts |** We repeat our PLS analysis after including additional contrasts for the human sevoflurane dataset (2% vol and burst-suppression), showing that results remain consistent, with correlated LV2 gene loadings (a), and correlated LV2 brain scores for human, macaque, marmoset, and mouse (b-e).

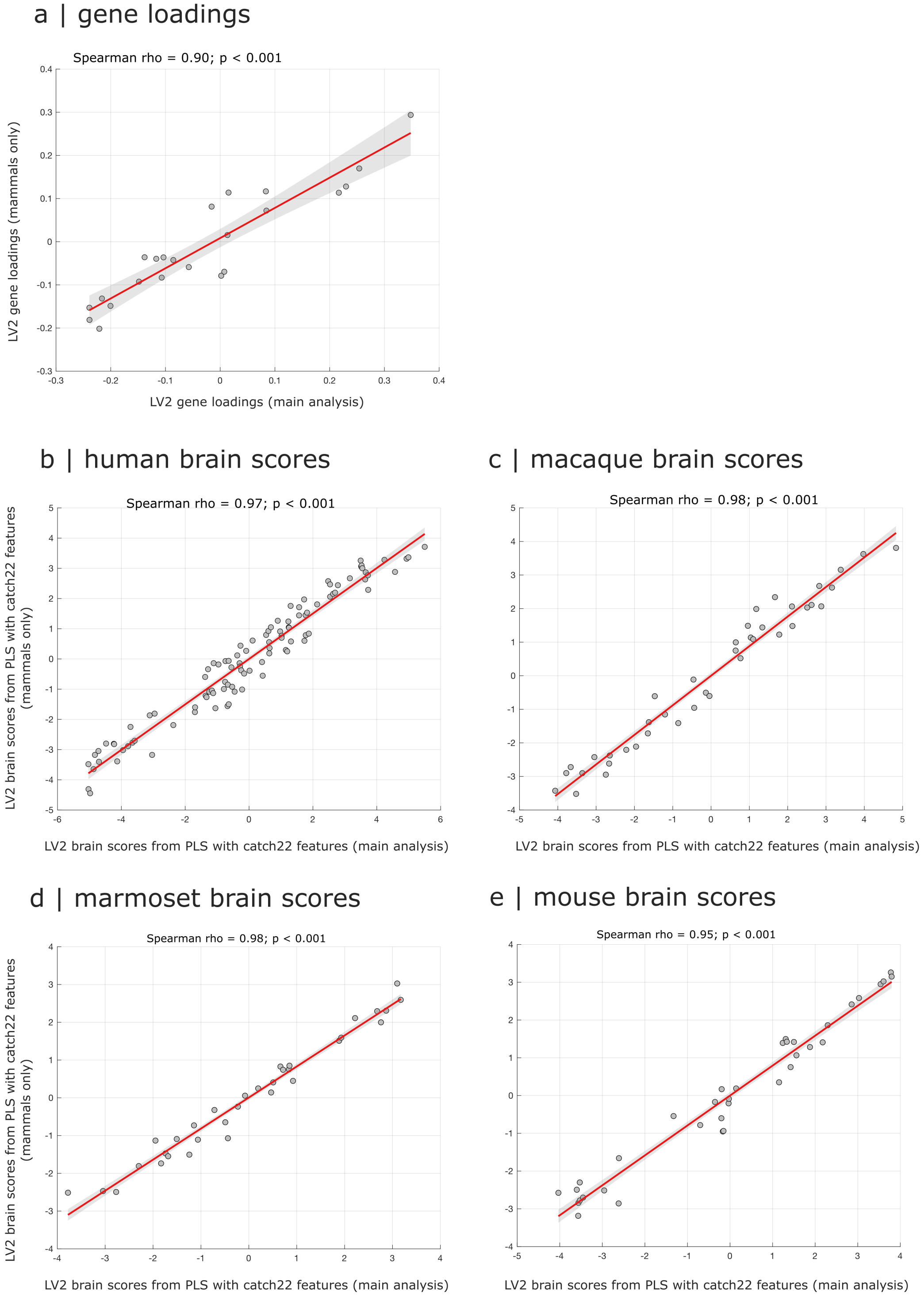


Figure S21. **Replicating conserved significant dimensions of gene-dynamics association when only including data from mammalian species (human, macaque, marmoset, mouse) for the identification of consistent features |** We repeat our PLS analysis after excluding the nematode and zebrafish calcium imaing data, showing that results remain consistent, with correlated LV2 gene loadings (a), and correlated LV2 brain scores for human, macaque, marmoset, and mouse (b-e).

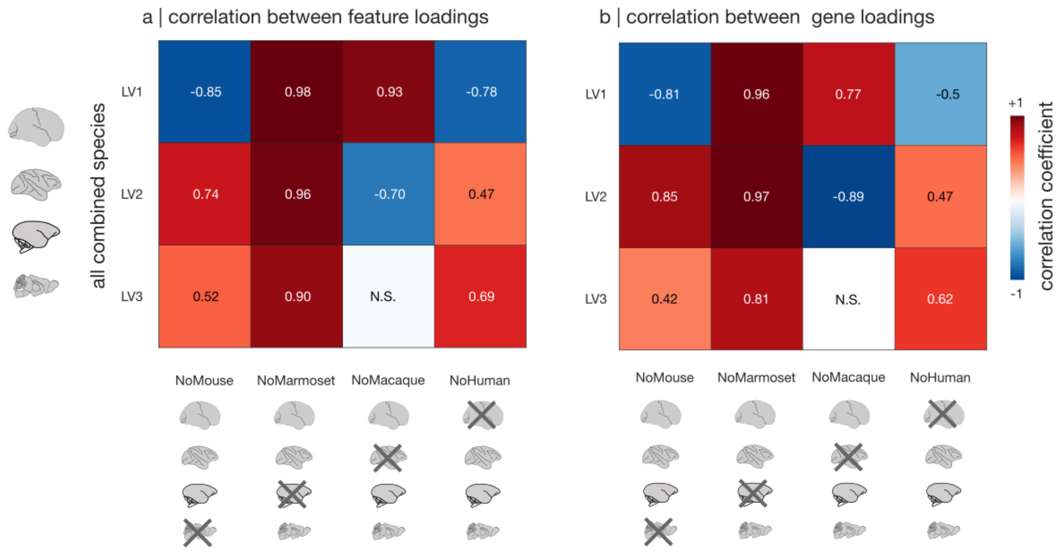


Figure S22. **PLS results are not critically dependent on any individual species included |** We perform a leave-one-species-out cross-validation analysis, whereby we systematically repeat the genes-features PLS analysis after excluding one species (human, macaque, mouse, or marmoset) each time. (**a**) Correlations between dynamical feature loadings for each of the first 3 latent variables, between the version with all four species, and each of the versions with one species excluded. (**b**) Correlations between gene loadings for each of the first 3 latent variables, between the version with all four species, and each of the versions with one species excluded. N.S., not significant. Note that PLS sign is arbitrary.

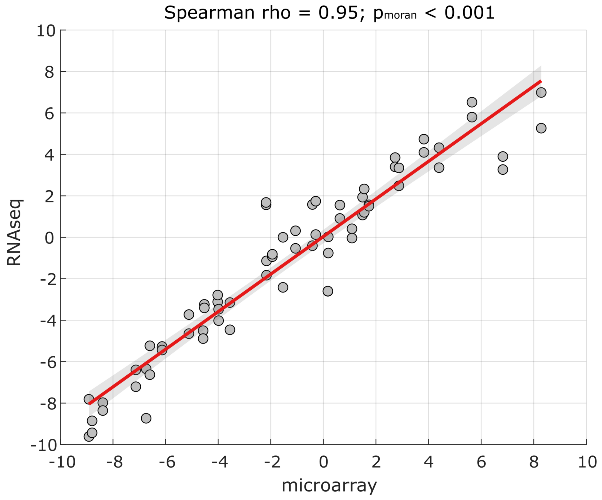


Figure S23. **Replicating human significant PLS brain scores with RNA-seq gene expression |** Abscissa: human LV2 brain scores from microarray gene expression. Ordinate: human LV2 brain scores from RNA-seq gene expression. Each data-point is one cortical region.

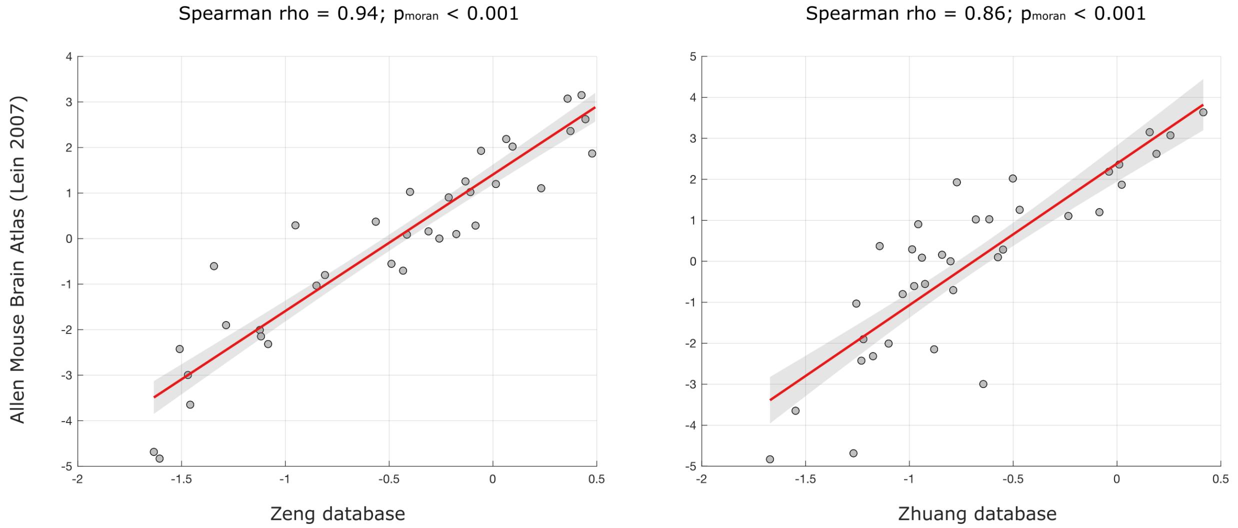


Figure S24. **Replicating mouse significant PLS brain scores with alternative databases of mouse gene expression |** Left: replication with the database of [80]. Right: replication with the database of [81]. Axes reflect LV2 brain scores for the mouse. Each data-point is one cortical region of the mouse brain.

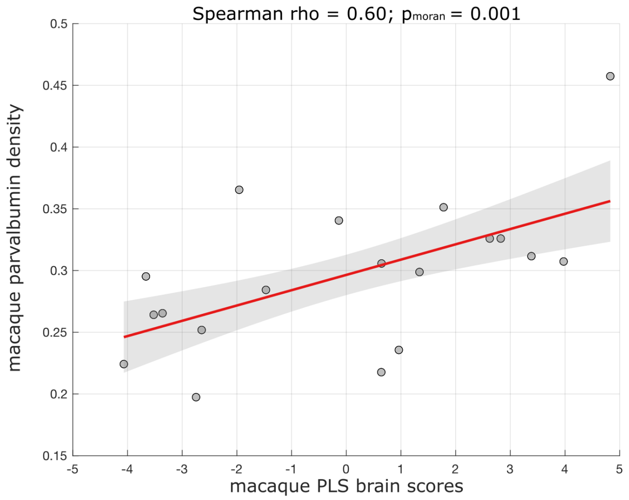


Figure S25. **Macaque significant PLS brain scores are significantly associated with regional parvalbumin protein density |** Abscissa: macaque LV2 brain scores. Ordinate: macaque regional parvalbumin protein density from immunohistochemistry [82]. Each data-point is one cortical region of the macaque brain for which data are available in both modalities.

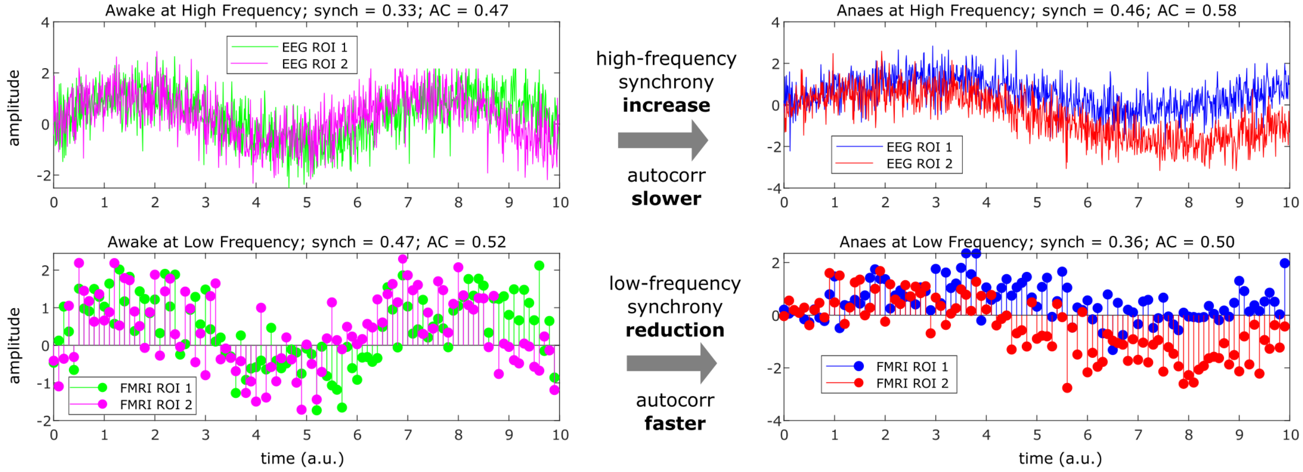


Figure S26. **Toy example showing that sampling rate influences estimated changes in synchrony and autocorrelation in the same time-series |** Left: the same pair of synthetic time-series are shown, sampled at fast rate (top) or slow rate (100x slower; bottom). Right: a different pair of synthetic time-series, also sampled at fast or slow rate. When considering the fast (EEG-like) sampling, the time-series on the right appear more synchronous and more autocorrelated than those on the left, coinciding with the effects of anaesthesia on EEG signals described in the literature. However, when considering the slower (fMRI-like) sampling, the time-series on the right appear *less* synchronous and *less* autocorrelated than those on the left, matching the results from the present report. This toy example shows that observing lower autocorrelation and reduced synchrony at slow timescales is compatible with higher autocorrlation and more synchrony at fast timescales, not only in theory but also in practice.

**Supplementary Tables**

| Species | Awake conditions | Anaesthesia conditions | Imaging modality | Temporal resolution | Time points | Spatial coverage | Reference |
|---|---|---|---|---|---|---|---|
| Human (*Homo sapiens*) N=15 | -awake (15 scans) -recovered (15 scans) | sevoflurane (15 scans) | fMRI 3T BOLD | 1838 ms | 346 | 100 | Ranft et al., (2016) *Anesthesiology* |
| Macaque (*Macaca mulatta*) N=5 | awake (24 scans) | -sevoflurane (11 scans) -propofol (23 scans) -ketamine (22 scans) | fMRI 3T MION | 2400 ms | 500 | 82 | Uhrig et al., (2018) *Anesthesiology* |
| Macaque (*Macaca mulatta*) N=5 | -awake (36 scans) -reawakened by central thalmus DBS (25 scans) | -propofol (28 scans) -propofol with ventral thalamus DBS (18 scans) | fMRI 3T BOLD | 1250 ms | 500 | 82 | Tasserie et al., (2022) *Science Advances* |
| Marmoset (*Callithrix jacchus*) N=4 | awake (48 scans) | -propofol (48 scans) -sevoflurane (48 scans) -isoflurane (48 scans) | fMRI 9.4T BOLD | 2000 ms | 150 | 104 | Muta et al., (2023) *Cerebral Cortex* |
| Mouse (*Mus musculus*) N=43 | awake (10 scans) | -med-iso (14 scans) -halothane (19 scans) | fMRI 7T BOLD | 1000 ms (1200 ms halo) | 1414 | 72 | Gutierrez-Barragan et al., (2023) *Current Biology* |
| Nematode (*C. elegans*) N=10 | 0% iso (10 scans) | 4% isoflurane (10 scans) | GCaMP6s calcium imaging | 500 ms | 500 | 120 | Awal et al., (2020) *Anesthesiology* |
| Larval zebrafish (*Danio rerio*) N=7 | awake (7 scans) | tricaine (7 scans) | GCaMP calcium imaging | 1014 ms | 250 | 352 | This article |

TABLE S1. **Overview of the seven datasets included in the present study, and their key parameters.** For human, macaque, marmoset, and mouse, 'Spatial coverage' indicates the number of cortical regions. For the nematode and zebrafish, it indicates the number of neurons or brain regions (groups of neurons). MION, monocrystalline iron oxide nanoparticle contrast agent.

| Category | Example |
|---|---|
| **Distribution** | Mean, variance, outliers, tests for distribution family |
| **Autocorrelation** | Nonlinear autocorrelation, lagged autocorrelation |
| **Periodic** | Power spectrum, seasonality, wavelet |
| **Entropy** | Entropy measures and related quantities |
| **Symbolic** | Discretisation of the signal, motifs, transitions between motifs |
| **Forecasting** | Forecasting from past to future, model fit measures |
| **Transformations** | Scaling, surrogates |
| **Variability in Time** | Change-points, stationarity |
| **Complex Systems Measures** | Time-reversal, fractality, embeddings |
| **Miscellaneous others** | Visibility graph |

TABLE S2. **Broad categories of time-series features**.

| Feature name | Category | Description |
|---|---|---|
| mode_5 | Distribution | 5-bin histogram mode |
| mode_10 | Distribution | 10-bin histogram mode |
| outlier_timing_pos | Distribution | Timing of positive extreme event |
| outlier_timing_neg | Distribution | Timing of negative extreme event |
| acf_timescale | Autocorrelation | First $1/e$ crossing of the ACF |
| acf_first_min | Autocorrelation | First minimum of the ACF |
| low_freq_power | Periodic | Power in the lowest 20% frequencies |
| centroid_freq | Periodic | Centroid frequency |
| forecast_error | Forecasting | Error of 3-point rolling mean forecast |
| whiten_timescale | Forecasting | Change in autocorrelation timescale after incremental differencing |
| high fluctuation | Variability | Proportion of high incremental changes in the series |
| stretch_high | Symbolic | Longest stretch of above-mean values |
| stretch_decreasing | Symbolic | Longest stretch of decreasing values |
| entropy_pairs | Entropy | Entropy of successive two-symbol motifs in the symbolized series |
| ami2 | Autocorrelation | Histogram-based automutual information (lag 2, 5 bins) |
| time_revers | ComplexSystems | Time reversibility |
| ami_timescale | Autocorrelation | First minimum of the AMI function |
| transition_variance | Symbolic | Transition matrix column variance |
| periodicity | Periodic | Wang's periodicity metric |
| embedding_dist | ComplexSystems | Goodness of exponential fit to embedding distance distribution |
| rs_range | Transformations | Rescaled range fluctuation analysis (low-scale scaling) |
| dfa | Transformations | Detrended fluctuation analysis (low-scale scaling) |
| standard_deviation | Distribution | Standard deviation of the distribution of time-series values |

TABLE S3. Representative subset of dynamical features known as the CAnonical Time-series CHaracteristics (*catch22*) from Lubba et al. [67], along with broad category assignment.

| Gene | Full name |
|---|---|
| ADRA1A | Adrenergic receptor, $\alpha_{1A}$ |
| ADRA2A | Adrenergic receptor, $\alpha_{2A}$ |
| CALB1 | Calbindin 1 |
| CALB2 | Calbindin 2 (calretinin) |
| CHRM1 | Cholinergic receptor, muscarinic 1 |
| CHRM2 | Cholinergic receptor, muscarinic 2 |
| CHRNB2 | Nicotinic acetylcholine receptor subunit $\beta_2$ |
| CNR1 | Cannabinoid receptor 1 |
| DRD1 | Dopamine receptor D1 |
| DRD2 | Dopamine receptor D2 |
| GABRA1 | Gamma-aminobutyric acid receptor subunit $\alpha_1$ |
| GRIA1 | Glutamate receptor, ionotropic, AMPA 1 |
| GRIN3A | Glutamate receptor, ionotropic, NMDA 3A |
| GRM1 | Glutamate receptor, metabotropic 1 |
| GRM5 | Glutamate receptor, metabotropic 5 |
| HCN1 | Hyperpolarization-activated cyclic nucleotide-gated channel 1 |
| HTR1A | Serotonin receptor 1A |
| MBP | Myelin basic protein |
| OPRK1 | Opioid receptor, kappa 1 |
| OXTR | Oxytocin receptor |
| PVALB | Parvalbumin |
| VIP | Vasoactive intestinal peptide |
| SST | Somatostatin |

TABLE S4. List of brain-related genes included in the present study, which are available across cortical regions for human, macaque, mouse, and marmoset.

| Parameter | Score Range | Description |
|---|---|---|
| Exploration of the surrounding world | 0 to 2 | • 0 = Total absence<br>• 1 = Small search of external clues<br>• 2 = Total investigation of the environment (e.g., head orientation to a sound) |
| Spontaneous movements | 0 to 2 | • 0 = Total absence<br>• 1 = Small torso and/or limb movement<br>• 2 = Large torso and/or limb movement |
| Shaking / prodding | 0 to 2 | • 0 = Total absence<br>• 1 = Small body movement<br>• 2 = Large body movement |
| Toe pinch | 0 to 2 | • 0 = Total absence<br>• 1 = Small reflex (weak body movement, eye blinking, or cardiac rate change)<br>• 2 = Clear reaction (strong body movement, eye blinking or opening, and cardiac rate change) |
| Eyes opening | 0 to 2 | • 0 = Total absence<br>• 1 = Small blinks or eye movements<br>• 2 = Full eye opening |
| Corneal reflex | 0 to 1 | • 0 = Absent<br>• 1 = Present |

TABLE S5. Scoring criteria for behavioral assessment.
